# Calibrated Variant Effect Prediction at the Residue Level Using Conditional Score Distributions

**DOI:** 10.1101/2025.11.24.690189

**Authors:** Gal Passi, Sapir Amittai, Dina Schneidman-Duhovny

**Affiliations:** The Rachel and Selim Benin School of Computer Science and Engineering, The Hebrew University of Jerusalem, Jerusalem, Israel

## Abstract

Effective clinical use of variant effect prediction (VEP) requires models that are both accurate and well-calibrated. Calibration refers to a model’s ability to produce meaningful and reliable probability estimates. Benchmarking 30 VEPs, we show that while models appear well-calibrated on average, they remain markedly miscalibrated within specific variant subgroups. We propose VEP calibration at the residue level and introduce a calibration approach based on differential score mapping per variant subgroup. When calibration targets are chosen appropriately, our calibration approach can also improve model discrimination. Leveraging these insights, we develop RaCoon (Residue-aware Calibration of Conditional Distributions), implemented on ESM1b, which provides multicalibrated and interpretable predictions. RaCoon introduces Gaussian mixture models-based sampling fitted on model score distribution which substantially reduces labeled-data requirements for calibration. The method maintains low calibration error across variant subgroups and datasets while also improving AUROC on multiple benchmarks. Our calibration strategy is readily transferable to other VEPs.

## Introduction

Accurate variant effect prediction (VEP) is central to protein design, medical genomics, and variant forecasting. Recent advances in deep learning have enabled models that effectively capture evolutionary patterns from protein sequences and structures^1–6^, achieving groundbreaking accuracy in VEP. These models can be broadly categorized into supervised, unsupervised, and weakly supervised approaches.

VEPs are typically evaluated using datasets of clinically reported genetic variants, such as ClinVar^7^, or experimental benchmarks such as ProteinGym, which aggregates data from multiple deep mutational scans (DMS) assays^8^. Two broadly adopted metrics to assess VEP performance are the global AUROC, which measures ranking performance across all proteins, and per-protein AUROC, which captures discriminative ability within individual proteins. State-of-the-art (SOTA) models achieve AUROCs as high as 0.89-0.93 (unsupervised) to 0.98 (supervised) on the ProteinGym Clinical Substitutions subset and on ClinVar-derived benchmarks^9^.

While supervised approaches^10–13^ achieve impressive accuracy in specific proteins or families, they are sensitive to sparse and biased clinical annotations and generalize poorly to unseen protein regions. Consistently, AUROC is strongly driven by protein features^14,15^, using the per-protein label ratio alone on ClinVar was shown to achieve an AUROC of 0.914^6,16,17^. Notably, in ClinVar, only ∼4% of genes have more than five labeled pathogenic and benign variants. These biases also extend to experimental datasets, where VEP performance varies with assay type, protein context, and DMS depth^18,19^.

These limitations have motivated the growing use of unsupervised VEPs, trained without labeled data and therefore less affected by annotation bias. Unsupervised VEPs include alignment-based approaches, trained on multiple sequence alignments (MSAs)^1,20^, protein language models (PLMs) trained on unaligned protein sequences^4,21–24^, and hybrid approaches^4,21–23,25^. While alignment-based approaches perform well, their inference is limited in regions with shallow alignments, such as intrinsically disordered regions (IDRs) or proteins with few homologs. PLMs provide an alternative approach that does not rely directly on explicit alignments, and can offer more consistent inference in such regions, although their ability to generalize beyond MSAs remains an open question^26^. Hybrid retrieval-augmented models can further improve performance by selectively incorporating MSA information.

While not exposed to labeled data, unsupervised VEPs still embed assumptions that fail in certain variant subgroups^18,27,28^. To address these biases, recent works employ forms of weak supervision, calibrating scores to specific datasets^6,29,30^, proteins^1,31,32^, diseases^33^ or organisms^34^ at inference time using a subset of labeled data from the relevant domain.

Classifier calibration is defined as a model’s ability to provide meaningful probability estimates^35^. For example, an assigned score of 0.8 should correspond to ∼80% of similar variants being pathogenic. However, large-scale models often exhibit poor calibration^36^, and performance frequently degrades on out-of-domain data^37^, such as VEPs evaluated on test sets not overlapping with ClinVar^9^. Calibration is not always correlated with accuracy^38^ and oftentimes, training objectives intended to boost accuracy can reduce calibration^39^.

Several examples motivate examining VEP calibration within variant subgroups, notably variants in IDRs. IDRs lack stable structure, evolve rapidly, and are more tolerant to mutations^40–43^. Although IDRs are estimated to comprise roughly 26-35% of the human proteome^44,45^, only 12-15% of ClinVar pathogenic variants occur in these regions, with over 80% of variants in these regions annotated as benign^42^. In these regions, VEPs tend to favor benign predictions, with high AUROC masking reduced sensitivity to pathogenic variants^14,28,42^. The opposite label composition is observed in N-terminal methionine variants, where ∼95% are pathogenic in ClinVar. For these variants, VEPs exhibit pronounced, model-specific imbalances between sensitivity and specificity^42^. Other examples include variants with distinct physico-chemical properties or few homologs^29,46^. Together, these examples highlight differences in model behavior across residue subgroups which may be obscured by pooled evaluations.

As VEPs reach unprecedented accuracy^15^ and wider clinical use, identifying and calibrating for distribution shifts has gained increasing attention^29,47–49^ While models appear globally calibrated, they may be systematically miscalibrated for specific subgroups, posing challenges for clinical decision making^50^. A multicalibrated model, whose predicted probabilities remain accurate not only overall but also within each relevant subgroup of variants^51^, would therefore be highly desirable for VEPs. Recent studies suggest that multicalibration can often be achieved with minimal post-hoc adjustment^52^ and typically does not compromise, and may even improve, discriminatory performance^51,53^.

Here, we show that current calibration approaches lose reliability when examined at the residue level. We propose a weakly supervised framework to correct residue-level miscalibration and demonstrate that model-specific score distributions can identify optimal calibration targets. This approach not only closes existing reliability gaps but, for most models, also yields modest improvements in AUROC. Finally, we introduce RaCoon (Residue-aware Calibration of Conditional Distributions), a calibrated PLM based on ESM1b, capable of producing interpretable and well-calibrated pathogenicity predictions across residue subgroups while improving the baseline model’s AUROC^4^.

## Results

### Benchmarking 30 VEPs reveals profound residue-level miscalibration

To benchmark calibration across 30 SOTA VEPs (**Table 1**), we analyze model behavior in biologically relevant residue contexts, such as disordered regions and protein interfaces. We examine two calibration metrics: First, we assess whether classification thresholds and model performance vary across residue-defined subgroups. Second, we assess reliability directly by asking whether predicted probabilities match the observed fraction of pathogenic variants within each subgroup.

**Table 1.** List of VEP benchmarked in the study. HDIV - trained on HumDiv, HVAR- trained on Humvar, noAF and addAF whether allele frequency is used, HGMD - Human Gene Mutation Database, ExAC - Exome Aggregation Consortium, PPI - protein-protein interface. Metapredictors employ multiple classifiers with various features. VARITY (R) and (ER) trained on rare or very-rare variants ^a^ MSA also includes models trained on various homology models, ^b^ also report the scaled PHRED score which we do not use, ^c^ scores scaled around 0, ^d^ also report the scaled EVE-score used in Figure 1c. ^e^ variants not observed in humans defined as pathogenic, common variants defined as benign. ^f^ UniProt polymorphisms, dbSNP, 1000 Genomes, ExAC, and UK10K cohorts. ^g^ A balanced set of ClinVar variants used to stop training during fine-tuning, optimize hyperparameters, and fit a logistic-regression model to the raw model scores. ^h^ VARITY (R) mean allele fraction <0.5% and ER < 10*e* − 6 % ^i^ Two variants in ClinVar_BM and 44 variants in the ProteinGym clinical substitution benchmark (∼0.07%) map to proteins included in CPT-1’s DMS training data.

| Model Name | Supervision Type | Features Used<sup>a</sup> | Train Data (Supervised) | Bounded Scores (0-1) | Reference |
| --- | --- | --- | --- | --- | --- |
| PrimateAI | Unsupervised | MSA, structural features | - | Yes | 54 |
| LIST-S2 | Unsupervised | MSA | - | Yes | 55 |
| PROVEAN | Unsupervised | MSA | - | No | 56 |
| ESM1b | Unsupervised | Sequence only | - | No | 4 |
| SIFT | Unsupervised | MSA | - | Yes | 57 |
| MutationAssessor | Unsupervised | MSA | - | No | 58 |
| ESM1v | Unsupervised | Sequence only | - | No | 21 |
| EVE | Unsupervised | MSA | - | No<sup>d</sup> | 1 |
| GEMME | Unsupervised | MSA | - | No | 20 |
| PoET | Unsupervised | Sequence only | - | No | 2 |
| LRT | Unsupervised | MSA | - | Yes | 59 |
| TranceptEVE | Unsupervised | Protein sequence, MSA | - | No | 3 |
| AlphaMissense | Semi-Supervised | MSA, structural features | ClinVar (Calibration, training-stop<sup>g</sup>), Allele frequency | Yes | 6 |
| CPT-1 | Unsupervised<sup>i</sup> | Protein sequence, MSA, structural features, residue features | DMS data from five human proteins | Yes | 60 |
| FATHMM | Supervised | MSA | HGMD + UniProt/SwissProt | No | 61 |
| DANN | Supervised | MSA, genomic features | Allele frequency<sup>e</sup> (1000 Genomes Project) | Yes | 62 |
| MPC | Supervised | Metapredictor, residue features | Population frequency (ExAC) | No | 63 |
| DEOGEN2 | Supervised | MSA, structural gene, residue and pathway features | Humsavar | Yes | 64 |
| PolyPhen-2 (HDIV) | Supervised | MSA, structural and residue features | HumDiv | Yes | 65 |
| PolyPhen-2 (HVAR) | Supervised | MSA, structural and residue features | HumVar | Yes |  |
| CADD | Supervised | MSA, genomic features | Population frequency<sup>e</sup> (1000 Genomes Project) | No<sup>b</sup> | 66 |
| gMVP | Supervised | MSA, structural and genomic features | ClinVar + Population frequency (gnomAD) | Yes | 67 |
| REVEL | Supervised | Metapredictor | HGMD and Population frequency (1000 Genomes/ESP cohorts) | Yes | 68 |
| BayesDel (noAF) | Supervised | Metapredictor | ClinVar, UniProt Population frequency<sup>f</sup> | No<sup>c</sup> | 69 |
| BayesDel (addAF) | Supervised | Metapredictor | ClinVar, UniProt Population frequency<sup>f</sup> | No<sup>c</sup> |  |
| VEST4 | Supervised | MSA, structural, residue and genomic features | HGMD, ClinVar, Population frequency (Exome Sequencing Project/1000 Genomes) | Yes | 70 |
| ClinPred | Supervised | Metapredictor | ClinVar | Yes | 10 |
| MetaRNN | Supervised | Metapredictor | ClinVar | Yes | 11 |
| VARITY (R) | Supervised | Metapredictor scores, structural and residue features, PPI, protein families and conservation | ClinVar common and rare variants<sup>h</sup> HGMD, HumsaVar, gnomAD, and 12 DMS datasets. | Yes | 71 |
| VARITY (ER) | Supervised |  |  | Yes |  |

Classification thresholds are analyzed using a rigorous ClinVar-derived dataset curated by Radjasandirane et al.^9^ (ClinVar_BM). ClinVar_BM is designed to minimize data circularity and annotation biases, providing a robust test set (Methods, **Table 2**). We compare performance across residue-level features, such as physicochemical and structural properties, as well as protein-level features, such as homology (Methods, **Table 3**), enabling systematic evaluation of subgroup-specific behavior.

**Table 2.** Summary of dataset used in the study.

| Dataset Name | Total variants | Pathogenic variants | Benign variants | Unique sequences | Details | Purpose | Figure Used |
| --- | --- | --- | --- | --- | --- | --- | --- |
| ClinVar_HQ | 171,196 | 52,634 | 118,562 | 14,564 | Curated directly from ClinVar | Primary clinical dataset used for residue-level score-distribution comparisons. | 3b-c, S4-5, S7 |
| ClinVar_BM | 14,053 | 2,296 | 11,757 | 6,599 | variants published in clinvar after may 2021, protein with unbalanced annotations excluded<sup>9</sup> | Standard ClinVar benchmark used for VEPs performance benchmarking - however too small for differential calibration evaluation. | 1a, S1-2 |
| ClinVar_Balanced | 40,474 | 19,526 | 20,948 | 1,337 | Proteins with at least 6 variants and pathogenic to benign ratio between 80:20 and 20:80 | Main clinical evaluation dataset for RaCoon, designed to reduce protein-level label-ratio bias while retaining sufficient data for calibration-tree construction. | 5a-c, S3, S8-9 S11-12 S15a-c, S16-17 |
| Balanced Protein Subset (ClinVar) | 9,354 | 4,390 | 4,964 | 46 | Proteins with at least 100 variants and pathogenic to benign ratio between 70:30 and 30:70 | Stringent clinical evaluation subset used to test RaCoon on well-annotated, label-balanced proteins | S14 |
| ClinVar_AFM | 56,368 | 18,765 | 37,603 | 6,272 | Published by Cheng et al.<sup>6</sup> proteins with no matching transcript excluded | AlphaMissense-specific control dataset used to avoid overlap with variants used for AlphaMissense calibration. | 1c |
| Well-annotated ClinVar proteins | 86,211 | 42,056 | 44,155 | 1,285 | Sequences with at least 10 variants of which at least 4 pathogenic and 4 benign variants | Supplementary protein-level analysis dataset used to compare residue-level and protein-level pathogenic-fraction signals. | S10 |
| ProteinGym | 62,727 | 32,000 | 30,727 | 2,525 | ProteinGym zero-shot clinical substitution benchmark<sup>8</sup> obtained in June 2025 | Primary experimental benchmark used throughout the study for VEPs benchmarking, differential calibration analysis and RaCoon evaluation | 1b, 2a-c, 5a-c, S3-4, S8-9, S11, S15-17 |

**Table 3.**
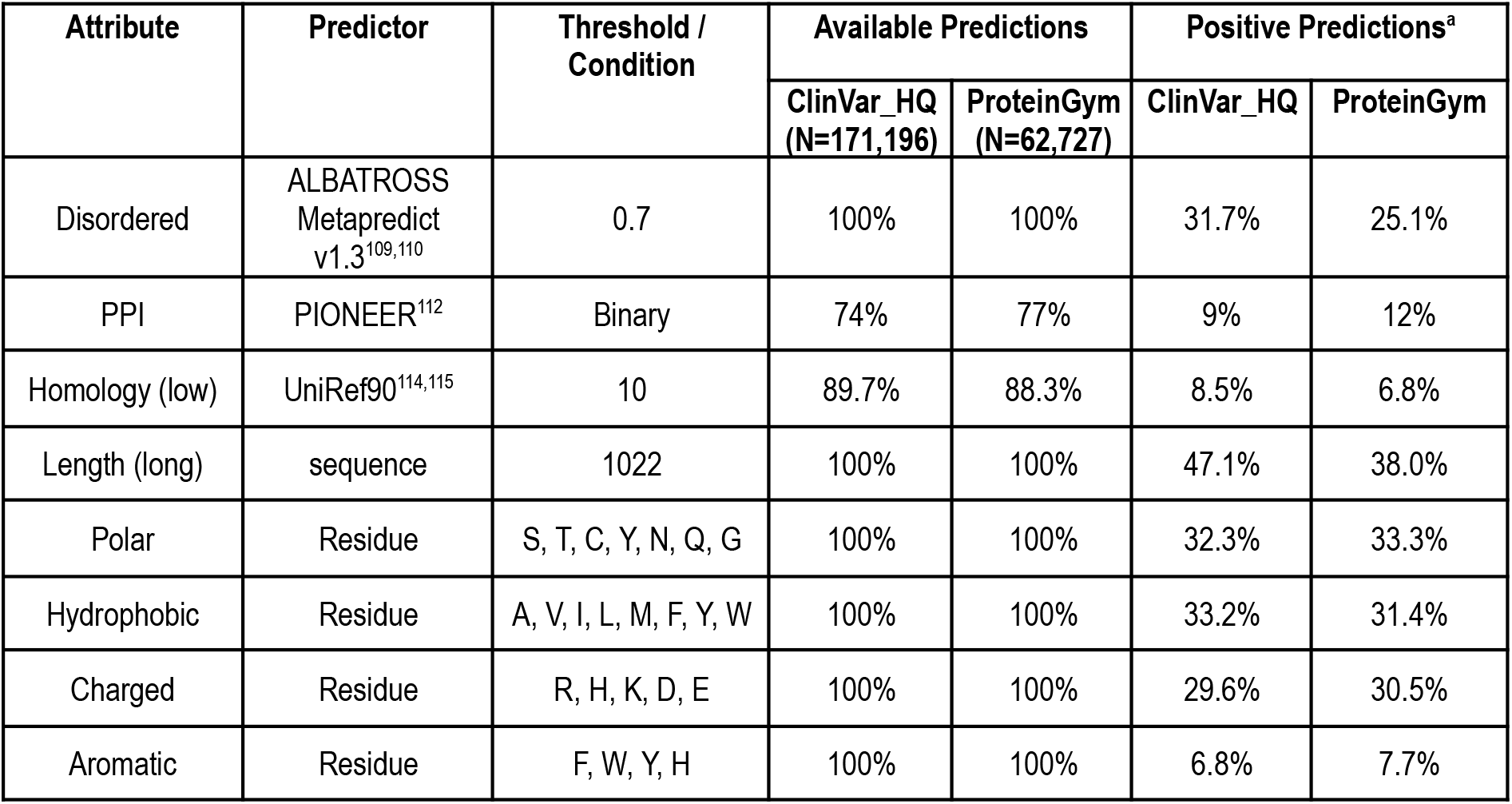
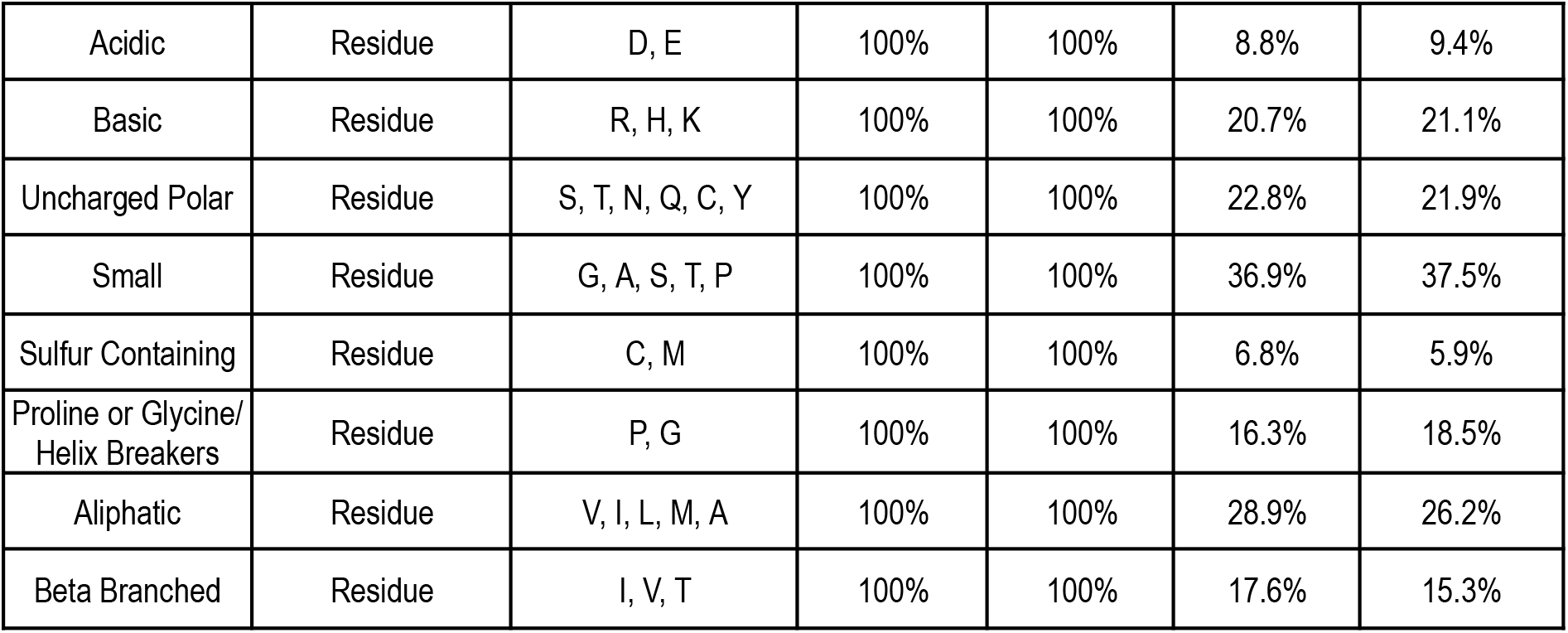
Summary of protein and residue properties. ^a^percent of available prediction.

| Attribute | Predictor | Threshold / Condition | Available Predictions |  | Positive Predictions<sup>a</sup> |  |
| --- | --- | --- | --- | --- | --- | --- |
|  |  |  | ClinVar_HQ (N=171,196) | ProteinGym (N=62,727) | ClinVar_HQ | ProteinGym |
| Disordered | ALBATROSS Metapredict v1.3<sup>109,110</sup> | 0.7 | 100% | 100% | 31.7% | 25.1% |
| PPI | PIONEER<sup>112</sup> | Binary | 74% | 77% | 9% | 12% |
| Homology (low) | UniRef90<sup>114,115</sup> | 10 | 89.7% | 88.3% | 8.5% | 6.8% |
| Length (long) | sequence | 1022 | 100% | 100% | 47.1% | 38.0% |
| Polar | Residue | S, T, C, Y, N, Q, G | 100% | 100% | 32.3% | 33.3% |
| Hydrophobic | Residue | A, V, I, L, M, F, Y, W | 100% | 100% | 33.2% | 31.4% |
| Charged | Residue | R, H, K, D, E | 100% | 100% | 29.6% | 30.5% |
| Aromatic | Residue | F, W, Y, H | 100% | 100% | 6.8% | 7.7% |
| Acidic | Residue | D, E | 100% | 100% | 8.8% | 9.4% |
| Basic | Residue | R, H, K | 100% | 100% | 20.7% | 21.1% |
| Uncharged Polar | Residue | S, T, N, Q, C, Y | 100% | 100% | 22.8% | 21.9% |
| Small | Residue | G, A, S, T, P | 100% | 100% | 36.9% | 37.5% |
| Sulfur Containing | Residue | C, M | 100% | 100% | 6.8% | 5.9% |
| Proline or Glycine/<br>Helix Breakers | Residue | P, G | 100% | 100% | 16.3% | 18.5% |
| Aliphatic | Residue | V, I, L, M, A | 100% | 100% | 28.9% | 26.2% |
| Beta Branched | Residue | I, V, T | 100% | 100% | 17.6% | 15.3% |

To enable comparison on a common scale, we map all scores to the [0,1] range using modified min-max normalization. Scores are flipped when necessary so that higher values indicate pathogenicity. This monotonic rescaling preserves discriminative performance (Methods). Optimal classification thresholds are derived globally and within each variant subgroup by maximizing the Youden J-statistic (Methods).

Classification thresholds vary substantially for variants in protein-protein interfaces (PPIs) and disordered regions across nearly all models, with many showing shifts in both subgroups (**Figure 1a**). Variants in sulfur-containing residues (methionine, cysteine) and proteins with few homologs also display consistent shifts (**Figure S1a, Table S1**) likely reflecting the influence of homology-based features incorporated in many models (**Table 1**). Among tested VEPs, FATHMM and SIFT are the most stable, showing no significant changes. Furthermore, shifts tend to be model-specific, for example, polar and aromatic residues show pronounced shifts in AlphaMissense and gMVP, less evident in other models (**Figure S1a**), likely reflecting the effect of structural features. Importantly, these shifts do not necessarily indicate a change in subgroup specific performance (**Figure S2a-b, Table S1**) but thresholds must be corrected to ensure consistent behavior across variant subgroups.

**Figure 1.**
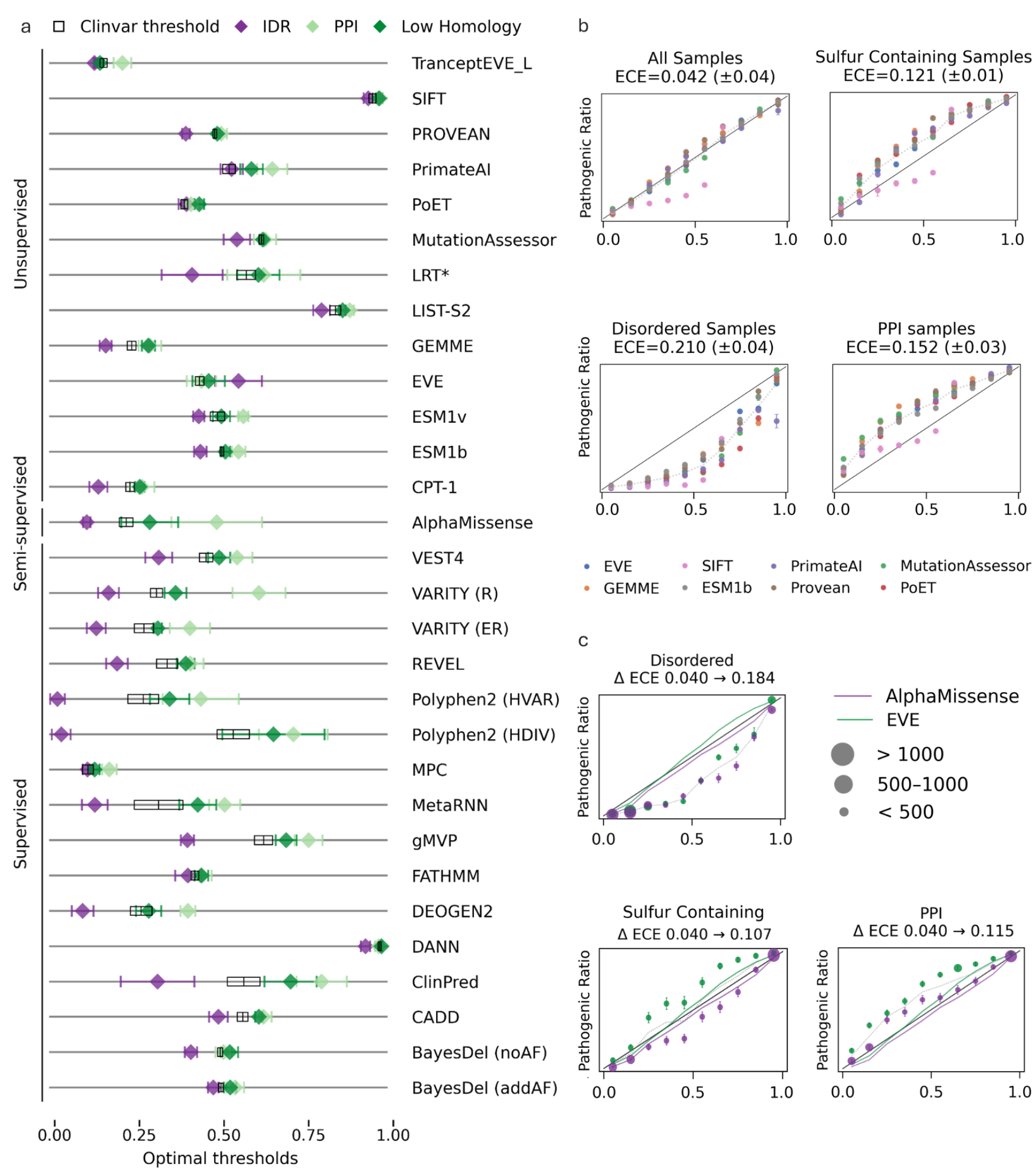
Residue-level miscalibration. **a.** Optimal classification thresholds shift substantially across residue subgroups (diamonds; maximal J-statistic) in ClinVar_BM dataset. Error bars indicate ±1 SD over 1,000 non-parametric bootstrap iterations. Black rectangle marks the naive threshold (mean ± SD) across ClinVar_BM, model scores normalized using modified min-max normalization, * log-transform applied to raw model scores (Methods) **b.** Globally calibrated models are well calibrated at the dataset level (top left) but miscalibrated within residue subgroups. Reliability histograms show predicted confidence (x-axis, 10 equal-width bins) versus observed pathogenic frequency (y-axis). Perfect calibration follows y=x; dotted lines show the mean trend across models. Circles mark per-model bin frequencies; error bars indicate ±1 SD across 100 global calibration iterations. Panels show the full ProteinGym dataset (top left), variants in sulfur-containing residues (top right), disordered variants (bottom left), and PPI variants (bottom right). ECE = Expected Calibration Error (mean across models). **c.** Miscalibration of EVE and AlphaMissense calibrated scores, with global ECE increasing at the residue level. Green and purple lines show calibration trends over the entire dataset for EVE and AlphaMissense, respectively; circle size denotes bin sample count. Other details as in (b).

As a ClinVar-derived dataset, ClinVar_BM contains N-terminal methionine (M1) variants that may disrupt translation initiation (n=164), most of which are pathogenic (n=148). Given their distinct mechanism, we provide additional analyses excluding M1 variants (**Figure S1b-c, Figure S2c-d**). M1 exclusion mainly attenuated sulfur-containing versus global threshold differences, reducing the number of models with non-overlapping ±1-SD intervals from 12 to 8. The largest gap reductions occurred in PolyPhen2 and PROVEAN, whereas gaps increased in AlphaMissense and VEST4. Global AUROC increased in 24/30 models, with the largest gains in DEOGEN2 and PROVEAN and more pronounced AUROC gains within sulfur-containing variants. Sequence-based models such as ESM1b, ESM1v, PoET and TranceptEVE showed little change in overall AUROC with M1 exclusion.

Despite these findings, the standard practice when using VEPs is to apply a single global threshold, which has also been shown to mask heterogeneous performance at the gene level^72^. Some methods allow more stringent cutoffs to tune precision with recent works suggesting to employ gene-level thresholds^73,74^, but to the best of our knowledge, no current framework recommends thresholds specific to different variant subgroups. Consequently, global thresholds provide no guarantees of performance within individual subgroups, an issue recently addressed by Fawzy et al. for variants in IDRs^75^. These observations underscore the advantage of residue-level calibration, as it captures variation that may be missed when calibration is performed only at the global or protein level.

### Globally and per-protein calibrated VEPs are not robust at the residue level

Multicalibrated models aim to provide reliable probabilities across diverse subpopulations. While some VEPs output probabilities inherently (SIFT, PolyPhen), others incorporate downstream calibration. EVE combines global and per-protein GMM calibration, whereas AlphaMissense applies a global logistic regression fitted on a subset of ClinVar variants. To assess the effect of global calibration, we calibrate raw scores of eight SOTA unsupervised VEPs using the ProteinGym Clinical Substitution benchmark. These models are trained using sequence-based, alignment-based, and structural features.

Simple global calibration was performed by training a logistic regression model on random subsets of 6,000 variants (∼10% of the dataset; Methods). To assess calibration quality, we use reliability histograms^76,77^, which plot model accuracy as a function of predicted confidence (Methods). For a perfectly calibrated model, the curve follows the identity line. Deviations above or below this line indicate overconfidence and underconfidence in predictions, respectively. We quantify miscalibration using the Expected Calibration Error (ECE), where ECE = 0 indicates perfect calibration and larger values reflect increasing miscalibration (Methods).

Following global calibration, almost all models achieve near-perfect calibration, with a mean ECE of 0.042 (**Figure 1b**, top left). SIFT retains a visible deviation, but its global ECE remains decent (∼0.096). Its scores are mostly concentrated in the well-calibrated edge bins, whereas the miscalibrated intermediate bins contain only approximately 20% of variants.

However when analysis is restricted to specific residue subgroups, including disordered regions, PPIs, and sulfur-containing residues, calibration consistently breaks down across all models (**Figure 1b**). Models show underconfidence in disordered residues and overconfidence in PPIs and sulfur-containing residues, even in the absence of M1 variants in ProteinGym. In these subgroups ECE increases by up to five folds relative to the globally calibrated baseline. Miscalibration worsens at lower prediction confidence, where accuracy deviates further from the identity line.

To establish that residue-level miscalibration is not an artifact of our calibration protocol or hyperparameter choices, we next examine the pre-calibrated scores provided by EVE and AlphaMissense (**Figure 1c**), using a subset of ClinVar variants not included in the AlphaMissense calibration (**ClinVar_AFM;** Methods, **Table 2**). While AlphaMissense employs a similar global calibration protocol, EVE uses a distinct hybrid calibration approach, combining global and protein-level calibration. Similar patterns persist for variants in disordered regions and PPIs, but AlphaMissense shows an opposite trend in sulfur-containing residues, exhibiting underconfidence (M1 variants are absent in ClinVar_AFM). Notably, AlphaMissense provides relatively well-calibrated scores for variants in PPIs, potentially reflecting the incorporation of structural features.These results motivate a calibration approach that explicitly corrects subgroup-specific reliability errors while preserving overall model calibration.

### Residue-level calibration using differential score mapping

To correct residue-level miscalibration without sacrificing the benefits of global calibration, we propose differential score mapping, a simple extension of the global logistic-regression calibration used above. Instead of fitting a single logistic rescaling function across all variants, we fit separate rescaling functions for predefined residue subgroups and apply the appropriate function according to each variant’s residue context, for example calibrating disordered and ordered residues independently. This strategy is designed to preserve calibration at the dataset level while correcting systematic reliability errors within subgroups.

Using the ProteinGym clinical substitution benchmark, we evaluate three calibration schemes based on residue subgroups that showed substantial miscalibration in our calibration analysis: (1) interface versus non-interface residues, (2) disordered versus ordered residues, and (3) a joint scheme further partitioning ordered residues into interface and non-interface groups (disordered residues were not subdivided further because of limited sample size). For each scheme, subgroup-specific logistic regression models were trained using 2,000 samples per group and evaluated on held-out variants (Methods).

This approach preserves global calibration at approximately the same level achieved by standard global calibration (ECE ≈ 0.040), while substantially reducing subgroup-specific miscalibration (**Figure 2a-b**). Subgroup-specific ECE closely matches the global ECE, decreasing from 0.210 to 0.050 in disordered residues and from 0.115 to 0.047 in PPI residues.

**Figure 2.**
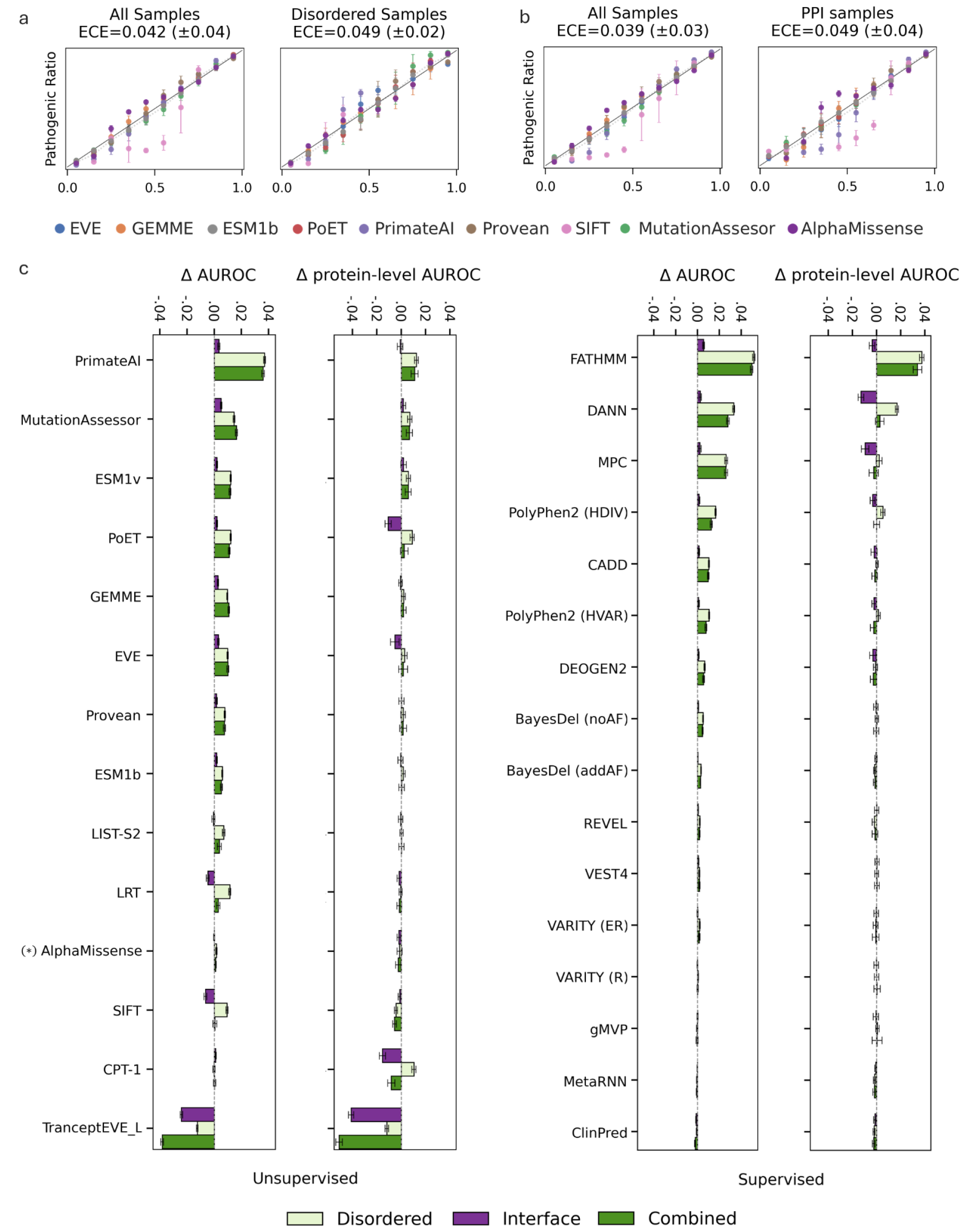
**Differential score mapping**. **a-b.** Differential score mapping corrects calibration within residue subgroups (right panels) while preserving global calibration (left panels). Models calibrated by fitting separate logistic regression functions using 2000 samples per subgroup on the ProteinGym clinical substitution benchmark, averaged over 100 calibration iterations. Reliability histograms are as described in Figure 1. **a.** Disorder vs. ordered residues **b**. interface vs. non-interface residues. **c.** Differential score mapping improves global AUROC (top) while preserving per-protein AUROC (bottom) across unsupervised (left) and supervised (right) VEPs. ΔAUROC and Δper-protein AUROC are reported relative to the uncalibrated model on the ProteinGym clinical substitution benchmark; error bars indicate ±1 SD over 1,000 bootstrap iterations. (*) semi-supervised.

Importantly, unlike global calibration, differential score mapping is not guaranteed to preserve global discriminative performance. Although each subgroup-specific mapping is monotonic within a subgroup, these mappings can alter the relative ordering of predictions across subgroups, thereby affecting AUROC. In practice, differential score mapping improves AUROC across nearly all unsupervised models, and many supervised predictors, without sacrificing per-protein AUROC (**Figure 2c**). The only exception is TranceptEVE_L, for which differential score mapping decreases AUROC.

However, choosing which residue subgroups should be calibrated separately remains non-trivial. ECE directly measures miscalibration, but it does not necessarily predict which subgroup partitions will improve AUROC after differential score mapping. Across models, disorder level and interface status are consistently among the most miscalibrated attributes (**Figure 3a, Figure S3**). Yet, calibrating interface residues yields much lower AUROC gains than calibrating disordered residues across nearly all models (**Figure 2c**). This mismatch is evident in MutationAssessor, EVE, GEMME, PROVEAN, and ESM1b, where interface residues show higher ECE than disordered residues but lower AUROC gains. TranceptEVE provides a more extreme example. It accounts for nine of the ten highest subgroup ECE values, including for disordered (ECE ≈ 0.29) and PPI residues (ECE ≈ 0.23; **Table S2**). Yet, its global AUROC decreases after differential score mapping (**Figure 2c**). As we next show, model score distributions can provide a more informative predictor of AUROC gains.

**Figure 3.**
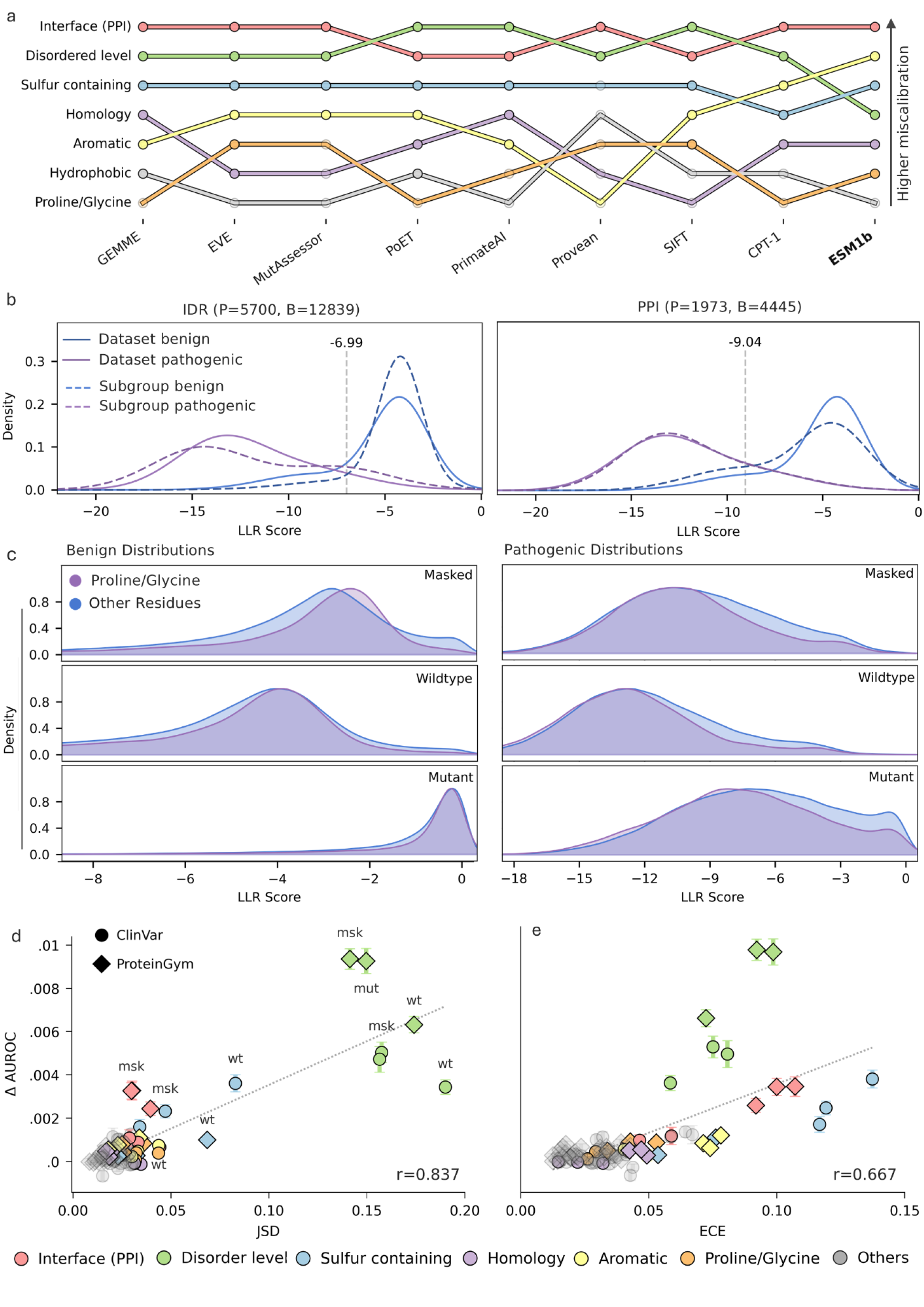
Miscalibration and score-distribution differences across residue subgroups. **a.** Calibration rank trajectories across selected residue attributes and predictors in ProteinGym. Attributes are ordered from highest (top) to lowest (bottom) ECE after global calibration. Opaque points indicate that the relative rank among the seven displayed attributes matches the absolute rank among all 15 attributes tested (Figure S3); Transparent points indicate that these ranks differ (e.g., sixth among the seven displayed attributes, but ninth among the full set of attributes). Interface, disordered, and sulfur-containing residues consistently rank among the most miscalibrated attributes. **b.** Class-conditional LLR-score distributions for disordered residues (left) and PPIs (right). Although both subgroups show pronounced miscalibration, only disordered residues show a substantial shift in score distributions. Distributions are fitted with two-component GMMs for the full dataset (dotted curves) and each subgroup (solid curves). *P* and *B* denote the numbers of pathogenic and benign variants within each subgroup, respectively. Dotted lines indicate optimal classification thresholds (J-statistic). **c.** LLR-score distributions for masked (top), wild-type (center), and mutant (bottom) sequence representations, grouped by benign (left) and pathogenic (right) variants. Differences between helix-breaking (proline/glycine) and other residues are amplified under the masked representation. **d-e** Change in global AUROC (y-axis; relative to the uncalibrated model) as a function of changes in score distribution (JSD) (d) and ECE (e). Changes in score-distribution better predict AUROC gains from differential subgroup calibration compared to ECE and are less dataset-dependent. Differential calibration was performed by fitting separate subgroup-specific logistic rescaling functions using one-third of the data for training and two-thirds for testing (Methods). Error bars indicate ±1 SD across 100 non-parametric bootstrap test iterations per fold. Circles and diamonds correspond to results on the ClinVar_Balanced ProteinGym clinical substitution benchmarks respectively. Colored symbols highlight selected calibrated attributes; transparent gray symbols denote other tested attributes (Table S3). wt, mut, and msk denote wild-type, mutant, and masked sequence representations, respectively. Dotted lines indicate linear regression fits.

### Subgroup-specific score distributions explain calibration-induced AUROC gains

To understand the ranking mismatch between ECE and calibration-induced AUROC gains across residue subgroups, we focus on ESM1b. ESM1b is particularly relevant because interface residues have the highest subgroup ECE, whereas disordered residues rank only fourth (**Figure 3a, Figure S3**). Nevertheless, calibrating for disordered residues produces a substantially larger AUROC gain than calibrating for interface residues, ∼0.01 compared with ∼ 0.002 (**Figure 2c**). An additional advantage of ESM1b, a PLM trained without clinical labels, is that its scores are less affected by dataset-specific biases.

We inspect the class-conditional distribution of model scores *P*(*X*|*M_i_*, *Y* = *y*) across ProteinGym and ClinVar (**Figure 3b, Figure S4**). Here, *M_i_* denotes a residue-level subgroup defined by a specific attribute, typically represented as a binary partition (e.g., disordered vs. ordered residues), *X* denotes a model score such as the log-likelihood ratio (LLR) and *Y* denotes the variant label. We analyze scores separately for benign *P*(*X*|*M_i_*, *Y* = 0) and pathogenic variants *P*(*X*|*M_i_*, *Y* = 1) because they each occupy different regions of the score axis (**Figure S4-5**). To control for differences in class proportions between subgroups, we resample variants within each subgroup to match the global pathogenic fraction of the dataset. Since differential score mapping fits calibration functions over model scores, a subgroup whose score distribution diverges strongly from the global distribution is expected to require a distinct regressor, motivating the use of score distributions for calibration target selection.

In alignment with AUROC gains, variants in IDRs show the strongest subgroup-specific score-distribution differences, with both benign and pathogenic distributions shifted relative to the global distributions (**Figure 3b, Figure S4**). In contrast, despite having the highest ECE, PPI residues show more modest differences. Other residue attributes, including sulfur-containing residues, polar residues, and proteins with few homologs, show weaker score-distribution differences (**Figure S4**). Similar patterns are observed in the ProteinGym clinical substitution benchmark (**Figure S4**).

An important nuance of PLM score distributions is the effect of different input representations, which may affect the conditional score-distribution divergence. LLR scores can be computed from the wild-type sequence, the mutant sequence, or a masked sequence in which the mutated position is replaced by a mask token (Methods). To demonstrate this effect, we compare subgroup-specific LLR distributions across these three representations (**Figure 3c, Figure S5**).

Variants in PPI and disordered residues show the clearest separation under the wild-type representation (**Figure S5**). In contrast, differences between Proline/Glycine residues, corresponding to helix-breaking amino acids, and other residues are most pronounced under masked marginals (**Figure 3c**). Previous work reported limited benefit of masked representations for variant effect prediction, often not justifying their additional computational cost^21,78^. However, our results indicate that input representation may play an important role in revealing subgroup-specific differences and should be considered in the context of calibration.

Putting these observations together, we compare AUROC gains from differential score mapping across residue subgroups, datasets, and input representations with both ECE and score-distribution differences (**Figure 3d-e**). To estimate score-distribution differences, we use the Jensen-Shannon divergence (JSD). In brief, we compute the JSD between a subgroup and complementary LLR distributions separately for benign and pathogenic variants, and sum the two terms (Methods, Appendix 1):

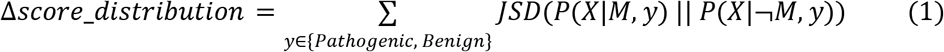

Here, *X* denotes the distribution of LLR values. For example, for disordered residues (*M*) versus ordered residues (¬*M*). Score distribution differences show a strong association with calibration-induced AUROC gains (*r* = 0. 84, *p* < 1*e* − 24; **Table S3**), whereas ECE is substantially less predictive (*r* = 0. 67, *p* < 1*e* − 12). Score distributions mostly resolve the ranking mismatch observed with ECE. Using ECE, sulfur-containing and interface residues rank above disordered residues despite their smaller AUROC gains (**Figure 3e**). In contrast, using score-distribution differences, the ranking better matches the AUROC gains: disordered residues show the highest divergence, followed by sulfur-containing and interface residues in close proximity (**Figure 3d**). Importantly, Score-distributions differences also appear less dataset-dependent than ECE. For instance, using ECE, sulfur-containing and interface residues form distinct ClinVar and ProteinGym clusters, whereas using score-distribution differences they cluster more by attribute than by dataset (**Figure 3d-e**).

Importantly, M1 variants account for approximately 20% of sulfur-containing variants in ClinVar_Balanced (634/3,133). Their exclusion materially reduces miscalibration within this group, approximately halving ECE and maximal calibration error (MCE; Methods) to 0.07 and 0.21, respectively, approaching the values observed in ProteinGym, where M1 variants are absent (**Figure S6c**). Nevertheless, sulfur-containing residues remain among the most miscalibrated attributes (**Figure S3a–b; Table S2**). ECE understates this pattern because it is dominated by relatively well-calibrated edge bins, whereas the reliability histogram reveals persistent deviations. MCE indicates appreciable residual miscalibration, comparable in magnitude to that of disordered residues (MCE = 0.28; **Table S3**). Calibration-induced AUROC gains are attenuated but persist after M1 exclusion. Under wild-type marginals, they closely match those for interface, aromatic, and beta-branched residues at approximately 0.001 (**Figure S6; Table S3**). JSD shows minimal change with M1 exclusion, highlighting its robustness to dataset composition (**Figure 3d, Figure S6a**). Extending this analysis across models, ECE rankings remain largely preserved (mean Kendall’s τ = 0.724; **Figure S3a,b; Table S2**), with the exception of CPT-1, for which sulfur-containing residues shift from second to sixth.

Finally, Score-distribution differences also require much less labeled data than either ECE or direct ΔAUROC-based target selection. Estimating ECE for a candidate subgroup requires fitting a calibration regressor and retaining enough held-out variants to estimate bin-level pathogenic frequencies, which is sensitive to sample size^39,79^. Directly using ΔAUROC is even more data-demanding, requiring fitting separate subgroup-specific regressors and preserving sufficient test data to estimate the resulting AUROC gain. By contrast, score-distribution divergence can be estimated directly in score space before fitting subgroup-specific calibration functions and, as we show below, can be estimated with as few as 400 samples per class.

### RaCoon - Residue-aware Calibration of conditional distributions

Leveraging our findings we sought to design a calibrated VEP that guarantees consistent performance across variant subgroups. In addition, we aim to develop a clinically interpretable model capable of producing meaningful pathogenicity probabilities. We chose ESM1b as the base model for RaCoon due to its strong performance, unsupervised nature, and lack of reliance on MSAs^4,8,9^.

Our calibration approach (**Figure 4**) targets three binary residue attributes supported by the score-distribution analysis and calibration-induced AUROC gains across both datasets (**Figure 3d**): disorder level, interface status, and sulfur-containing residues (methionine or cysteine versus other residues). These attributes also show consistent miscalibration across models and datasets (**Figure 3a, Figure S3**). We use wild-type marginal scores given their widespread adoption and pronounced score-distribution differences across these residue subgroups. Because inference on long proteins in ESM1b (>1,022 residues) requires a sliding-window approach (Methods), which has been shown to affect performance^4,60,80^, we treat protein length as a separate calibration subgroup. Accordingly, we introduce an additional technical division between residues in long and short proteins.

**Figure 4.**
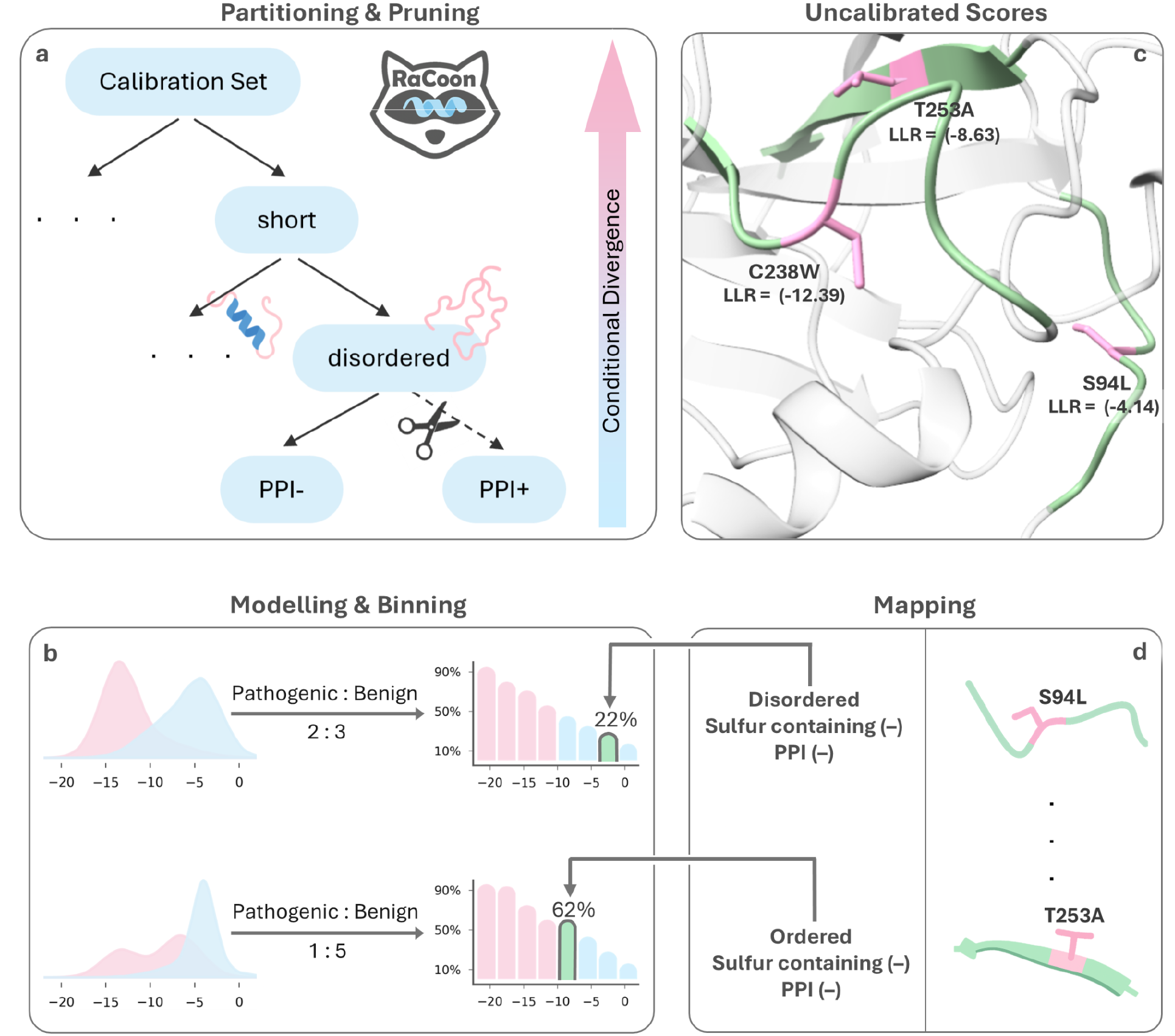
The RaCoon pipeline. **(a)** A calibration set is partitioned across binary residue attributes, with attributes showing greater class-conditional score-distribution divergence placed closer to the root (color-graded arrow). Leaves with insufficient data are pruned (scissors icon). **(b)** Nodes’ pathogenic and benign LLR score distributions are modeled using two GMMs and the proportion of pathogenic variants is estimated. Synthetic samples are subsequently drawn from the GMMs according to the estimated pathogenic-to-benign ratio to construct calibration histograms (x-axis LLR scores, y-axis pathogenic fraction). **(c)** Uncalibrated per-variant LLR scores (pink sticks, PDB 2pcx) are obtained from ESM1b. **(d)** Each variant is mapped to its node and histogram bin to yield a residue-level calibrated pathogenicity probability.

The RaCoon pipeline consists of four main steps (Methods). First, attribute combinations are partitioned into a calibration tree (**Figure S11**), where each node represents a unique set of binary residue attributes. The partitioning process is guided by class-conditional subgroup-specific score distributions, placing attributes with stronger divergence near the root to prioritize partitions more likely to improve AUROC as previously shown (**Figure 3d**). Nodes with insufficient data are subsequently pruned (**Figure 4a**). Second, per-node score distributions are modeled by fitting two GMMs to the LLR scores of pathogenic and benign variants. Third, LLR scores are binned into pathogenicity histograms by sampling from the GMMs according to the estimated node’s proportion of pathogenic variants. Each bin expresses an estimated pathogenic percentage for a defined LLR range (**Figure 4b**), providing a frequency-based representation intended to support clinical interpretation^81,82^. Finally, at inference time, variants are mapped to their corresponding node and histogram bin, mapping LLR scores to interpretable calibrated probability scores (**Figure 4c-d**).

### GMM-based sampling provides efficient calibration with minimal labeled data

A key novelty of RaCoon is the use of GMM-based sampling to construct calibration histograms rather than direct binning of labeled samples. Standard calibration approaches directly estimate *P*(*Y* = *y*| *X*, *Mi*) where *Y* is the pathogenic label, *X* the LLR score and *Mi* the residue subgroup. However, direct estimation requires substantial labeled data within each subgroup, which is impractical for the sparse residue subgroups considered by RaCoon.

Instead, RaCoon estimates the score distributions of pathogenic and benign variants within each residue subgroup and derives posterior pathogenicity probabilities using Bayes’ rule (Equation 4):

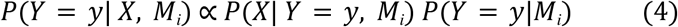

Here, *P*(*X*| *Y* = *y*, *M_i_*) is estimated by fitting GMMs, while *P*(*Y* = *y*|*M_i_*) is estimated empirically from a limited number of variants (Methods). The normalization term *P*(*X* | *M_i_*) is estimated by marginalizing over the two class labels (Equation 4.1):

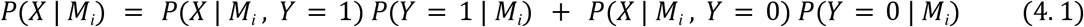

Fitting separate GMMs reflects the distinct score distributions of benign and pathogenic variants (**Figure 3b, Figures S4–S5**) and permits probability mappings beyond a single sigmoid^83^. We show that GMM-only calibration with as few as 100 training samples per GMM performs better than direct binning, which requires tens of thousands of labeled variants. We hypothesize that this reflects a denoising effect of the GMMs. By fitting a small number of parameters (two means), they capture the dominant structure of the score distribution while suppressing noise from individual samples. Consistent with this interpretation, a hybrid approach that combines GMM sampling with direct binning shows progressively degraded performance as more raw samples are introduced, indicating that these samples primarily add noise rather than useful signal (**Figure S12a**). We also compared GMM-based histogram calibration with logistic regression fitted separately within each calibration node. While AUROC for GMM-based calibration continues to improve with training size, logistic-regression AUROC plateaus early (**Figure S12b**; Methods). However, because pruning thresholds are set according to the number of training samples required per GMM to ensure sufficient data for both training and held-out evaluation, increasing GMM training size reduces the number of retained calibration subgroups (**Figure S12b**). To balance accuracy with subgroup coverage, an important consideration for multicalibration, we set the final training size to 400 samples per GMM, which preserves most calibration nodes.

Importantly, while RaCoon uses labeled examples to estimate class-conditional score distributions and subgroup pathogenicity priors, its AUROC performance does not increase when it is evaluated on the examples used for calibration (Appendix 3). We hypothesize that this reflects the GMMs modeling aggregate pathogenic and benign score distributions, thereby limiting sensitivity to individual calibration examples. Although the estimated pathogenic fractions remain susceptible to dataset-specific annotation biases, they may also capture important biological signals that we aim to model. As shown in Appendices 1-2, restricting calibration to attributes with strong score-distribution divergence, together with pruning nodes with insufficient data, increases the likelihood that these distributional signals and priors reflect genuine biology rather than noise. As we next show, RaCoon maintains consistent performance when calibrated and evaluated across independent datasets, suggesting that its calibration is not driven primarily by dataset-specific biases.

### RaCoon outperforms ESM1b on clinical and experimental datasets while improving calibration

To evaluate RaCoon against the baseline ESM1b, we report global and per-protein AUROC on both experimental and clinical datasets (Methods). While ClinVar_BM provides a robust benchmark with reduced gene-level annotation bias, it is too small to construct deep calibration trees using RaCoon. Instead, for clinical data we use ClinVar_Balanced, which bounds protein-level label imbalance to reduce such biases while retaining enough labeled examples to build deeper calibration trees (Methods, Table 2). Additionally, we evaluate performance on a stringent subset of 46 well-annotated, balanced ClinVar proteins, each containing at least 100 annotations and a pathogenic to benign label ratio not exceeding 70:30. For experimental data, we use the ProteinGym Clinical Substitution benchmark. All reported values are based on 100 random calibration iterations, each evaluated using 100 independent non-parametric bootstrap test iterations. Classification thresholds are determined by maximizing the Youden J-statistic.

We first evaluate RaCoon across datasets by calibrating on ClinVar_Balanced and testing on ProteinGym (**Figure 5a-b**). This setting reduces the possibility that AUROC gains arise from adaptation to dataset-specific priors in the calibration set. Under this evaluation, RaCoon improves global AUROC from 0.910 ± 0.001 for raw ESM1b LLR scores to 0.918 ± 0.001, with improvements observed in all 100 calibration iterations. RaCoon also yields a modest but significant improvement in per-protein AUROC, from 0.895 ± 0.000 to 0.897 ± 0.001, outperforming raw ESM1b in 98 of 100 calibration iterations. Excluding M1 variants from the ClinVar_Balanced calibration set preserved the global and per-protein AUROC gains on ProteinGym (**Figure S13a–b**). These findings demonstrate that the calibration procedure generalizes across datasets.

**Figure 5.**
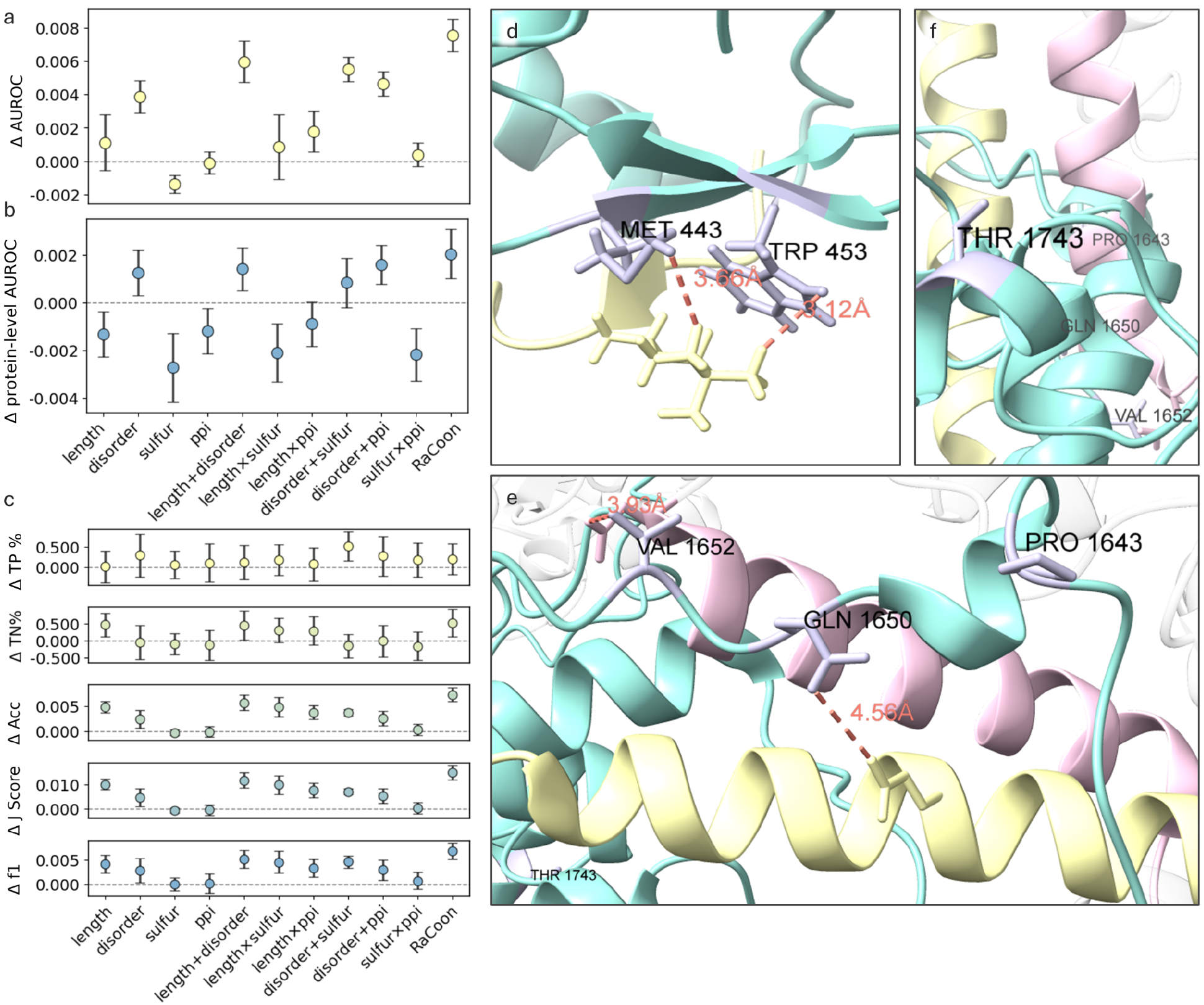
RaCoon performance evaluation across datasets. a–c. Changes in performance on the ProteinGym clinical substitution benchmark after calibration on ClinVar_Balanced, relative to uncalibrated ESM1b. RaCoon improves both global and per-protein AUROC. Error bars indicate ±1 SD across 100 randomized calibration iterations, each evaluated using 100 non-parametric bootstrap test resamples. (a) overall AUROC (b) per-protein AUROC (c) Classification metrics. Single and paired attributes on the x-axis indicate the attributes used for calibration; “disorder” denotes ordered versus disordered residues, and “RaCoon” denotes the full pipeline. TP = true positive; TN = true negative. Classification thresholds were optimized per model using the Youden J-statistic. **d-f.** Structural examples of variants correctly reclassified after calibration in RAG2 (PDB 8T4R, d) and ARID1A (PDB 6LTJ, e-f). ClinVar-annotated variants (purple sticks) are shown with interface distances (orange, Å). Interface partners: yellow - Histone H3 (RAG2) and SMCE1 (ARID1A); pink - SMRD1.

We next evaluated RaCoon on a subset of 46 well-annotated, strictly balanced proteins whose variants were not seen during calibration. These proteins satisfied a stricter balance criterion than ClinVar_Balanced, with a label ratio not exceeding 70:30. Notably, 27 of the 46 proteins were close to perfectly balanced (label ratio ≤ 60:40). Although these variants were still derived from ClinVar, this setting substantially limited the transfer of gene-level priors from the calibration set. In this setting, RaCoon significantly improved AUROC in 12 proteins while degrading performance in only 4 (**Figure S14,** Methods), increasing global AUROC from 0.918 ± 0.002 to 0.927 ± 0.002 and per-protein AUROC from 0.929 ± 0.002 to 0.933 ± 0.002. RaCoon improved global AUROC in all 100 calibration iterations and per-protein AUROC in 99 of 100 iterations. Removing the 47 M1 variants had little effect on the results, preserving the global and mean per-protein AUROC gains and significant improvements in the same 12 proteins. One additional protein, GRIN2A, showed a marginally significant degradation relative to the uncalibrated scores (q = 0.0499; **Figure S14b**).

Finally, we evaluate RaCoon within-dataset - where calibration and test sets are derived from the same benchmark (**Figure S15**). While this setting risks inflating AUROC by exposing the model to dataset-specific priors during calibration, it remains an important practical benchmark, as calibration is often used to adapt a model to a specific dataset. On ClinVar_Balanced, RaCoon improves AUROC to 0.919 ± 0.001 from 0.911 ± 0.001. These gains were also retained when M1 variants were excluded from ClinVar_Balanced before partitioning into calibration and test sets (**Figure S13d-e**). A similar improvement is observed on ProteinGym, where RaCoon reaches 0.925 ± 0.001 compared with 0.912 ± 0.000 for the uncalibrated model. Per-protein AUROC also improves modestly but significantly on both ClinVar_Balanced (0.916 ± 0.002 vs. 0.913 ± 0.001) and ProteinGym (0.909 ± 0.002 vs. 0.903 ± 0.001). Importantly, when stratified by protein pathogenic label ratio, RaCoon outperforms or matches ESM1b across nearly all bins in both within-dataset settings, including the more balanced bins (**Figure S16**), further supporting that the observed gains are not primarily driven by memorization of protein-level label priors. Across all test-sets, the ROC curve of RaCoon consistently lies above that of the uncalibrated model, indicating superior performance across classification thresholds (**Figure S17**).

Although calibration on ClinVar_Balanced is designed to reduce protein-level pathogenic-fraction bias, residual bias may still remain. To test whether the observed gains are driven by learned protein-level pathogenic proportions, we perform a label-ratio permutation analysis (**Table S6**, Methods), in which pathogenic fractions are permuted among test proteins by resampling variants within each protein to match the assigned fraction. RaCoon was calibrated once on the original, unpermuted ClinVar_Balanced dataset and applied to all permutations without refitting. This is a particularly stringent setting because RaCoon cannot adapt to the reassigned protein-level priors, while the permutation also disrupts the subgroup-specific pathogenicity priors learned during calibration. Even under this stringent control, across 1,000 permutations, RaCoon consistently outperformed uncalibrated ESM1b in nearly all head-to-head comparisons, with a mean AUROC improvement of approximately 0.003 ± 0.001 across all test sets.

A key property of RaCoon is its ability to provide multicalibrated predictions across residue subgroups. To evaluate calibration quality directly, we report ECE and MCE for the 13 retained calibration nodes (Methods). M1 variants were excluded from these analyses. RaCoon achieves consistently low calibration error across these subgroups within and across datasets (**Figure S11**, **Table S7**). On ClinVar_HQ, all 13 nodes achieve ECE ≤ 0.05, and all but one achieve MCE < 0.20. When calibrated on ClinVar_HQ and tested on ProteinGym, 11 of 13 nodes achieve ECE < 0.08, with the remaining two at 0.09 and 0.13. By comparison, naive global logistic calibration yields MCE ≥ 0.25 in 8 of 13 nodes (**Table S7**). Including M1 variants yielded similar results: all 13 ClinVar_HQ nodes achieved ECE ≤ 0.05 and all but one achieved MCE < 0.20, while global logistic calibration yielded MCE ≥ 0.25 in 7 of 13 nodes (**Table S7**).

### RaCoon improves borderline variant interpretation in RAG2 and ARID1A

Complete calibration yields additional AUROC gains and significantly improves threshold-based metrics, enhancing discrimination of both benign and pathogenic variants. These benefits are most apparent for borderline cases, where misclassification risk is highest, as illustrated by example cases in RAG2 and ARID1A (**Figure 5d-f**).

RAG2 is a core V(D)J recombinase component whose mutations cause severe immunodeficiencies^84–86^. Two variants at residue M443 (M443I pathogenic, M443T likely-pathogenic^87,88^), are misclassified as benign by ESM1b raw LLR scores. RaCoon correctly identifies them as pathogenic, assigning interpretable and calibrated probabilities of 73% (M443I) and 74% (M443T). This improvement reflects methionine’s classification as a sulfur-containing residue, which places these variants into a distinct calibration subgroup. In contrast, a nearby W453 variant (non-sulfur containing) affecting the same interaction is correctly classified as pathogenic by both models.

Context dependence is illustrated by the LLR scores mappings: M443I (-7.03) maps to 73% pathogenic probability, whereas W453R, assigned to a different node, receives a more negative LLR (−9.25) yet a similar calibrated probability (74%). Although more negative LLRs typically imply stronger pathogenicity, this example shows how RaCoon adjusts predictions according to local distributional context rather than LLR magnitude alone.

An opposite effect is seen in ARID1A, where calibration correctly reclassifies 12 variants from pathogenic to benign. ARID1A is a core subunit of the BAF (SWI/SNF) chromatin-remodeling complex^89^ and a major tumor suppressor recurrently altered across cancers^90–93^. Its large size, long disordered regions, and multiple PPIs make variant interpretation particularly challenging. Two benign ClinVar variants, P1643T and T1743M (**Figure 5e-f**), receive uncalibrated LLRs in the pathogenic range (−8.91 and −8.50) but are correctly reclassified as benign once residue context is incorporated. Notably, P1643T lies near the SMARCD1/SMARCE1 interface, yet RaCoon maps it to a pathogenicity ratio of 29%. Nearby variants (Q1650H, V1652A) remain misclassified, likely due to predicted interface involvement, but their calibrated probabilities (46% and 44%) are borderline. For comparison, an interface-associated variant in KCNQ2 (Q341H), with a comparable LLR (−9.05), receives a much higher calibrated pathogenicity of 68%. Together, these examples illustrate how region-specific score mappings can yield different pathogenicity estimates for similar LLRs, highlighting the importance of local residue context.

### Ablation studies identify specific contributors to performance gains

To disentangle the contributions of individual components in our calibration pipeline, we compare naive ESM1b to our calibrated model under multiple single and pairwise feature ablations (**Figure 5a-b, Figure S12)**. In addition to AUROC, we report complementary binary classification metrics, all averaged over repeated training and bootstrap iterations, and reported as changes relative to the uncalibrated ESM1b on variants not seen during the calibration process.

Calibration to protein length and disordered residues emerge as key contributors to AUROC across all datasets, with largely additive effects. Notably, differences in protein length that are not reflected in the class-conditional score distribution, remain important, underscoring the need to correct for context length in transformer-based models. Pairwise calibration to disordered residues and sulfur-containing residues also yielded consistent gains.

While several attributes contributed to AUROC improvement, their impact at the optimal classification threshold are more subtle (**Figure 5c**), illustrating the limitations of global thresholds. AUROC-based gains primarily reflect improved global ranking^94,95^, whereas per-protein AUROC shows smaller gains. Per-protein AUROC is a less stable metric for proteins with a few labeled variants^96,97^ and does not account for inter-protein ordering, imperative for genome-wide screening^4^.

Finally, we assess whether the calibration step gains AUROC performance by memorizing specific examples. We compare RaCoon’s performance when evaluated on variants used to fit the per-node GMMs with its performance on matched held-out variants with the same pathogenic-to-benign ratio (Appendix 3, Methods). Even under this deliberate-leakage setting, AUROC remains essentially unchanged on both ClinVar_Balanced and ProteinGym.

### RaCoon open-access server and clinical interpretation

RaCoon is publicly available through an open-access web server and GitHub repository. The server reports calibrated pathogenicity estimates together with prediction uncertainty and subgroup-specific calibration metrics. We note that RaCoon calibrated estimates should not be interpreted as posterior probabilities of pathogenicity under the ACMG/AMP framework. To support interpretation within that framework, the server also reports the corresponding PP3/BP4 evidence category based on the ClinGen computational-evidence calibration framework^29,30^ (Table S8; Methods).

## Discussion

Recent years have seen a significant progress in missense variant effect prediction, leading to growing adoption in both research and clinical variant classification^29,47^. As VEPs become more deeply integrated into clinical workflows, the need for reliable interpretation frameworks that account for biases in training and evaluation has grown urgent. Various approaches have been proposed to calibrate VEPs^29,34^, with global calibration on specific datasets and gene-level calibration among the most widely used^1,6^.

However, few studies have examined the robustness of current calibration approaches at the residue level. We observe widespread shifts in optimal classification thresholds across residue subgroups (**Figure 1a, Figure S1**). This extends to similar gene-level observations reported by Tejura et al. (2024)^72^ at the residue-level, raising concrete concerns about the use of global classification thresholds and highlighting the potential value of incorporating residue-level assessments into benchmarking protocols. Furthermore, our analysis shows that both global and gene-level calibration degrade substantially in reliability when evaluated within specific residue subgroups, most notably variants in disordered regions, protein-protein interfaces, and sulfur-containing residues (**Figure 1b-c**). Together, these findings motivated the development of calibration approaches that can capture residue-level variation.

We present a multicalibration approach based on differential score mapping (**Figure 2**) which learns separate rescaling functions for predefined residue subgroups. Importantly, unlike standard global calibration approaches^39^, differential score mapping can change the global ordering of variants across subgroups and therefore affect model discrimination. When calibration targets are chosen appropriately, it not only preserves AUROC but also improves it across models (**Figure 2c**). Although these improvements are generally modest, they are significant, consistent across models and datasets, and achieved without additional model training. We do not view this as merely a feature of differential score mapping, but as possible evidence that calibration can correct subgroup-dependent misordering caused by distributional differences.

This raises a central question: how should calibration targets be selected? In the calibration literature, miscalibration is often associated with two main types of distribution shift: label shift^98–100^ and shifts in model score distribution^99,101–103^. A key novelty of our framework is the use of subgroup-specific score distributions to detect these shifts and guide calibration. Compared with label frequencies or directly using ECE estimates, score distributions offer several advantages. First, they are less affected by labelling biases, especially for unsupervised models, providing a more reliable signal to detect distribution shifts (**Figure S7**, Appendix 1). Second, they capture model-specific rather than dataset-specific shifts (**Figure 3d-e**). Third, they can reveal how different encoding strategies capture subgroup variation (**Figure 3c**). Consistent with this, we find that changes in model score distributions are strong predictors of calibration targets likely to improve AUROC (**Figure 3d, Figure S8**). These advantages extend beyond residue-level calibration and could also be used to guide other calibration schemes, such as identifying informative gene-level subgroups.

Importantly, while our manuscript introduces the added value residue-level calibration, there are yet important biological signals that can only be captured at the gene-level such as inheritance pattern or disease prevalence^14,104^. Although this is outside the scope of the present study combining residue and gene level calibration would likely capture additional signals. A natural extension of this work would be to explore whether differential score mapping at the gene level, or hybrid residue-gene calibration, can provide further gains in model accuracy. Notably, as differential mapping only affects ordering across subgroups, similar gains from protein-level calibration would be captured using global AUROC rather than per-protein AUROC.

Building on these insights, we introduce RaCoon (**Figure 4**), a multicalibrated variant effect predictor based on ESM1b^4,105^ that applies residue-level calibration across key attributes identified in this study. The framework learns class-conditional LLR distributions using GMMs and converts them into interpretable pathogenicity probabilities through calibration histograms. A central novelty of RaCoon is the use of GMM sampling to construct calibration histograms, which reduces the training set size required for stable calibration from thousands of labeled variants to only a few hundred samples. This provides a practical foundation for applying histogram-based calibration in other settings as well. Beyond delivering multicalibrated and interpretable predictions, RaCoon also improves AUROC relative to the uncalibrated model across benchmarks (**Figure 5**). RaCoon can be readily extended to other unsupervised VEPs, enabling calibration guided by model-specific conditional score distributions.

Importantly, RaCoon’s calibrated probabilities should not, on their own, be interpreted as clinical evidence or as standalone variant classifications. Consistent with the subgroup-dependent threshold shifts identified here, developing subgroup-specific mappings to ACMG/AMP evidence strengths, rather than relying on a single global mapping, represents an important direction for future work. Nevertheless, to support clinical interpretation and public use, we provide a global ClinGen-based mapping of RaCoon outputs to PP3/BP4 evidence categories through the open-access RaCoon server (Table S8). These categories should be integrated with other evidence under the ACMG/AMP framework^29^.

## Author contributions

Conceptualization was jointly carried out by G.P., S.A., and D.S. Methodology, data curation, benchmarking, and statistical analyses were carried out by G.P. RaCoon was jointly developed and validated by G.P. and S.A. S.A. led the software development and implementation of RaCoon and its web server and carried out most of the ablation studies. All authors contributed to writing and revising the manuscript. D.S. supervised the study.

## Supporting information

Supplementary tables columns description

## Acknowledgements

We thank Haimasree Bhattacharya for the development of the RaCoon web-server. D.S is supported by the Israeli Science Foundation (ISF 737/23), the European Research Council (ERC) under the European Union’s Horizon Europe research and innovation programme (ERC Consolidator Grant 101171306), the European Innovation Council (EIC) Pathfinder programme (Project 101130574), and the Israeli Ministry of Science and Technology (MOST 4707). G.P. is supported by the Azrieli Foundation through the Azrieli Graduate Studies Fellowship. The funders had no role in study design, data collection and analysis, decision to publish, or preparation of the manuscript. Molecular graphics and analyses were performed with UCSF ChimeraX, developed by the Resource for Biocomputing, Visualization, and Informatics at the University of California, San Francisco, with support from National Institutes of Health R01-GM129325 and the Office of Cyber Infrastructure and Computational Biology, National Institute of Allergy and Infectious Diseases.

## Methods

### Datasets curation (Table 2)

#### ClinVar_HQ

The complete ClinVar^7^ data was downloaded in GRCh38 VCF format (July 2025). Records missing core ClinVar annotations (CLNSIG, CLNVC, CLNREVSTAT, or CLNHGVS) were excluded. We retained only single-nucleotide variants and required a review status ≥1 star, based on the CLNREVSTAT stars mapping. This filtering step removed conflicting variants, variants lacking clinical classification, and those with no assertion criteria, yielding 1,751,722 variants. Next, all transcript identifiers were resolved using the Entrez API^106^, and only variants with valid protein-level variant symbols were retained (that is, those beginning and ending with canonical amino acids whose wild-type residue matched the reference transcript). Variants of uncertain significance were excluded, retaining only variants annotated as Likely pathogenic, Pathogenic/Likely pathogenic, Pathogenic, Benign, Benign/Likely benign, or Likely benign. These categories were mapped to binary labels. Finally, we unified duplicate variants, sharing the same binary label in the same transcripts and removed variants with contradicting labels. The resulting ClinVar_HQ dataset contained 171,196 variants across 14,564 unique protein sequences, of which 52,634 were pathogenic and 118,562 benign.

#### ClinVar_BM

This dataset represents a rigorously filtered subset of ClinVar designed for benchmarking VEPs and minimizing circularity and annotation biases. We use the dataset recently published by Radjasandirane *et al.*^9^, which includes 14,053 variants (11,757 benign and 2,296 pathogenic), all published after May 1, 2021. Variants that also appeared in the Humsavar dataset^107^ or showed unbalanced annotations (ratio exceeding 60/40) were excluded. All remaining variants were verified to represent valid protein-level alterations. Because transcript identifiers were not provided, variants were mapped to sequences using gene symbols, leveraging the transcript mappings established for the ClinVar_HQ dataset.

#### ClinVar_Balanced

This dataset is an extension of ClinVar_BM for unsupervised VEPs which do not require the removal of potential training data. Curated from ClinVar_HQ - we retain only proteins with at least six annotated variants and enforce a balanced label distribution at the protein level, requiring a pathogenic-to-benign ratio between 80:20 and 20:80. The resulting dataset contains 40,474 variants (19,526 pathogenic and 20,948 benign) across 1,337 unique protein sequences. The dataset is designed to increase sample size while remaining robust to gene-level biases in AUROC analysis.

#### Balanced Proteins Subset

This dataset is a robust subset of ClinVar_Balanced used to evaluate RaCoon’s gene-level performance. We retain only proteins with at least 100 annotations and a pathogenic-to-benign ratio between 70:30 and 30:70. The resulting dataset contains 9,354 variants (4,390 pathogenic and 4,964 benign) across 46 unique protein sequences.

#### ClinVar_AFM

AlphaMissense, published in 2023, includes a supervised calibration process that could potentially introduce variants present in the ClinVar_BM dataset. To prevent such circularity, we used a subset of ClinVar variants published by Cheng et al.^6^ that were explicitly excluded from the AlphaMissense calibration process. We removed variants for which no matching transcript could be found in the ClinVar_HQ dataset, ensuring that all remaining variants met our established filtration criteria. The final dataset consists of 56,368 variants, including 18,765 pathogenic and 37,603 benign variants across 6,272 unique protein sequences.

#### Well-annotated ClinVar proteins

This dataset was compiled directly from the ClinVar_HQ dataset, retaining only sequences with at least ten variants, of which we required a minimum of four pathogenic and four benign variants. The final dataset consists of 42,056 pathogenic and 44,155 benign variants across 1,285 protein sequences.

#### ProteinGym

Throughout the study we use the ProteinGym zero-shot clinical substitution benchmark^8^ obtained in June 2025. For attribute analysis we use the provided protein sequences, labels are assigned using the *DMS_bin_score* column. The benchmark contains 62,727 variants (32,000 pathogenic and 30,727 benign) across 2,525 unique protein sequences.

#### Exclusion of M1 variants

As a supplementary sensitivity analysis, we excluded N-terminal methionine (M1) variants from ClinVar-derived datasets containing these variants because of their distinct potential effects on translation initiation. Exclusion was performed before partitioning the data into training/calibration and test sets, ensuring that M1 variants were absent from both model fitting and evaluation.

#### VEPs scores

were mostly computed using Ensembl VEP^108^, EVE, CPT-1 and AlphaMissense that were queried directly from the published precomputed scores. GEMME and PoET were inferred directly from the models using MSAs obtained from the AlphaFold server. For the ProteinGym clinical substitution benchmark, we used the provided precomputed scores for all models except ESM1b. ESM1b scores were computed directly using the *’esm1b_t33_650M_UR50S’* model, with LLR scores calculated as described below.

### Log Likelihood Ratio Scores (LLR)

We use the LLR score derived from ESM1b as a quantitative measure of predicted pathogenicity. Given a protein sequence *S* with a missense variant at residue position *i*, the LLR score can be computed using either the wild-type (wt), mutant (mut), or masked (msk) sequence. These three inputs differ only at residue *i*, where the masked marginal substitutes the original residue with a special mask token.

Let 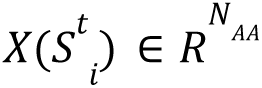 denote the model output vector (logits) for position *i* in sequence type *t* ∈ {*wt*, *mut*, *msk*}, where *N_AA_* is the number of amino acid tokens.

We define the LLR score as:*_i_*

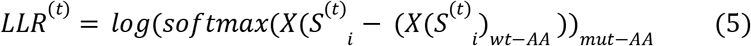

That is, we first subtract the wildtype entry from the entire output vector of residue *i*, apply a softmax to the resulting log-odds ratios, and finally take the mutant entry of this normalized vector as the LLR score.

We note that in some cases, prior works computed the LLR scores by first applying the *log_softmax* function and then subtracting the wildtype score. However, this approach mathematically cancels the softmax normalization, reducing the computation to the raw logit difference. This follows directly from the definition of the softmax function. Given an unnormalized logits vector *l* it holds that:

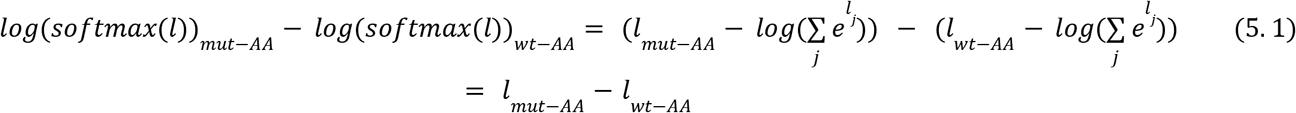

Although we did not observe substantial performance differences using this formulation, all reported comparisons in this study were performed using the implementation described in (5).

#### Extending LLR to long sequences

ESM1b has a maximum input context of 1,024 tokens, which limits the model’s ability to process full-length proteins exceeding this length. To handle such cases, we applied a sliding-window approach with a window size of 1,022 residues and a 250-residue overlap between consecutive windows. For each window, the model computes ESM1b scores for the central 1,022 − 2×250 positions, excluding the overlapping regions to avoid redundancy. In the first and last windows, the terminal residues are also included to ensure full sequence coverage. This strategy preserves the model’s maximal input length while preventing indexing errors and maintaining smooth coverage across long proteins. Following window extraction, LLR computation proceeds identically to the method described above, using the wildtype, mutant, or masked representations as appropriate.

### Classification Performance Metrics

#### Global and per-protein AUROC

The global and per-protein Area Under the Receiver Operating Characteristic curve (AUROC) are widely used metrics to assess VEP performance. For all models, we assume that higher scores indicate greater pathogenicity. This assumption holds for all cases where calibration using logistic regression was performed. When raw scores are analyzed (Figure 1a), we first learn a monotonic rescaling function using the modified min-max normalization described below and, if necessary, flip the scores so that values closer to 1 correspond to pathogenic variants and those closer to 0 to benign variants. Because this transformation is strictly monotonic, it preserves the global rank order of scores and thus does not affect the AUROC.

The per-protein AUROC was computed by first calculating the AUROC independently for each protein sequence and then averaging across all valid sequences. Proteins lacking either pathogenic or benign labels were excluded from the mean calculation. Consistent with the ProteinGym benchmark protocol, we report the unweighted mean of per-protein AUROCs. We also tested a weighted-average variant, in which each protein’s contribution was proportional to its number of annotated variants; however, the results were nearly identical to the unweighted mean, and thus only the standard approach is reported.

All AUROC computations were performed on held-out test sets, strictly excluding any samples used during the calibration process. When comparing RaCoon to the uncalibrated ESM1b model, we always evaluated performance on identical test sets to ensure a fair comparison. We note that RaCoon’s ablation experiments (Figure S12) involved multiple calibration settings, which altered the composition of the resulting test sets leading to slight variations in reported AUROC values across experiments.

### Residue Attributes

All attributes were directly inferred from the protein sequence associated with the variant. Predictors used as well as associated thresholds are specified in **Table 3**.

#### Disordered regions

Residues in disordered regions were inferred using ALBATROSS Metapredict v1.3^109,110^. We precomputed disorder scores per residue for all isoforms of reviewed human proteins in UniProt^111^, as well as for all transcripts in our datasets. Each residue was locally mapped to the corresponding disorder score based on its position within the matching transcript. Residues with scores above 0.7 were defined as disordered. Although the suggested threshold is 0.5, we found that a 0.7 cutoff better aligned with previous disorder estimates^44,109^, predicting approximately 28% of residues as disordered compared to 32% using the 0.5 threshold. Increasing the threshold mainly changes the predictions of residues in transition regions therefore, aiming for more rigorous disorder prediction, we use the 0.7 threshold.

#### Protein-protein interface

To predict residues involved in protein–protein interfaces (PPIs), we used PIONEER^112^. We obtain high-confidence interface predictions for all reviewed human UniProt sequences. PIONEER integrates binary predictions from three sources: the PIONEER model, experimentally resolved complexes from the Protein Data Bank (PDB)^113^, and homology models. A residue predicted to be involved in a PPI by any of these sources was considered PPI-positive. Predictions were available for 74% of variants in the ClinVar_HQ dataset, of which 9% were predicted to participate in PPIs (N = 11,558). In the ProteinGym dataset, predictions were obtained for 77% of variants, with 12% predicted to participate in PPIs (N = 5,749).

#### Homology

To distinguish between sequences with few versus many homologs, we directly queried the UniRef sequence clusters^114,115^. A key advantage of estimating homology using UniRef clusters is that it avoids systematic similarity searches across the entire human proteome, which would be impractical at inference time. We experimented with both UniRef90 and UniRef50 clusters, testing multiple thresholds to maximize separation of class-conditional distributions. We ultimately used UniRef90, defining sequences with ten or fewer homologs as low-homology. Using this threshold, approximately 9% and 7% of sequences in the ClinVar_HQ and ProteinGym datasets, respectively, were categorized as low-homology. While lower thresholds increased the class-conditional separation, they also produced very small subgroups. A threshold of nine therefore provided a suitable balance between subgroup size and distributional contrast.

#### Physico-chemical properties

We compiled an extensive set of amino acid physicochemical properties previously reported to influence the functional impact of missense variants (**Table** 3)^116–120^.

#### Optimal classification thresholds

Optimal classification thresholds were determined by maximizing the Youden J-statistic^121^ across all possible decision thresholds. That is, we chose the threshold that maximized the difference between the true positive rate and the false positive rate:

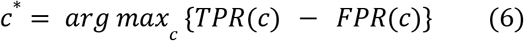

J-score is often preferred over accuracy in imbalanced test sets and, unlike the F1-score, it also accounts for true-negatives^122^. These make it particularly suitable for clinical and diagnostic tasks^123^ and it has also been adopted for VEP benchmarking^75^.

For the uncalibrated ESM1b model, the optimal threshold determined by maximizing the J-score on the ClinVar_HQ dataset was −8.22, which differs from the paper suggested threshold of −7.5. While the original rationale for the −7.5 threshold is not specified, using the J-score-derived threshold resulted in higher classification accuracy in our evaluation.

#### Mutual Information (MI) Estimation

To quantify the association between residue attributes and pathogenicity we computed the MI between selected attributes and the binary clinical label (**Appendix 2**, **Table S5**). At the residue level, MI was calculated directly from the binary attribute indicator and the pathogenic labels using scikit-learn’s *mutual_info_score*. At the protein level, each proteins’ residues were assigned to one of three groups, based on the proportion of residues exhibiting the attribute (corresponding to the violin bins in figure S3), MI was computed between these group assignments and the pathogenic labels of the residues. For both analyses, uncertainty was estimated using 1,000 non-parametric bootstrap iterations.

### Learning Gaussian Mixture Models Over LLR Scores

Gaussian Mixture Models (GMMs)^124^ are used in the study to fit the pathogenic and benign LLR distributions. We train all GMMs using the following scheme:

#### Extreme outliers removal (modified z-score)

We first removed extreme outliers from the full dataset (without separating pathogenic and benign variants) using the modified z-score^125,126^. For a set of variants *X* the modified z-score *z_i_* of a variant *x_i_* is defined as:

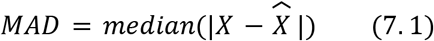

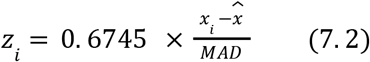

Where *X=median*(*X*). Samples with |*z_i_*| > 4. 25 were considered extreme outliers and excluded from GMM training. This approach removes outliers symmetrically from both tails of the distribution, effectively eliminating extreme benign and pathogenic scores. The chosen threshold corresponds to a highly stringent cutoff intended to exclude only extreme samples that could otherwise distort GMM fitting. For instance, across the full CalinVar_HQ dataset, this process excluded 576 variants corresponding to 0.33% of all entries.

#### Fitting GMMs

Following extreme outlier removal, we fit separate GMMs for the benign and pathogenic samples. GMMs were implemented using the GaussianMixture module from scikit-learn, configured with two components and diagonal covariance (equivalent to full covariance in one-dimensional data), while all other parameters were kept at default values.

The number of components was chosen by computing the Bayesian Information Criterion (BIC)^127^ across models with increasing component counts and selecting the point at which the BIC curve plateaued. This procedure minimizes the risk of overfitting while maintaining sufficient flexibility to capture multimodal score distributions.

#### Calibration with Logistic Regression (LR)

We used logistic regression to learn monotonic mappings for distinct residue-level attributes. All experiments followed a consistent sampling and fitting scheme, as outlined below.

#### Extreme outlier removal (percentile)

As per the GMM training protocol, we removed extreme outliers before fitting the logistic regression models. Specifically, samples falling in the top and bottom 0.1 percentiles (i.e., the most extreme 0.2% of values) were excluded. This proportion closely matches the number of outliers removed using the modified z-score method. However, because logistic regressions were fitted across multiple models with differing score ranges, we opted for a percentile-based criterion, which requires no model-specific adjustments (such as modifying the z-score threshold), ensuring consistent preprocessing across predictors.

#### Sampling

To mitigate majority-class bias^128,129^ we train all LR models on balanced datasets (either ClinVar_Balanced or the ProteinGym clinical substitution benchmark). To fit an LR model, we randomly sampled *n* random train variants leaving the rest for test. For Figure 1b we used *n* = 6, 000, for Figure 2a-c, we used *n* = 2000. For figure S8-9 the full dataset was partitioned into three disjoint subsets. In each iteration, one-third of the data was used for training, while the remaining two-thirds served as a test set. For the MCE,ECE comparisons of RaCoon against a naively-globally calibrated ESM1b we used 6,000 training samples.

#### Fitting logistic regression

Following outlier removal logistic regression (LR) models were fitted using the scikit-learn *LogisticRegression*. The model was trained with the Limited-memory Broyden–Fletcher–Goldfarb–Shannon (L-BFGS) solver^130^, using otherwise default parameters.

Given *x* the raw LLR score and *c*_1_, *c*_2_ the learned scale and intercept coefficient of the fitted LR the calibrated score is defined as:

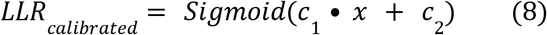

### Data Normalization

To analyze raw VEP scores within a unified reference frame (**Figure 1a**), we applied a modified version of the standard min-max normalization. First, extreme outliers, defined as values in the top and bottom 0.1 percentiles, were removed to prevent range distortion caused by outlier domination. Min-max normalization was then computed over the remaining inlier set. Following normalization, we verified that the mean score of pathogenic variants was higher that of benign variants, if not, the scores were flipped by subtracting them from 1. Finally, the excluded outliers were reinstated, assigning a score of 1 to pathogenic and 0 to benign variants. Scores that were already bounded within the [0, 1] range were not renormalized but were flipped if necessary. LRT scores were log-transformed as − *log*₁₀(*max*{*score*, 10⁻⁶}) before normalization.

### Jensen-Shannon divergence

To quantify divergence between class-conditional score distributions, we used the Jensen-Shannon divergence (JSD)^131^. JSD was empirically estimated by discretizing each subgroup’s LLR-scores into histograms with 100 equal-width bins.

To prevent zero-probability bins, a small smoothing factor ε=e^−12^ was added to all bins. JSD was then computed using the SciPy Jensen-Shannon implementation, the divergence is then obtained by squaring the Jenson-Shanon distance. The reported JSD value is calculated as the sum of the independently estimated JSDs between benign variants and between pathogenic variants across the two subgroups (equation 1, results):

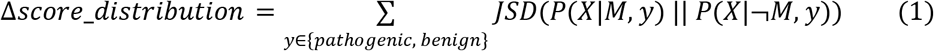

Where, *X* denotes the LLR-scores, *M* and ¬*M* partitions over residue attribute (e.g. polar vs. non-polar residues) and *y* - binary label (either pathogenic or benign).

To ensure JSD values are not affected by bin size, we computed the JSD for all attributes in and datasets analyzed in figure S7d using four bin resolutions (200, 100, 50, and 25 bins). Rank stability across bin sizes was assessed using Kendall’s τ, computed using the *SciPy kendalltau* implementation (**Table 4**). All pairwise comparisons showed strong positive rank correlations (τ ≈ 0.7-0.92), indicating that the relative ordering of attributes is highly robust to the choice of bin size.

**Table 4.** Bin size effect on JSD calculation. p-value two-sided test against τ = 0.

| Bin-size | Kendall $\tau$ | p-value |
| --- | --- | --- |
| 200 vs 100 | 0.857 | $p < 1.0e^{-12}$ |
| 200 vs 50 | 0.784 | $p < 1.0e^{-10}$ |
| 200 vs 25 | 0.721 | $p < 1.0e^{-9}$ |
| 100 vs 50 | 0.921 | $p < 1.0e^{-14}$ |
| 100 vs 25 | 0.851 | $p < 1.0e^{-12}$ |
| 50 vs 25 | 0.917 | $p < 1.0e^{-15}$ |

### Calibration Matrices

#### Reliability histograms

Calibration, or reliability histograms^76,77^, were used to visualize the reliability of the predicted model probabilities. Reliability histograms can only be applied to models that output probabilistic scores within the [0, 1] range. Histograms were constructed by discretizing model predictions into 10 equally-spaced bins. We then compute the per bin empirical pathogenic frequency. For a perfectly calibrated model, the average predicted confidence should match the observed frequency (up to binning artifacts), yielding points close to the identity line. Deviations above or below this line indicate overconfidence or underconfidence, respectively.

#### Expected Calibration Error

While reliability histograms provide intuitive visual insights, they are difficult to quantify and may be misleading when sample counts vary substantially across bins. To provide a quantitative measure of calibration, we computed the weighted Expected Calibration Error (ECE)^39^. For *M* bins *b*_1_,…,*b_M_* let *n_b_*, *freq_b_* and *conf_b_* denote, the empirical pathogenic frequency, and the mean predicted confidence in bin *b*, respectively and let *N* be the total number of variants.

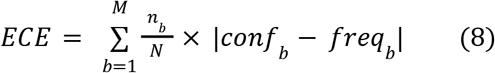

An ECE of 0 indicates perfect calibration, while higher values represent poorer calibration. Although ECE does not distinguish between overconfidence and underconfidence, these effects can be easily inferred from the accompanying reliability histograms.

#### Maximal Calibration Error

Reported as the maximal calibration error across all bins with > 50 sample:

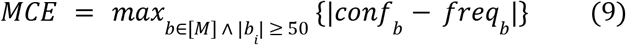

#### P-value and multiple hypothesis correction

For Table S4, p-values were computed using two-sided Fisher’s exact test and adjusted for multiple comparisons with the Benjamini-Hochberg procedure. For the mutual-information (MI) analysis (Figure S3, Table S5) p-values were calculated using the Mann-Whitney U test (SciPy’s *mannwhitneyu,* two-sided test) over 1,000 non-parametric bootstrap MI values per attribute at the residue vs the protein level. The p-value reported in Figure S3d-e,S8,S9 corresponds to the default two-tailed p-value from SciPy’s *pearson* method, calculated based on the t-distribution.

For the well-annotated, balanced protein analysis (Figure S14), we compared the AUROC of RaCoon with the raw ESM1b AUROC separately for each protein using a two-sided normal z-test. For a protein with *n* variants, the standard error was estimated as in Eq. 10.1, and the corresponding z-statistic was computed as in Eq. 10.2. Resulting p-values were corrected for multiple testing across proteins using the Benjamini-Hochberg procedure, and proteins with adjusted q-values below 0.05 were considered significant.

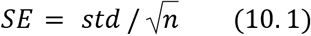

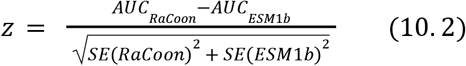

### The RaCoon Pipeline

#### Partitioning and Pruning

To construct the calibration tree, we first build a complete tree over all chosen binary residue attributes, where each leaf represents a distinct attribute combination. Because sparse attribute subsets are later pruned, the order of partitioning is important. We begin by dividing the data into residues from short and long proteins, and then iteratively split each node using the attribute that shows the most significant class-conditional distribution difference within that subset until all attributes have been exhausted (**Algorithm 1**, Methods). This greedy ordering prioritizes the most discriminative attributes, likely to improve AUROC, near the root and facilitates subsequent pruning by grouping nodes in ascending order of conditional distribution difference. After the initial tree is constructed, we iteratively prune leaves that do not meet minimum sample thresholds (**Algorithm 2**, Methods). These thresholds are determined by the number of samples required to fit each class-specific GMM and the need to retain held-out data for evaluation. Increasing the required training samples per GMM therefore raises the pruning thresholds and reduces the number of retained calibration subgroups (**Figure S12b**). Following hyperparameter tuning, we selected 400 training variants per GMM and required each retained leaf to contain at least 1,600 variants in total, including at least 400 pathogenic and 400 benign variants (**Figure S12**, Methods).

**Algorithm 1: Dataset partitioning**

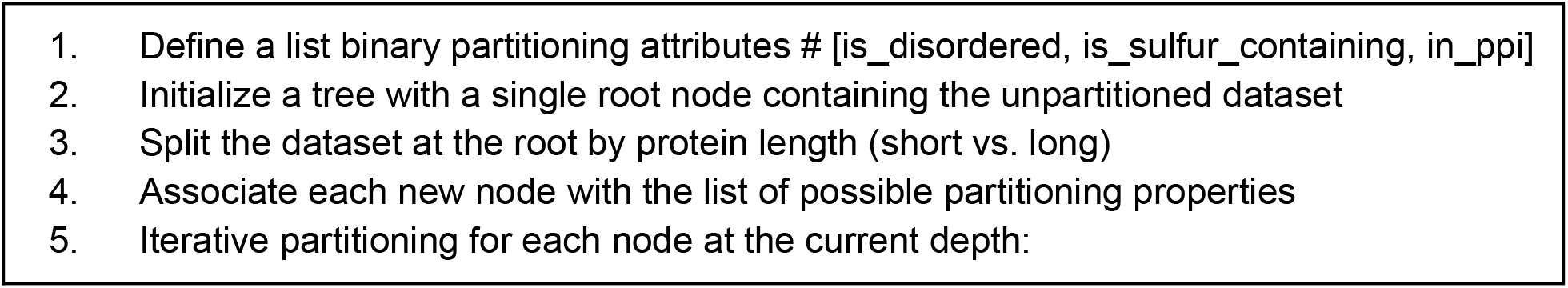
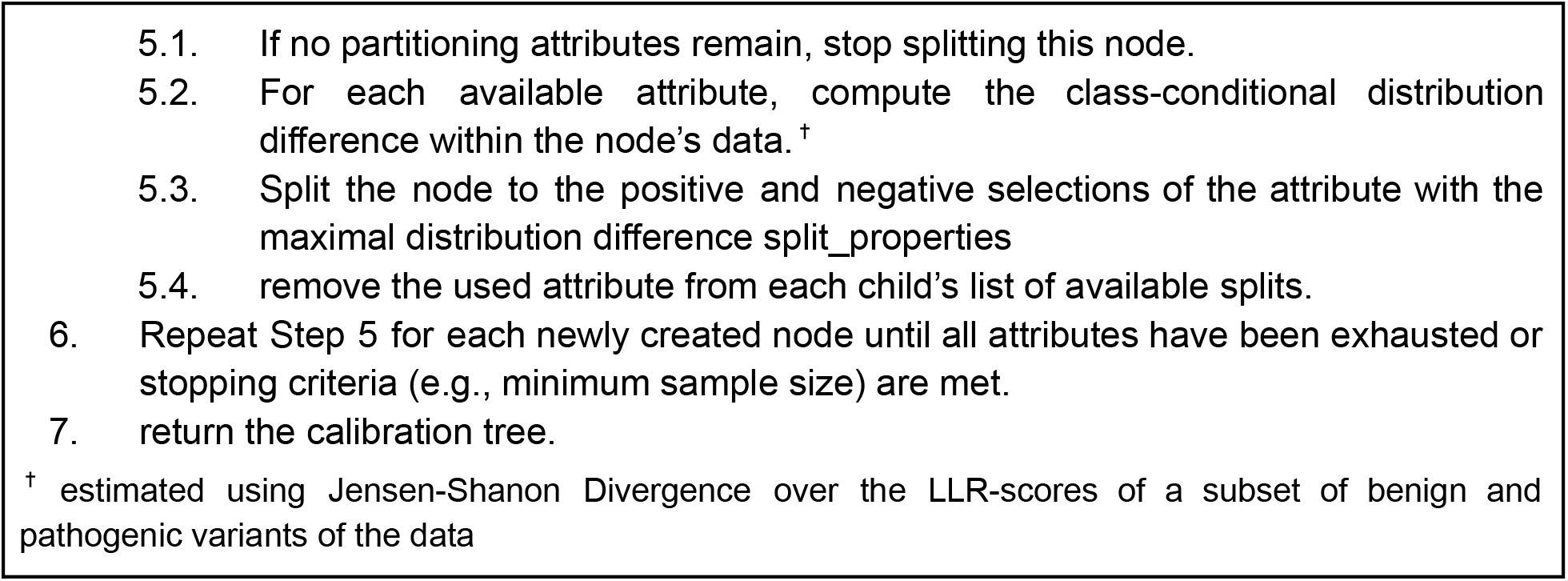

**Algorithm 2: Pruning**

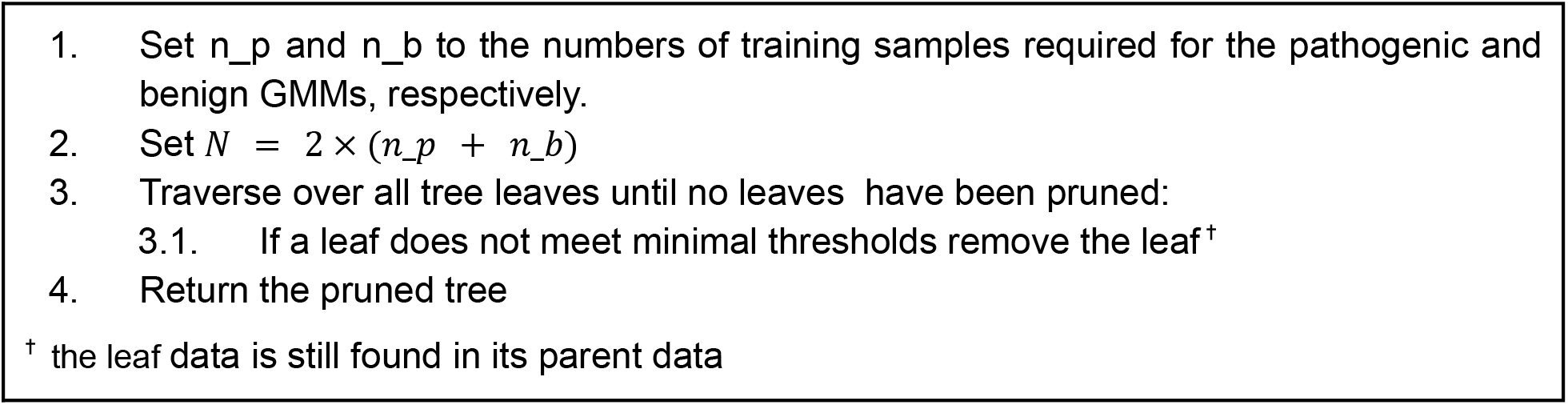

#### Modelling

Our calibration strategy relies on estimating the pathogenic and benign LLR score distributions rather than using explicitly labeled samples. A natural approach for modeling these distributions is to fit two GMMs for each node of the calibration tree, one for pathogenic and one for benign variants (**Algorithm 3, Methods**). We first estimate the fraction of pathogenic variants within each node using 500 randomly sampled variants (**Methods**). To reduce training size, the same variants are also used for GMM fitting. Each GMM is trained on 400 pathogenic and 400 benign variants, which we find yields results surpassing those obtained with tens of thousands of labeled examples (**Figure S12a-b**). Synthetic samples generated from the fitted GMMs, sampled according to the estimated pathogenic fraction, are then used to construct calibration histograms that map LLR values to pathogenicity probabilities.

The GMM-based strategy offers four key advantages. First, it effectively decouples calibration from the size of the node’s dataset, with only ∼800 samples needed to fit the model (**Figure S12a-b**). Second, our hyperparameter choices intentionally constrain the expressiveness of the GMMs by limiting the number of fitted parameters. This reduces the risk of overfitting, preventing the model from capturing noise or outliers. Third, as generative models, GMMs enable unlimited synthetic sampling, facilitating the construction of finer-grained calibration histograms and thus improving scores interpretability. Importantly, drawing additional samples may, at most, lead to overfitting of the GMM distribution. Finally, drawing from GMMs prevents the calibration process from direct exposure to labeled examples, maintaining minimal downstream supervision and avoiding memorization of specific variants used in calibration. In fact, even when GMMs are trained on the full node dataset, intentionally introducing potential leakage, no measurable improvement in AUROC is observed (Appendix 3).

**Algorithm 3: Fitting GMMs**

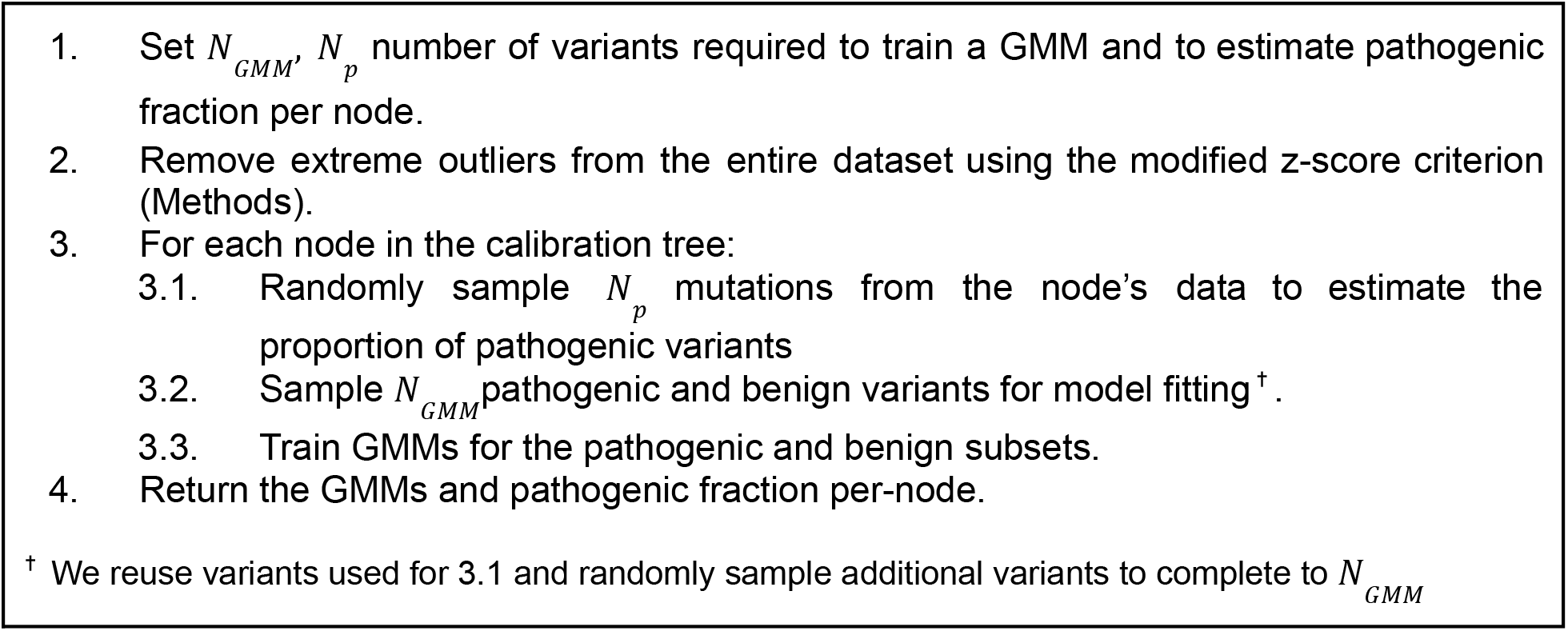

#### Binning

To translate LLR scores into meaningful pathogenicity estimates, we construct calibration histograms with equal-frequency bins where each bin represents a specific range of LLR values (**Algorithm 4, Methods**). As LLR scores are not uniformly distributed, using equal-frequency binning inherently adjusts bins granularity to match the underlying data density. Each bin is then assigned a pathogenic score given by the empirical proportion of pathogenic variants within that bin. In practice, we draw 40,000 synthetic samples from the fitted GMMs, sampling from the pathogenic and benign components according to the estimated pathogenic to benign ratio in the node’s data. We perform an extensive hyperparameter search over the number of bins and synthetic samples, and find that 40,000 synthetic samples and 50 bins yield optimal AUROC performance while producing finely grained and interpretable scores (**Figure S12c-d, Methods**).

**Algorithm 4: Binning**

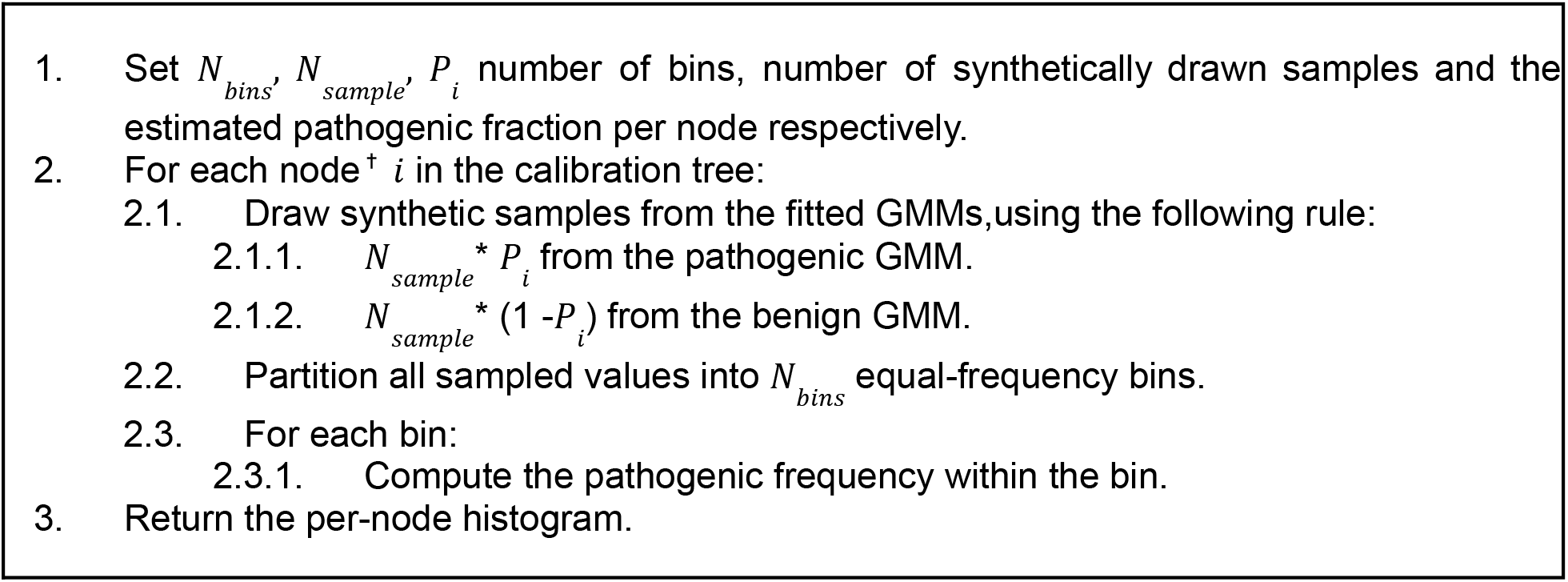
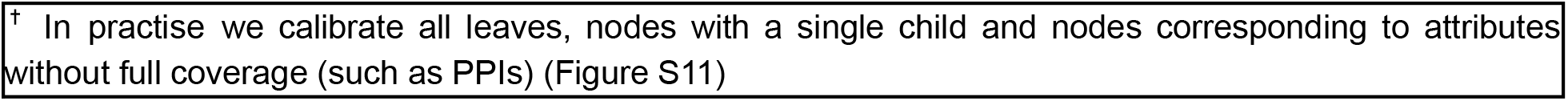

#### Mapping

At inference time, a new variant is first assigned to its corresponding node in the calibration tree. Variants are mapped to leaf nodes unless the matching child was pruned or, in edge cases, the next attribute annotation is missing, in which case they are mapped to the deepest reachable non-leaf node. The 1,600-variant pruning threshold applies only to leaves. All calibrated non-leaf nodes have sufficient data for GMM fitting, although some have insufficient held-out support for ECE/MCE reporting. The variant’s raw LLR score is then mapped to the appropriate bin, from which the corresponding pathogenic fraction is retrieved **(Algorithm 5, Methods).** This procedure produces calibrated, interpretable predictions that directly translate model outputs into clinically meaningful frequencies.

**Algorithm-5 Mapping:**

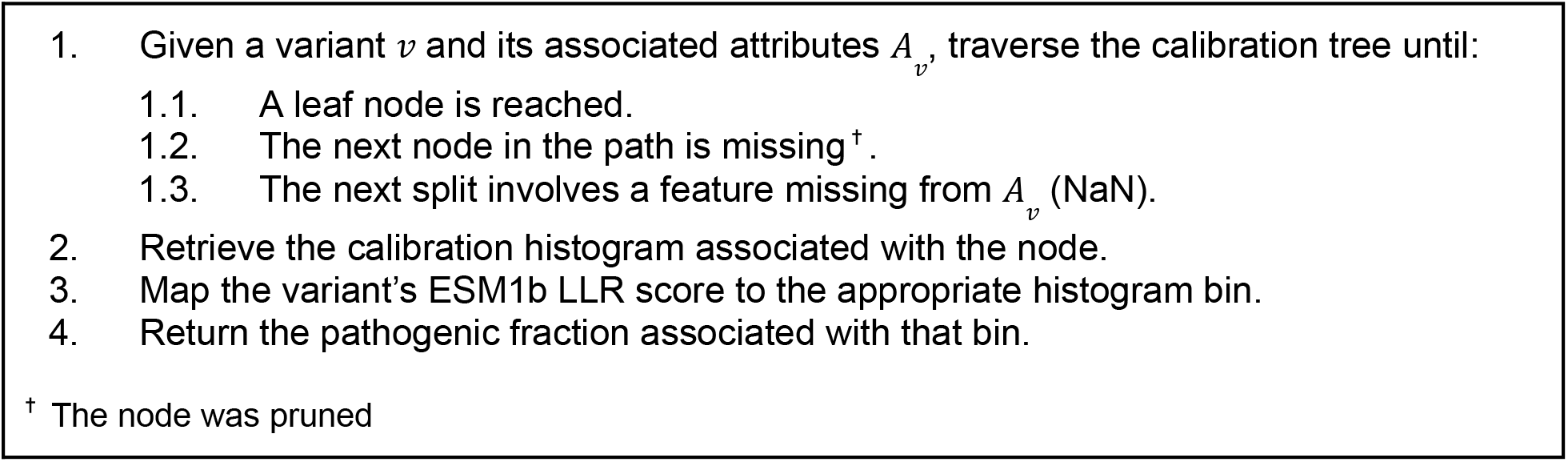

### Hyperparameters tuning

#### GMM training sample size

To isolate the effect of training set size on calibration performance we trained pathogenic and benign GMMs per-node with an increasing number of samples. We used a simplified calibration tree, partitioning only on length and residue disorder status (ordered versus disordered), both having 100% coverage and proven to improve AUROC on ESM1b throughout the study. This step was essential because pruning thresholds are set according to the required training samples per GMM. Increasing the training size can therefore lead to additional pruning, confounding the effects of tree composition and training size on overall performance. The simplified tree retained four calibration subgroups with sufficient data across all training sizes tested, keeping tree composition fixed. To construct calibration histograms we fixed the number of synthetically drawn samples at 40,000 and the number of bins at 50. Reported results were obtained using the ClinVar_Balanced dataset and represent the average over 100 independent training iterations, each consisting of 100 non-parametric bootstrap test iterations. Global AUROC was computed on an unseen test set composed of the remaining samples. For the logistic-regression comparison, we fitted one model per calibration node using 2*n* randomly sampled variants, matching the combined training size of the pathogenic and benign GMMs (*n* variants per class). The calibration-tree structure and evaluation procedure were unchanged (**Figure S12b**). We also report the effect of training sample size on the number of retained calibration subgroups using the full set of partitioning attributes (**Figure S12**b). We set RaCoon’s threshold at 400 training samples per GMM as it achieved significantly superior AUROC over the naive model while maintaining a large tree size.

Next, we compared the GMM-based calibration to direct histogram binning using the same simplified tree structure and hyperparameters. We progressively bin an increasing number of raw LLR, and evaluated AUROC on the remaining unseen samples. Direct binning yielded inferior results than the GMM approach by a significant margin even when using thousands of training samples. Furthermore, sampling over 2,000 scores did not improve performance, and may have even reduced it (**Figure S12a**).

Finally, we tested whether a hybrid strategy combining direct binning with synthetic sampling could further improve calibration. Using the same setup, we trained per-node GMMs on 400 samples, generated 40,000 synthetic samples and progressively combined them with increasing numbers of raw scores to construct the calibration histograms. The hybrid strategy only hindered the performance of the pure GMM approach, demonstrating degrading results with increased raw samples size (**Figure S12a-b**). We hypothesize that without incorporating denoising procedures, raw samples have a low signal to noise ratio requiring very large train size to reach comparable performance.

#### Estimation of pathogenic frequency

To estimate the pathogenic frequency, we used 500 randomly sampled labels, which could also serve for GMM training to maintain a minimal training size. The labels were modeled as Bernoulli-distributed variables (1 pathogenic, 0 benign), and the sample size was selected to ensure an estimated pathogenic frequency *p* achieves a 95% confidence interval within ±5% margin. Under the normal approximation, a rigorous estimation, without assuming any prior on *p*, and without finite-population correction requires *n* = 385 samples, we increased this to 500 to provide a conservative margin.

#### GMM-generated samples and number of bins

Having established the minimum number of variants required for GMM training and the superiority of synthetic sampling over direct binning, we next optimized the number of synthetic samples drawn from the GMMs and the number of bins used in the calibration histograms. We first computed the full calibration tree on the ClinVar_Balanced dataset using a minimum of 1600 total variants per node and the established thresholds of 400 benign and 400 pathogenic variants (Algorithms 1-3). All experiments were conducted over 100 independent GMM training iterations, each averaged over 100 non-parametric bootstrap test iterations.

To determine the optimal number of bins, we drew 100,000 synthetic samples from the GMMs and constructed calibration histograms with an increasing number of equal-frequency bins (**Figure S12c**). Drawing 100,000 samples ensured sufficient coverage without constraining bin granularity. Performance plateaued at approximately 40-50 bins, with no further improvement observed beyond this range. Increasing the number of bins above 50 may create an artificial impression of better calibration, as the apparent gain reflects over-splitting of existing bins rather than genuinely improved estimation of fine-grained frequencies. We therefore set the number of bins to 50, this provides a high resolution that accurately reflects the model’s learned distributions.

Next, we fixed the number of bins at 50 and analyzed AUROC as a function of the number of synthetic samples drawn from the GMMs (**Figure S12d**). Following Algorithm 4, samples were drawn according to the pathogenic-to-benign ratio of each node’s fitted GMMs. We found that performance plateaued at 40,000 synthetic samples per leaf. Fewer samples likely failed to estimate per-bin pathogenic probabilities accurately, given the relatively high number of bins, while larger sample sizes neither improved nor degraded performance. This result supports our earlier claim that while the calibration process can overfit the learned GMM distribution, the GMM itself acts as a regularizer, preventing overfitting to the raw LLR distribution. Consequently, we set 40,000 synthetic samples per node as the default for all subsequent experiments.

#### Permutation analysis of protein-level label-fraction bias

To perform the protein-level label-fraction permutation analysis, proteins were divided into three groups by the number of annotated variants: up to 20, 21-60, and more than 60 variants. Within each group, pathogenic fractions were randomly reassigned across proteins using a derangement, such that no protein retained its original pathogenic fraction. For each protein, we determined the number of benign and pathogenic variants to sample by iterating over all feasible total sample sizes and selecting the pathogenic/benign count combination that minimized the absolute error from the assigned pathogenic fraction, retaining the largest sample size in case of ties. Variants were then sampled without replacement according to these counts and pooled across proteins to form a permuted dataset. This procedure retained approximately 50% of variants and was repeated for 1,000 permutations. For each calibration setting, RaCoon was fitted once on the original, unpermuted calibration dataset and applied to all permutations without refitting.

#### Data leakage experiment

A control analysis showing that RaCoon does not gain performance from exposure to specific calibration examples is provided in Appendix 3.

#### ACMG/AMP PP3/BP4 evidence mapping

RaCoon-calibrated outputs were mapped to PP3/BP4 evidence strengths using the established clinical evidence-calibration framework developed by Pejaver et al. (2022) and Bergquist et al. (2025). To maintain independence between RaCoon model calibration and subsequent clinical evidence calibration, all variants present in the datasets used by the PP3/BP4 framework (ClinVar2019, ClinVar2020, or gnomAD) were excluded from the ClinVar cohort used for RaCoon calibration. The resulting RaCoon model was therefore disjoint from the datasets used to derive and evaluate the ACMG/AMP evidence mapping. records representing noncanonical amino-acid substitutions or whose protein-level annotations did not match the corresponding reference transcript were excluded, and duplicate protein-level substitutions were collapsed into a single record. Together, these filtering steps removed less than 2% of records across the three datasets.

Following the established framework, local posterior probabilities were estimated across the range of RaCoon-calibrated scores using ClinVar2019 and an adaptive sliding window containing at least 100 ClinVar2019 variants and 3% rare gnomAD variants. Benign observations were weighted using an estimated pathogenicity prior of (0.0441) to account for the enrichment of pathogenic variants in ClinVar. Sampling uncertainty was assessed using 1,000 bootstrap resamples, and evidence thresholds were defined using the conservative one-sided 95% confidence bound and the posterior-probability requirements corresponding to Supporting, Moderate, 3-point, and Strong PP3/BP4 evidence. Scores that did not reach supporting evidence in either direction were assigned to an indeterminate interval. The resulting global evidence mapping is provided in Table S8.

### RaCoon server version

RaCoon is made publicly available through an open-access server. The server version returns predictions averaged over the median of 75 out of 100 randomly initialized RaCoon models trained on the ClinVar_HQ dataset, allowing standard deviations to be reported for each prediction. In addition, node-specific ECE and MCE values are provided. Support for node-specific classification thresholds will be incorporated in a future release.

## Data Availability

The curated ClinVar-derived datasets and test set are available through Zenodo (record id 21914832). Source data underlying all figures and evaluations are available through Zenodo (record id 23765971). The original ClinVar GRCh38 records are publicly available from ClinVar, and the ProteinGym Clinical Substitution Benchmark is publicly available at https://proteingym.org. Other third-party datasets and predictor scores used in this study are publicly available from the sources cited in the Methods. No controlled-access data was used.

## Code Availability

The RaCoon source code and the custom scripts used for data processing, calibration and analysis are publicly available at the RaCoon GitHub repository.

