## Supplementary text, figures and tables for "Calibrated Variant Effect Prediction at the Residue Level Using Conditional Score Distributions"

##### Appendix 1 - Comparing subgroup-difference metrics for calibration target selection

This appendix provides a more detailed comparison of subgroup-difference metrics used to evaluate potential calibration targets and predict AUROC gains following the differential calibration protocol. Using ESM1b as a representative model, we consider both label-based metrics and LLR score-based metrics.

To quantify score-distribution differences our primary metric utilizes the Jensen–Shannon divergence (JSD)[1], which measures differences between continuous distributions - providing a natural approach to compare score distributions. For a given attribute $M$ and its complement $\neg M$, we compute the JSD between the corresponding LLR distributions separately for benign and pathogenic variants, and sum the two terms. Splitting the distributions by label reflects the fact that benign and pathogenic variants occupy different regions of the score axis (**Figure S4-5**). Formally:

$$\Delta score\_distribution = \sum_{l \in \{pathogenic,\ benign\}} JSD(P(X|M, l)\ ||\ P(X|\neg M, l)) \qquad (1)$$

Here, $X$ denotes the distribution of LLR values. For example, for disordered residues ($M$) versus ordered residues ($\neg M$). LLR scores can be computed from the wild-type sequence, the mutant sequence, or a masked sequence in which the mutated position is replaced by a mask token (Methods).

We compare the class-conditional JSD metric with two simpler alternatives. First, we evaluate a simpler ,label-agnostic,. score-based baseline defined as the absolute difference between the mean model prediction in the full dataset $E(X)$ and in the subgroup $E(X|M)$ (Equation 2).

$$\Delta mean\_prediction = |E(X)\ -\ E(X|M)| \qquad (2)$$

Second, we evaluate a purely label-based alternative, defined as the absolute difference between the global pathogenic fraction and the pathogenic fraction within a given subgroup (Equation 3, Appendix 1):

$$\Delta label\_fraction = |P(Y = pathogenic)\ -\ P(Y = pathogenic\ |\ M)| \qquad (3)$$

We first evaluate the correlation between changes in score distributions and changes in label fraction across three partitions of variants: substitutions at the wild-type amino acid (**Figure S7a**), substitutions to specific amino acids (**Figure S7b**), and variants involving residue-level attributes (**Figure S7c**). In the context of miscalibration, this analysis is important because changes in pathogenic fraction alter the subgroup pathogenicity prior and can therefore contribute to subgroup-specific miscalibration, as further discussed in Appendix 2. Each class is evaluated using the input representation required to capture its signal: wild-type sequences for substitutions at the wild-type amino acid, mutant sequences for substitutions to specific target amino acids, and masked sequences for residue-level attributes. Mutant sequences are essential for the target-amino-acid partition because only this representation encodes the introduced residue. Across all three, pronounced changes in pathogenic fraction can occur without corresponding changes in score distribution, likely reflecting labeling biases or limited model capacity, whereas the reverse is rarely observed, highlighting score-distribution differences as a more robust metric for evaluating calibration-target selection

We next examine the predictive capacity of the three approaches of AUROC gains following differential calibration. Class-conditional JSD shows the strongest association with calibration-induced AUROC gains ($r\ = 0.837,\ \ p\ < 1e - 24$), whereas mean-score differences show a weaker association ($r\ = 0.675,$ $p\ < 1e - 12$) and pathogenic-fraction differences are substantially less predictive (**Figure S8**, Table S3). This indicates that calibration target selection benefits from using the full score distributions, rather than only average score shifts or label-frequency differences. To control for differences in subgroup size, we also report a normalized AUROC contribution metric (**Figure S9**). The normalized metric shows an even slightly stronger correlation with JSD ($r\ = 0.863,\ \ p\ < 1e - 27$) .

We also compare JSD and ECE as discussed in the main text (**Figure 3d-e**), where we show that score-distribution differences better match AUROC gains after differential score mapping.

This analysis translates into measurable gains for specific residue attributes. In ESM1b, disorder level and protein–protein interface status yield the largest AUROC gains when calibrated separately under masked representations (up to ~0.01 and ~0.004, respectively), while aromatic and sulfur-containing residues show smaller but consistent improvements (up to ~0.003) under wild-type representations.

#### Appendix 2 - Pathogenic-fraction differences and their relation to score distributions

Two key types of distributional shift relevant to calibration are label shift[2–4], and score distribution shift[3,5–7]. These correspond to changes in the proportion of pathogenic variants $P(Y|Mi)$ and in the distribution of model scores $P(X|Mi)$ within a subgroup, respectively. The latter is the focus of the main text. As previously defined $Mi$ denotes a residue-level, typically represented as a binary partition (e.g., disordered vs. ordered residues). In this section, we examine the complementary label-level signal.

Changes in the proportion of pathogenic variants at the residue level have been previously observed across specific attributes such as disorder level[8–10], physicochemical properties[11], and solvent accessibility[12]. We extend these observations using high-quality clinical annotations from ClinVar (ClinVar_HQ; N = 171,196) and experimental data from the ProteinGym clinical substitution benchmark (N = 62,727). We define significant changes as residue subgroups in which the pathogenic fraction significantly deviates from the global baseline (**Table S4, Extended Table S4**).

Across both datasets, fewer pathogenic variants are found in intrinsically disordered regions and in sequences with few homologs, whereas pathogenic variants are more abundant in polar residues, sulfur-containing residues, and protein-protein interfaces (PPIs). The trends are consistent across both datasets, suggesting that they may capture biological signals beyond dataset-specific annotation biases. Excluding N-terminal methionine (M1) variants from ClinVar preserves these trends, with the largest decrease in pathogenic fraction among sulfur-containing residues (55.0% to 49.5%), but they remain strongly enriched relative to the revised dataset baseline of 30.2% (**Extended Table S4**).

The relationship between pathogenic-fraction differences and score-distribution differences varies across attributes. The lower pathogenic fraction in disordered regions likely reflects their increased mutational tolerance[13,14] and is well captured by ESM1b LLR-scores (**Figures S4-5**). In contrast, PPI residues show one of the strongest pathogenic-fraction differences but more modest score-distribution differences (**Figures S4-5**), suggesting that sequence-only models capture these effects only partially. This is consistent with studies showing improved PPI prediction when structural or contextual information is incorporated[15–17]. Likewise, the limited score-distribution differences observed in sequences with few homologs may reflect reduced model generalization or annotation biases (**Figures S4**).

Because VEP calibration is often performed at the protein level, we also examine whether residue-level trends are also reflected at the protein level. We analyze a set of well-annotated ClinVar proteins (N = 1,285; Methods), grouping proteins into three bins based on the fraction of residues exhibiting a given attribute. We then compare the distribution of pathogenic variants across these bins (**Figure S10**). We focus on disordered and interface residues, which display the most extreme changes in pathogenic fraction, and on homology, which displays a more subtle change.

Similar trends are observed at the protein level. For example, pathogenic variants are less frequent in highly disordered proteins and more frequent in proteins enriched for PPIs. These patterns suggest that residue-level effects contribute to variability observed at the protein level and help explain why protein-level calibration can capture meaningful signal[18–21]. At the same time, this analysis considers only residue-derived attributes and does not address additional sources of variation, such as inheritance patterns or gene-specific effects, that can only be captured at the protein level.

However, the analysis also reveals that the protein-level signal is noisier and less informative. The mutual information (MI) between disorder status and pathogenicity is markedly higher at the residue level (MI =

0.051) than at the protein level (MI = 0.014), with protein-level estimates exhibiting an order-of-magnitude higher variability (**Table S5**, Methods). These results support residue-level calibration as a complementary approach for capturing variation that may be obscured at the protein level.

Importantly, differences in pathogenic fraction alone are insufficient to identify appropriate calibration targets. While they may reflect biological effects, they may also arise from experimental or annotation biases. Effective calibration requires that subgroup differences are also captured by the model's predictions[3,22] which is better assessed through class-conditional score distributions motivating their use as our main approach. When pathogenic-fraction differences reflect biological signals captured by the model, they are expected to be reflected in score-distribution differences[3,5–7]. When they are not reflected in model scores, they are less likely to indicate useful calibration targets.

#### Appendix 3 - RaCoon data leakage ablation

To test whether RaCoon gains performance by memorizing specific calibration examples, we compare its performance on variants seen during calibration with its performance on unseen variants. For each node, we first create a balanced set of benign and pathogenic variants by downsampling the larger label class to match the smaller one. This balanced set is then split into equally sized, label-balanced calibration and test subsets. The calibration subset is used to fit the per-node GMMs. We then evaluate RaCoon either on the held-out test subset, representing the no-leakage setting, or on the same calibration subset used to fit the GMMs, representing deliberate data leakage. Because the two evaluation sets are matched in size and label fraction, differences in performance reflect exposure to the calibration examples rather than differences in class balance.

On ClinVar_Balanced, RaCoon achieves an AUROC of 0.889 ± 0.002 in both settings. On ProteinGym, RaCoon achieves an AUROC of 0.893 ± 0.002 without data leakage and 0.894 ± 0.002 with deliberate data leakage. Thus, even when evaluated directly on the variants used during calibration, RaCoon does not gain meaningful AUROC performance. This supports the conclusion that RaCoon's AUROC improvement is not driven by memorization of specific calibration samples. These AUROC values differ slightly from those reported in the main text because this experiment uses evaluation sets specifically constructed to equalize class distributions across all nodes.

#### Supplementary Tables

**Table S1. Classification thresholds and AUROC across residue subgroups in ClinVar_BM.**
This table is provided as a separate supplementary file.

**Table S2. Expected calibration error across residue attributes and predictors in ClinVar and ProteinGym.**
This table is provided as a separate supplementary file.

**Table S3. subgroup-difference metrics for calibration target selection.**
This table is provided as a separate supplementary file.

| Residue Attribute | Pathogenic Fraction | | Pathogenic Variants | | Benign Variants | | p-value[a] | |
|---|---|---|---|---|---|---|---|---|
| | ClinVar | ProteinGym | ClinVar | ProteinGym | ClinVar | ProteinGym | ClinVar | ProteinGym |
| Entire dataset | 0.31 | 0.51 | 52,634 | 32,000 | 118,562 | 30,727 | Baseline | |
| Disordered | 0.10 | 0.19 | 5,700 | 3,038 | 48,845 | 12,742 | $p \ll 1.0e-23$ | $p \ll 1.0e-23$ |
| Beta branched[b] | 0.18 | 0.36 | 5,414 | 3,495 | 24,777 | 6,104 | $p \ll 1.0e-23$ | $p \ll 1.0e-23$ |
| Low homology[c] | 0.19 | 0.42 | 2,361 | 1,503 | 10,108 | 2,068 | $p \ll 1.0e-23$ | $p \ll 1.0e-23$ |
| Hydrophobic | 0.28 | 0.48 | 16,173 | 9,419 | 40,674 | 10,287 | $p < 1.0e-23$ | $p < 1.0e-14$ |
| Polar | 0.37 | 0.57 | 20,388 | 12,004 | 34,797 | 8,907 | $p \ll 1.0e-23$ | $p \ll 1.0e-23$ |
| Helix Breakers[d] | 0.40 | 0.60 | 11,120 | 6,936 | 16,648 | 4,666 | $p \ll 1.0e-23$ | $p \ll 1.0e-23$ |
| Ordered | 0.40 | 0.62 | 46,934 | 28,962 | 69,717 | 17,985 | $p \ll 1.0e-23$ | $p \ll 1.0e-23$ |
| Aromatic | 0.45 | 0.68 | 5,156 | 3,244 | 6,333 | 1,557 | $p \ll 1.0e-23$ | $p \ll 1.0e-23$ |
| Sulfur Containing[e] | 0.55 | 0.66 | 6,277 | 2,438 | 5,146 | 1,255 | $p \ll 1.0e-23$ | $p \ll 1.0e-23$ |
| PPI | 0.61 | 0.73 | 6,931 | 4,376 | 4,445 | 1,659 | $p \ll 1.0e-23$ | $p \ll 1.0e-23$ |

**Table S4. Pathogenic fraction across selected residue subgroups in ClinVar_HQ and ProteinGym .**

ProteinGym - clinical substitution benchmark [a] multiple hypothesis correction applied (FDR), capped at 1.0e-23 [b] Valine, Isoleucine, Threonine. [c] sequences with Uniref90 cluster size < 10 (Methods). [d] Proline Or Glycine. [e] Methionine or Cysteine.

**Table S4_Extended.**
This table is provided as a separate supplementary file.

| Residue Attribute | Residue-level MI | | Protein-level MI | | p-value |
|---|---|---|---|---|---|
| | mean | std | Mean | std | |
| Disorder level | 0.051 | $6.88 \times 10^{-4}$ | 0.014 | $6.16 \times 10^{-3}$ | p≪1.0e-22 |
| Homology count | 0.014 | $4.75 \times 10^{-4}$ | 0.005 | $4.73 \times 10^{-3}$ | p= 0.21 |
| Interface status | 0.0002 | $2.00 \times 10^{-4}$ | 0.016 | $6.39 \times 10^{-3}$ | p=0.99 |

**Table S5. Mutual information between residue attributes and pathogenicity at the residue and protein levels.** p-value calculated using Mann–Whitney U-test (two sided) for residue-level MI > protein-level MI.

| Calibration Set | Test Set | # Test Variants | AUROC ESM1b | AUROC RaCoon | ΔAUROC | RaCoon > ESM1b[a] |
|---|---|---|---|---|---|---|
| ClinVar_Balanced | ProteinGym | 23,478 | 0.904±0.003 | 0.907±0.002 | 0.003±0.001 | 100% |
| ClinVar_Balanced | Balanced Protein Subset (ClinVar) | 6,137[b] | 0.916±0.004 | 0.919±0.005 | 0.003±0.002 | 94.0% |
| ClinVar_Balanced | ClinVar_Balanced | 23,408[b] | 0.904±0.003 | 0.907±0.003 | 0.002±0.001 | 100% |
| ProteinGym | ProteinGym | 26,713[b] | 0.905±0.003 | 0.908±0.004 | 0.003±0.002 | 94.1% |

**Table S6. Protein label-ratio permutation analysis across RaCoon evaluation settings.**

[a]Fraction of permutations where RaCoon outperformed ESM-1b [b]Examples used in RaCoon's calibration excluded from test.

**Table S7. Calibration error across RaCoon calibration-tree nodes.**
This table is provided as a separate supplementary file.

| Evidence category | Score interval (ClinVar 2019) | Validation LR (ClinVar 2020) | # Benign | # Pathogenic | GnomAD variants (%) |
|---|---|---|---|---|---|
| **BP4 strong** | – | – | – | – | – |
| **BP4 3-point** | $\leq 0.500$ | $0.02$ | $219$ | $2$ | $2.45$ |
| **BP4 Moderate** | $(0.500, 4.125]$ | $0.09$ | $2491$ | $103$ | $31.03$ |
| **BP4 Supporting** | $(4.125, 14.375]$ | $0.20$ | $1240$ | $112$ | $19.75$ |
| **Indeterminate** | $(14.375, 60.125)$ | – | – | – | $28.94$ |
| **PP3 Supporting** | $[60.125, 77.500)$ | $2.35$ | $275$ | $291$ | $6.38$ |
| **PP3 Moderate** | $[77.500, 90.750)$ | $6.30$ | $180$ | $510$ | $6.22$ |
| **PP3 3-point** | $[90.750, 98.750)$ | $22.46$ | $98$ | $990$ | $4.71$ |
| **PP3 strong** | $\geq 98.750$ | $424.58$ | $1$ | $191$ | $0.52$ |

**Table S8. Global mapping of RaCoon scores to ACMG/AMP PP3/BP4 evidence strengths.**

ClinVar2019 - pathogenic/likely pathogenic and benign/likely benign missense variants from the December 2019 ClinVar release; ClinVar2020 - missense variants added by the December 2020 release and absent from ClinVar2019; gnomAD - rare population variants (allele frequency <1%) from gnomAD. LR denotes the likelihood ratio between pathogenic and benign variants. # Benign and # Pathogenic indicate the respective numbers of variants in the ClinVar2020 validation dataset. An em dash indicates that the corresponding evidence strength was not reached or irrelevant.

### Supplementary Figures

**Figure S1.** Threshold shifts across residue subgroups

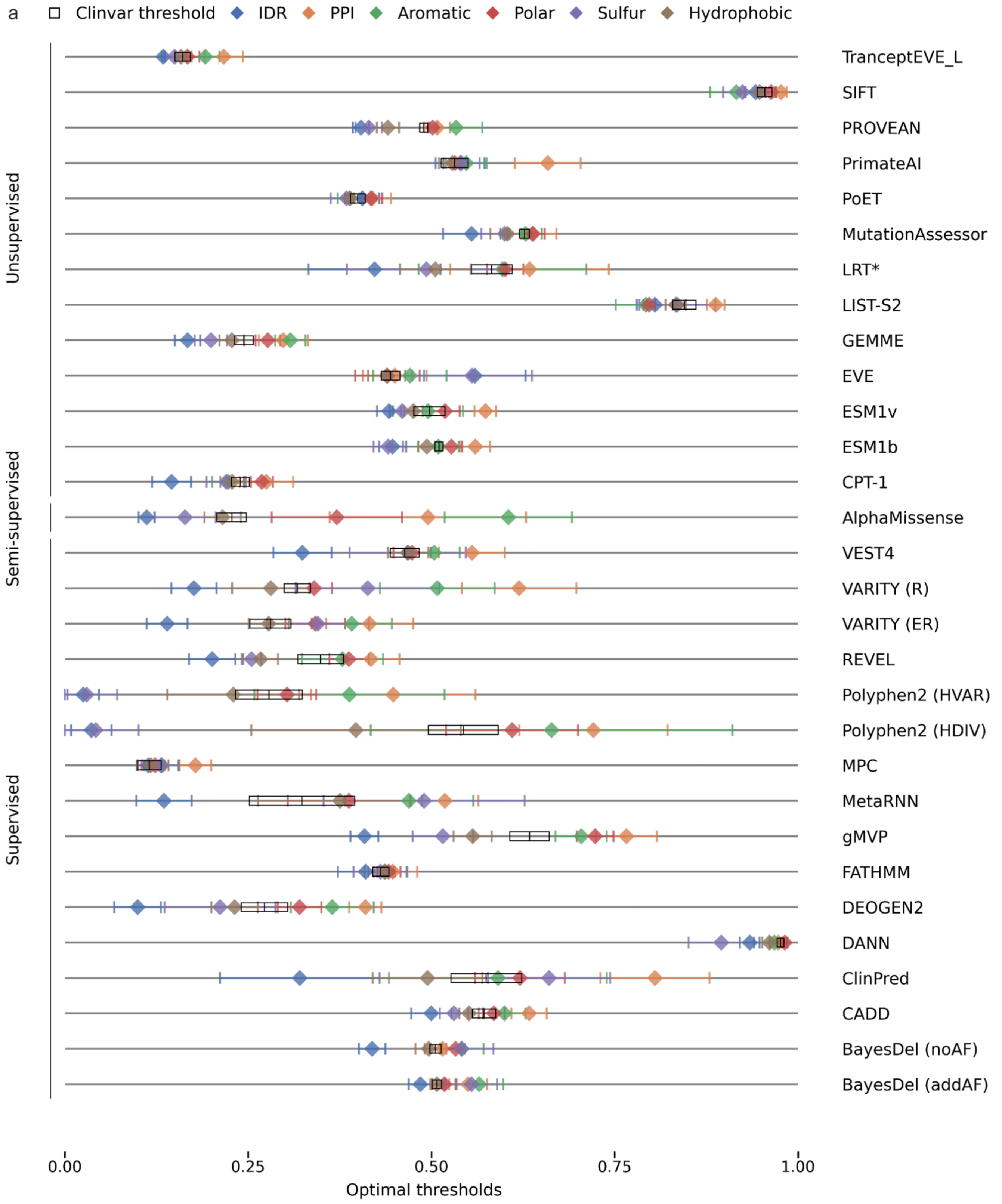

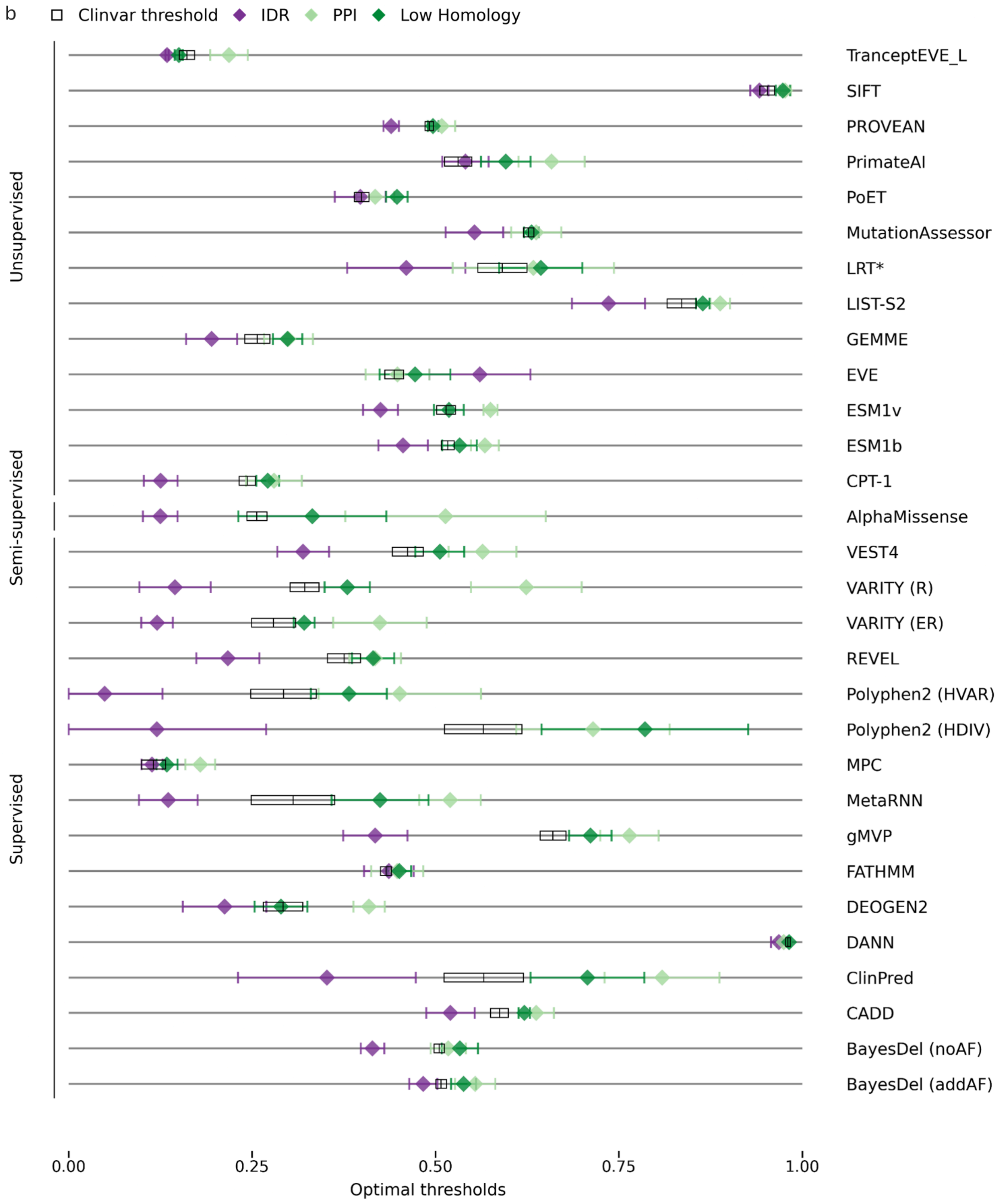
b
Clinvar threshold
IDR
PPI
Low Homology
Unsupervised
Semi-supervised
Supervised
TranceptEVE_L
SIFT
PROVEAN
PrimateAI
PoET
MutationAssessor
LRT*
LIST-S2
GEMME
EVE
ESM1v
ESM1b
CPT-1
AlphaMissense
VEST4
VARITY (R)
VARITY (ER)
REVEL
Polyphen2 (HVAR)
Polyphen2 (HDIV)
MPC
MetaRNN
gMVP
FATHMM
DEOGEN2
DANN
ClinPred
CADD
BayesDel (noAF)
BayesDel (addAF)
0.00
0.25
0.50
0.75
1.00
Optimal thresholds

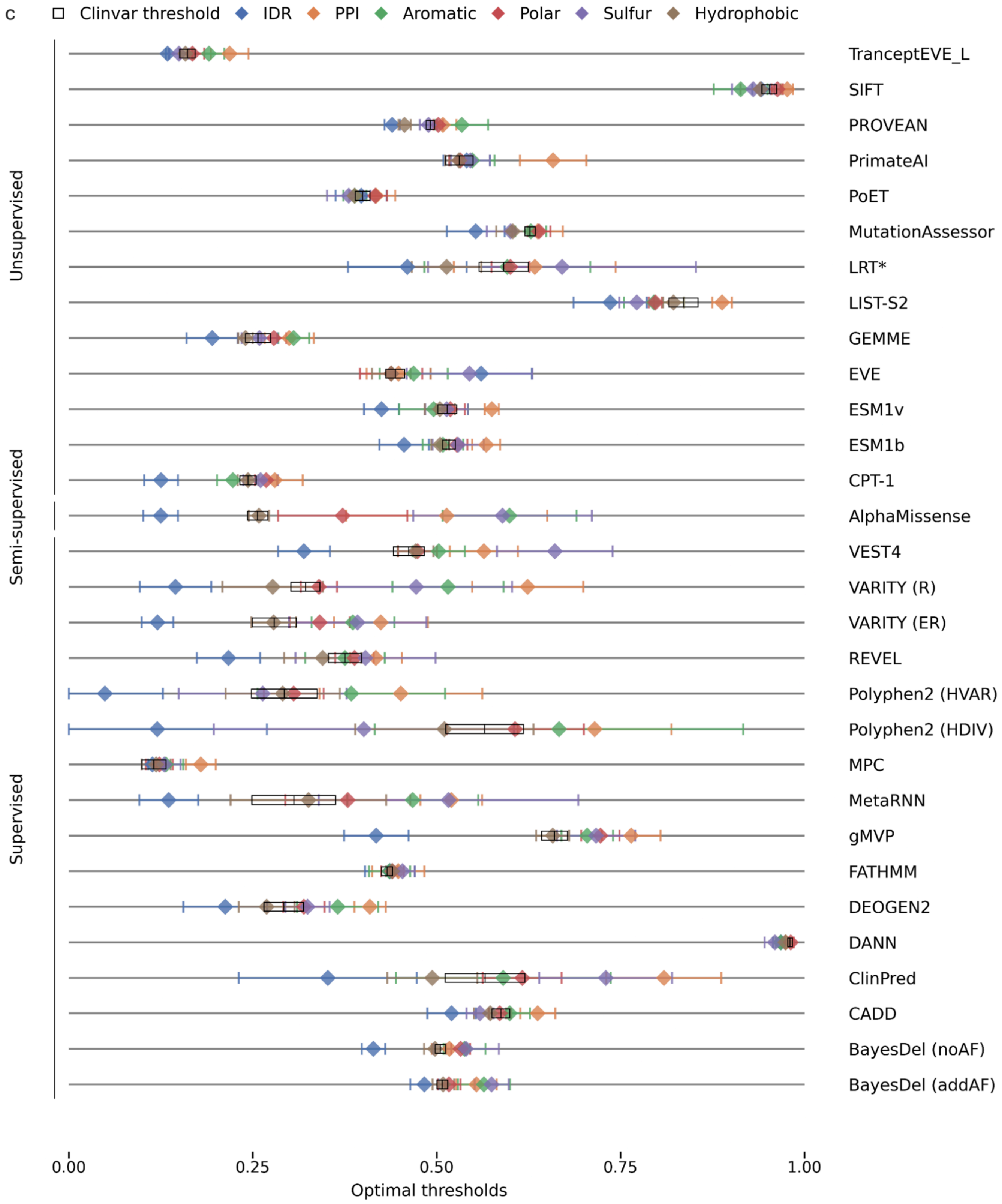

**Figure S1: Optimal classification threshold shifts across VEPs (extended).** Shifts in optimal discrimination thresholds (diamonds; maximal J-statistic) across normalized model scores in the ClinVar_BM dataset. Error bars indicate ±1 SD over 1,000 non-parametric bootstrap iterations. Black rectangle marks the naive threshold (mean ± SD) across the entire ClinVar_BM dataset. * log-transform applied to raw model scores (Methods) (a) full ClinVar_BM dataset (b-c) M1 variants excluded.

**Figure S2.** Changes in AUROC across residue subgroups

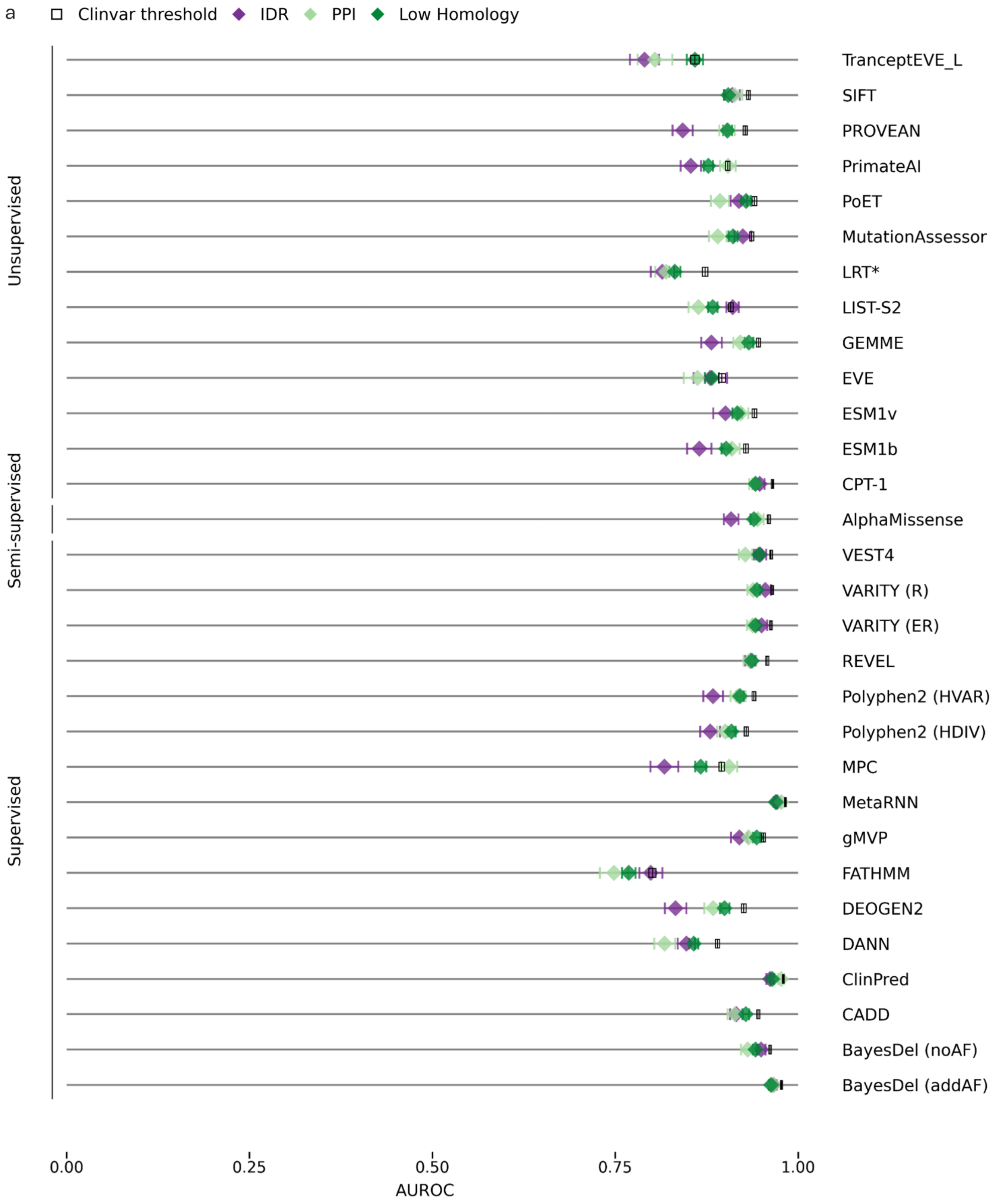

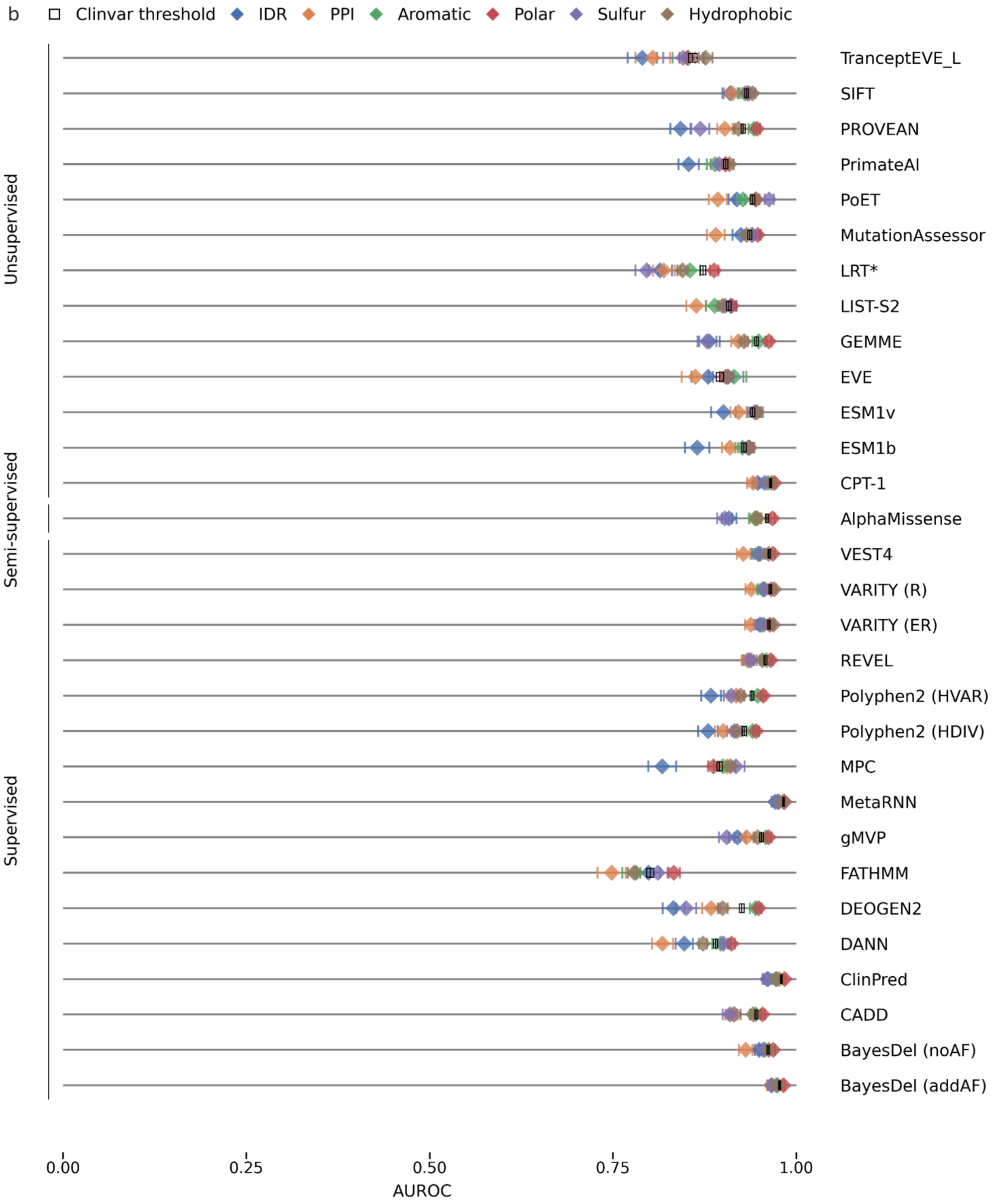

b
Clinvar threshold
IDR
PPI
Aromatic
Polar
Sulfur
Hydrophobic
Unsupervised
Semi-supervised
Supervised
TranceptEVE_L
SIFT
PROVEAN
PrimateAI
PoET
MutationAssessor
LRT*
LIST-S2
GEMME
EVE
ESM1v
ESM1b
CPT-1
AlphaMissense
VEST4
VARITY (R)
VARITY (ER)
REVEL
Polyphen2 (HVAR)
Polyphen2 (HDIV)
MPC
MetaRNN
gMVP
FATHMM
DEOGEN2
DANN
ClinPred
CADD
BayesDel (noAF)
BayesDel (addAF)
0.00
0.25
0.50
0.75
1.00
AUROC

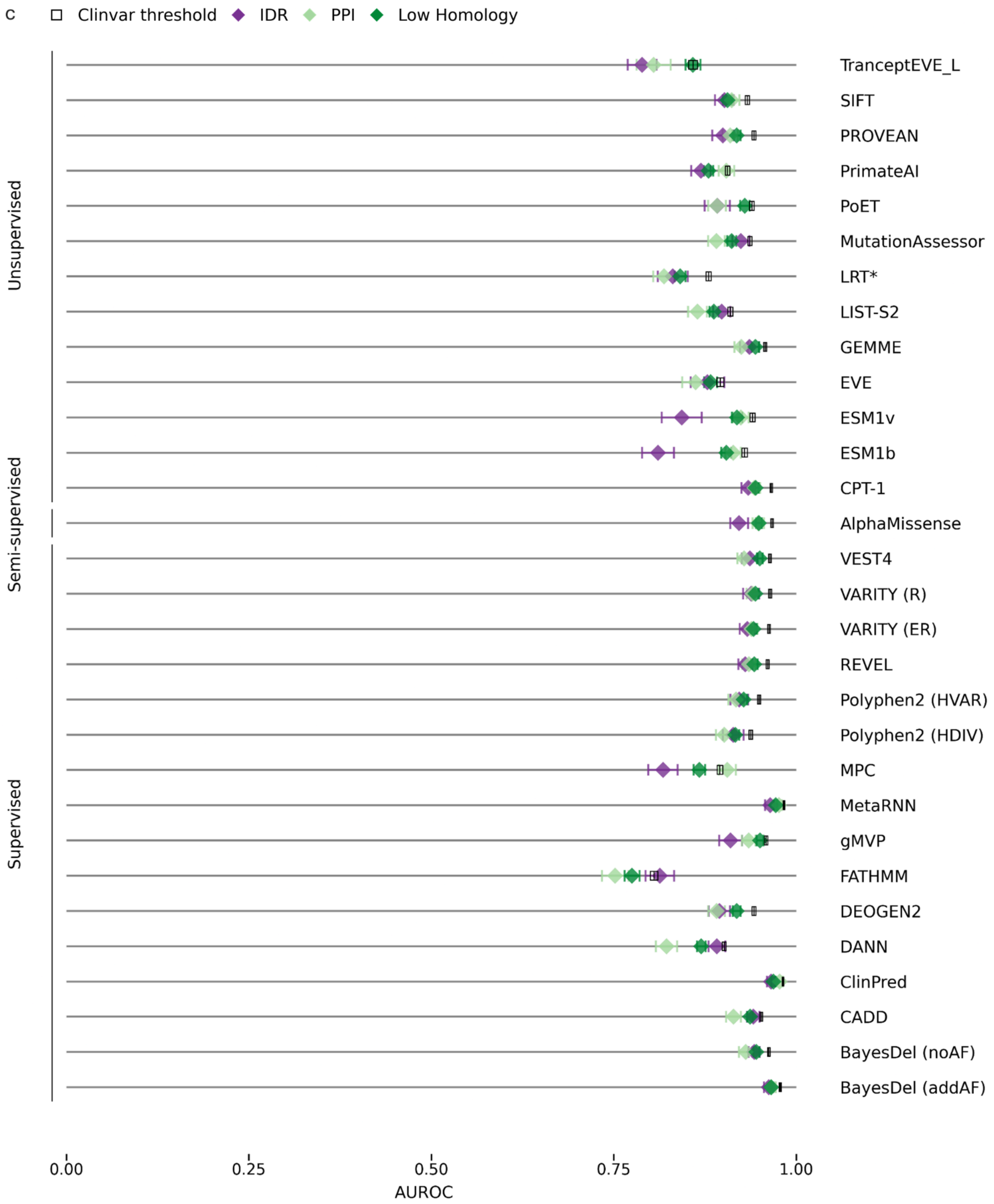
c
Clinvar threshold
IDR
PPI
Low Homology
Unsupervised
Semi-supervised
Supervised
TranceptEVE_L
SIFT
PROVEAN
PrimateAI
PoET
MutationAssessor
LRT*
LIST-S2
GEMME
EVE
ESM1v
ESM1b
CPT-1
AlphaMissense
VEST4
VARITY (R)
VARITY (ER)
REVEL
Polyphen2 (HVAR)
Polyphen2 (HDIV)
MPC
MetaRNN
gMVP
FATHMM
DEOGEN2
DANN
ClinPred
CADD
BayesDel (noAF)
BayesDel (addAF)
0.00
0.25
0.50
0.75
1.00
AUROC

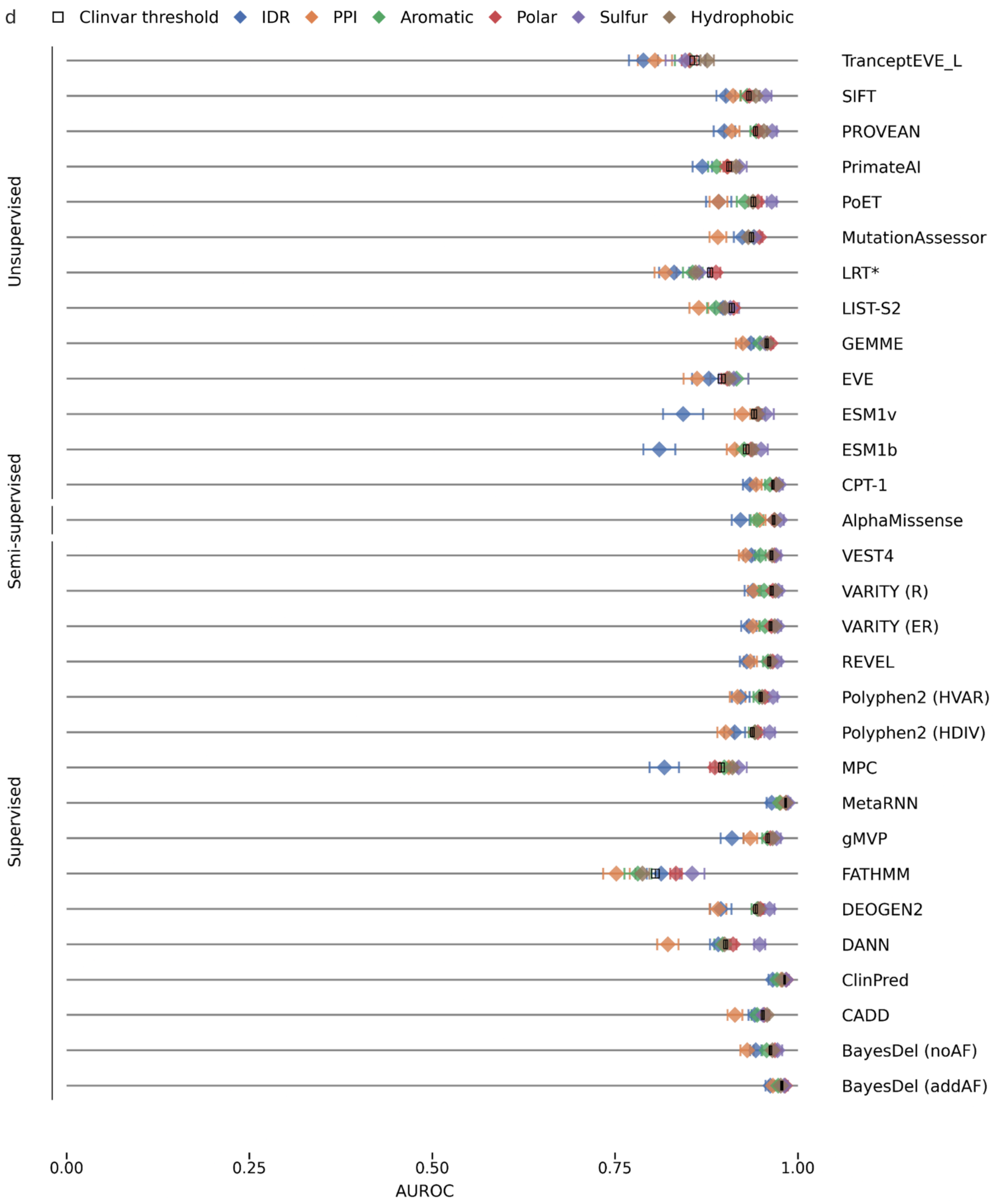

**Figure S2. VEP performance across variant subgroups.** Performance shifts shown as changes in AUROC (Diamonds) across all variants within each subgroup: (a,c) attributes showing the most significant shifts and (b,d) extended attributes set. Error bars indicate ± 1 SD over 1,000 non-parametric bootstrap iterations. Black rectangle marks the mean ± 1SD across the entire ClinVar_BM dataset. * log-transform applied to raw model scores (Methods) (a-b) full ClinVar_BM dataset (c-d) M1 variants excluded.

**Figure S3.** ECE rank heatmaps across residue attributes and predictors.

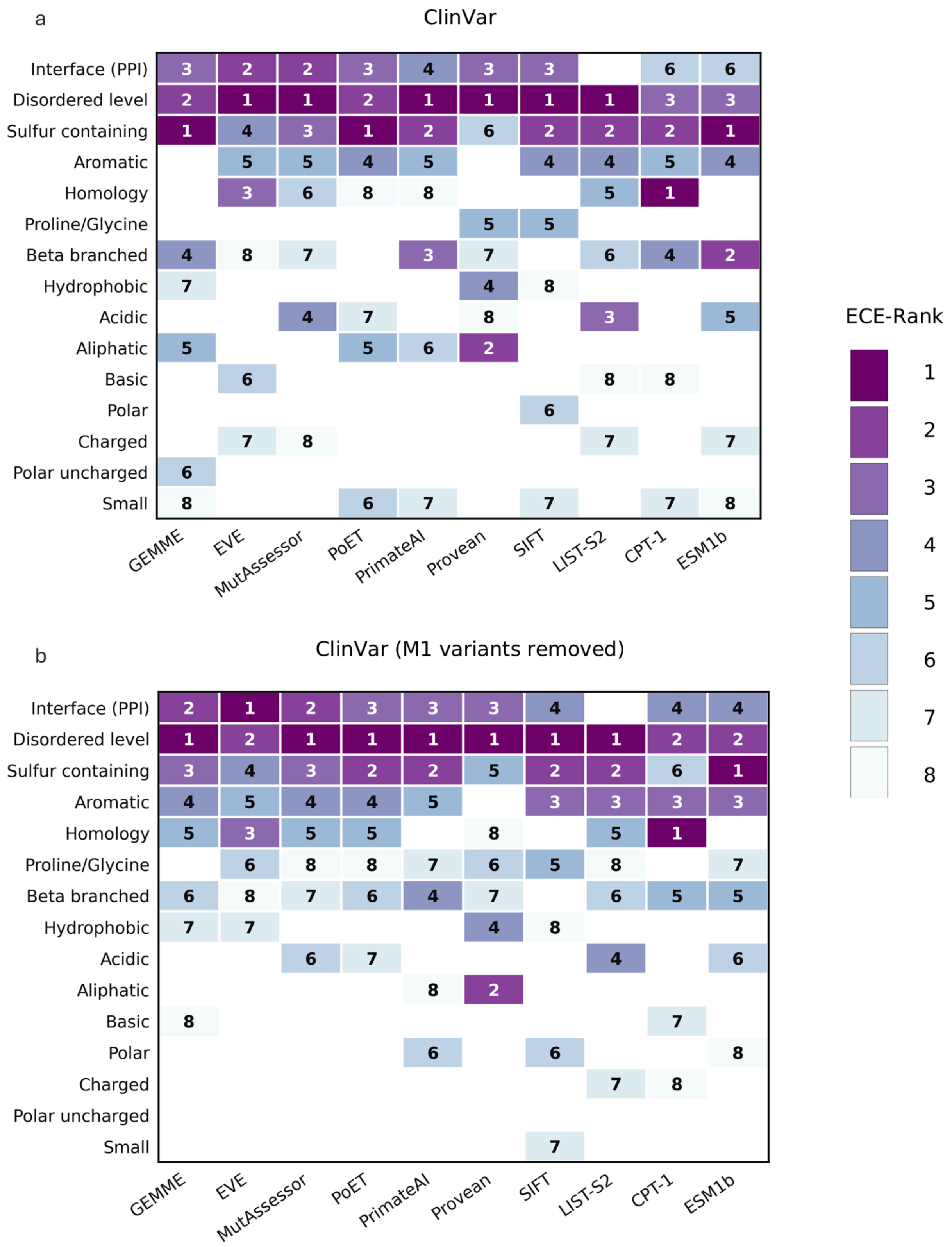

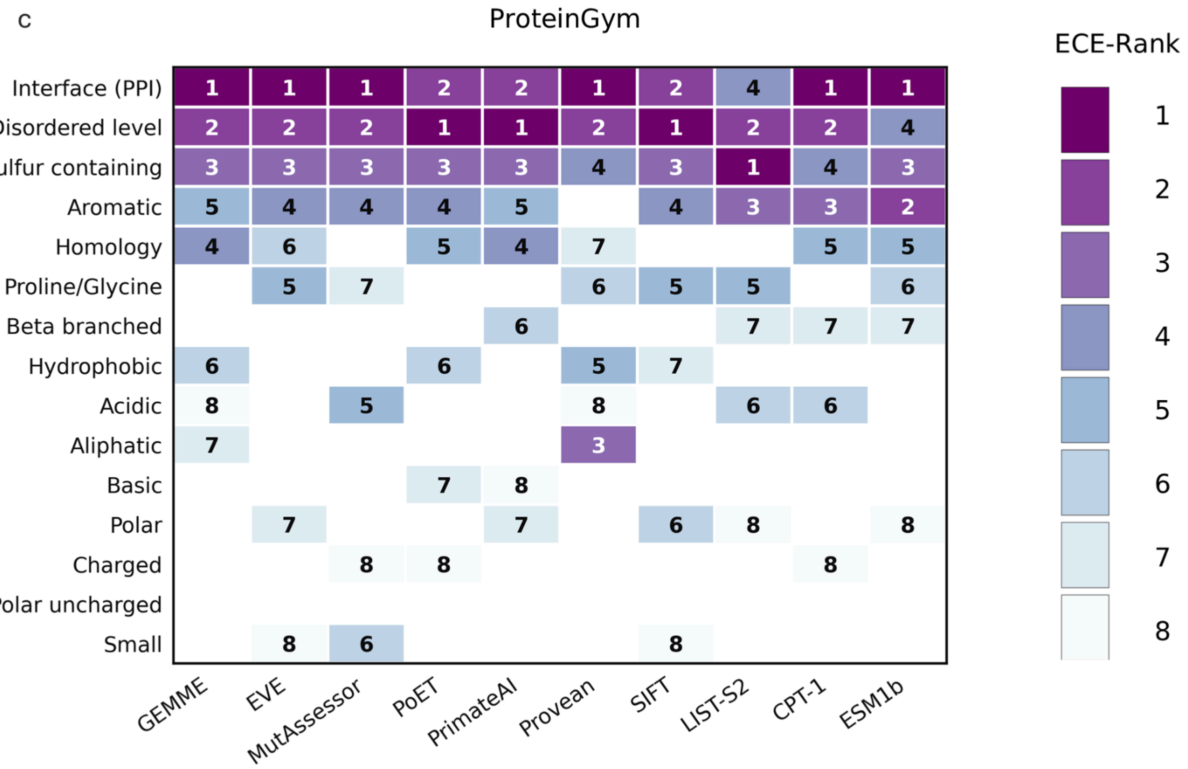

**Figure S3. Extended ECE rank heatmaps.** ECE ranks after global calibration using 6,000 training samples per iteration across residue attributes and unsupervised predictors in ClinVar_Balanced (a-b) and ProteinGym clinical substitution benchmark (c). Panel b repeats the ClinVar_Balanced analysis excluding first-methionine variants (n=634). Cells show within-predictor ranks based on mean ECE across 100 calibration iterations; rank 1 indicates the greatest miscalibration. Darker shading denotes a higher miscalibration rank. Blank cells indicate attributes outside the top eight ECE ranks. Exact ECE values are provided in Table S2.

**Figure S4.** LLR score distributions across residue subgroups in ClinVar and ProteinGym

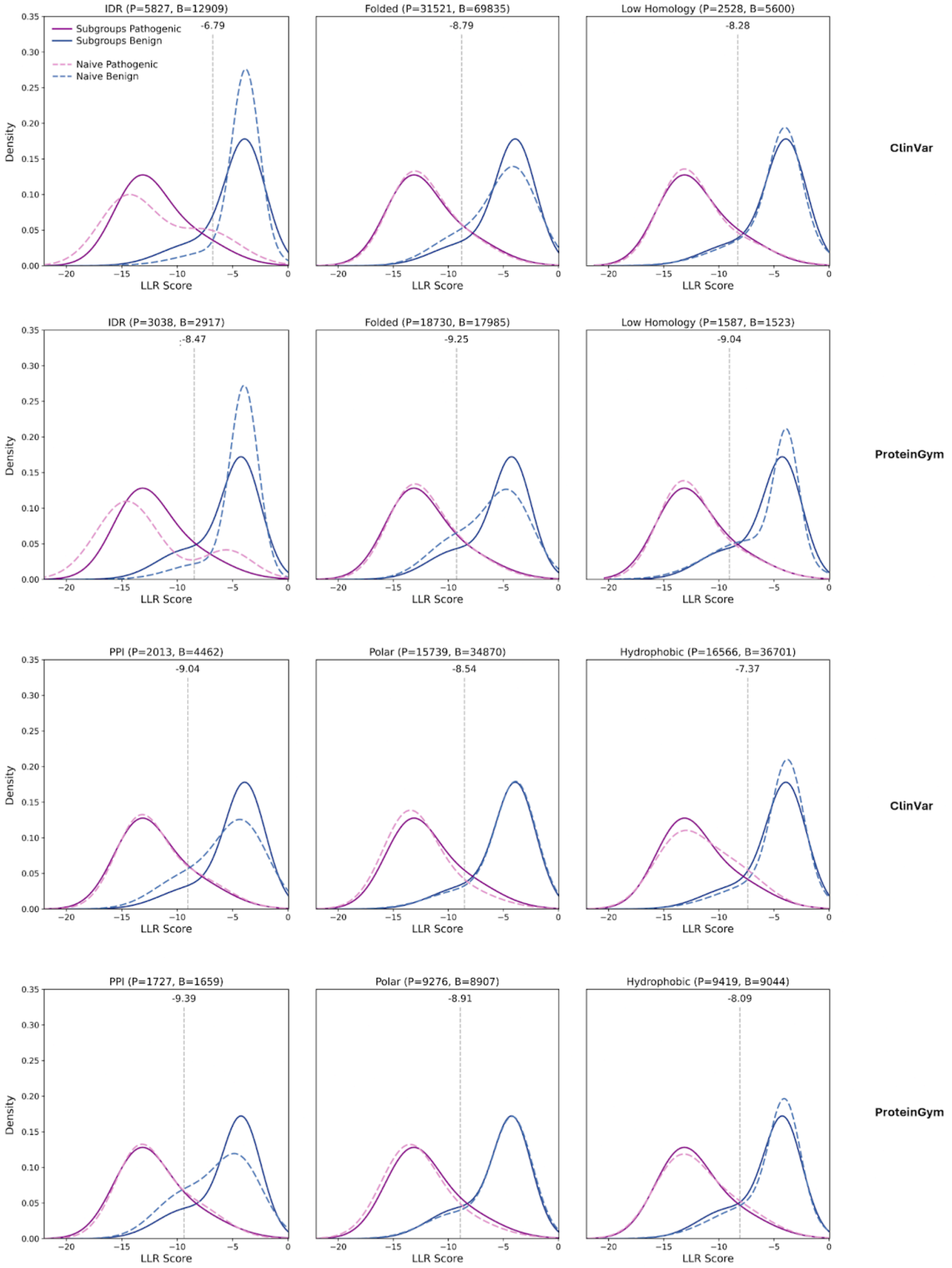

**Figure S4: LLR distribution of selected attributes.** Distribution of LLR scores for pathogenic and benign variants in ClinVar_HQ and ProteinGym datasets. Dotted purple and blue curves - pathogenic and benign GMMs distribution over the entire dataset. Solid curves - subgroup specific GMMs distributions. *P*, *B* denote the number of pathogenic and benign variants per subgroup. Dotted line - optimal classification thresholds (J-statistic) Distributions fitted using two-component GMMs.

**Figure S5.** LLR-score distribution differences with different scoring strategies

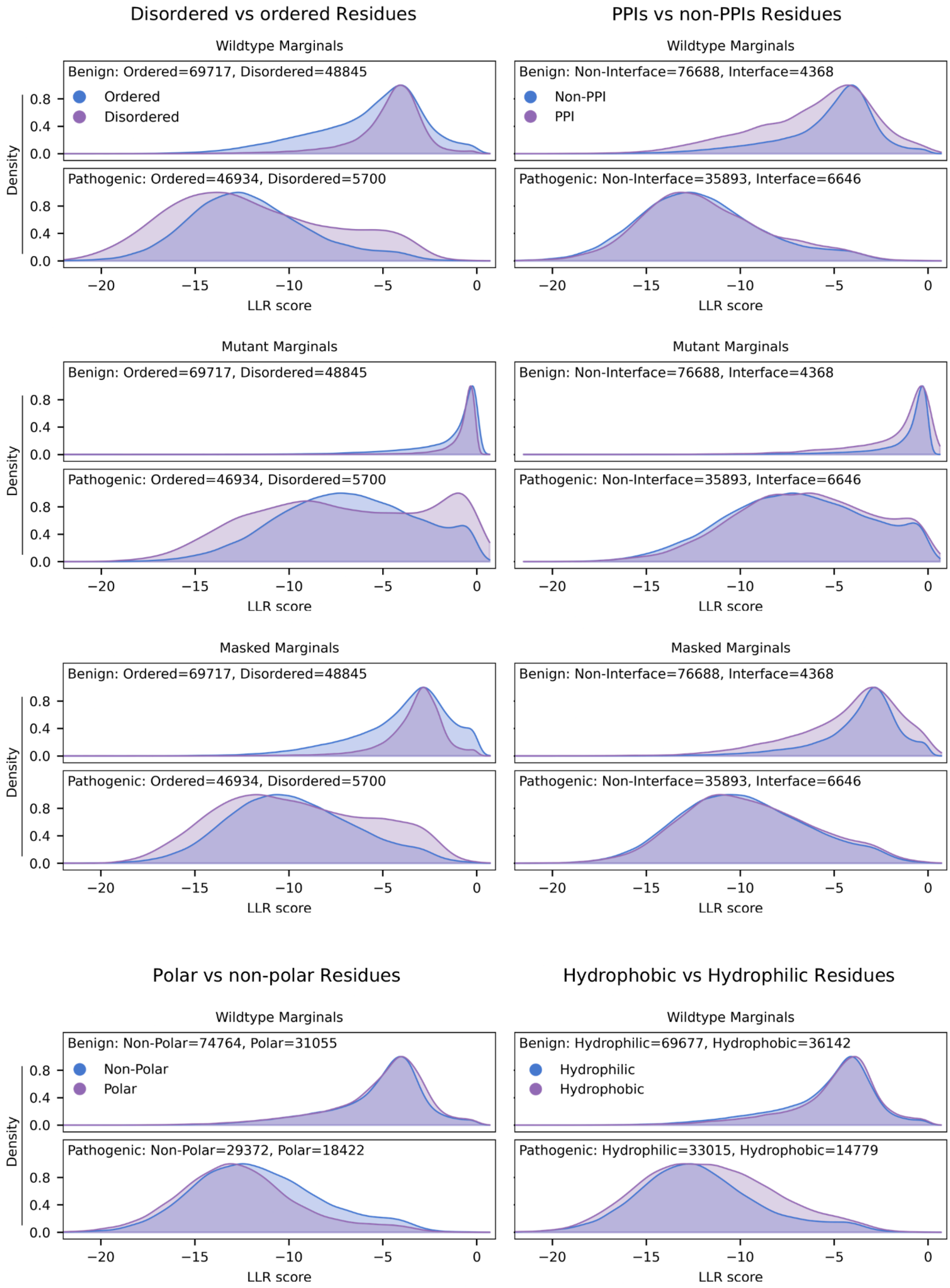

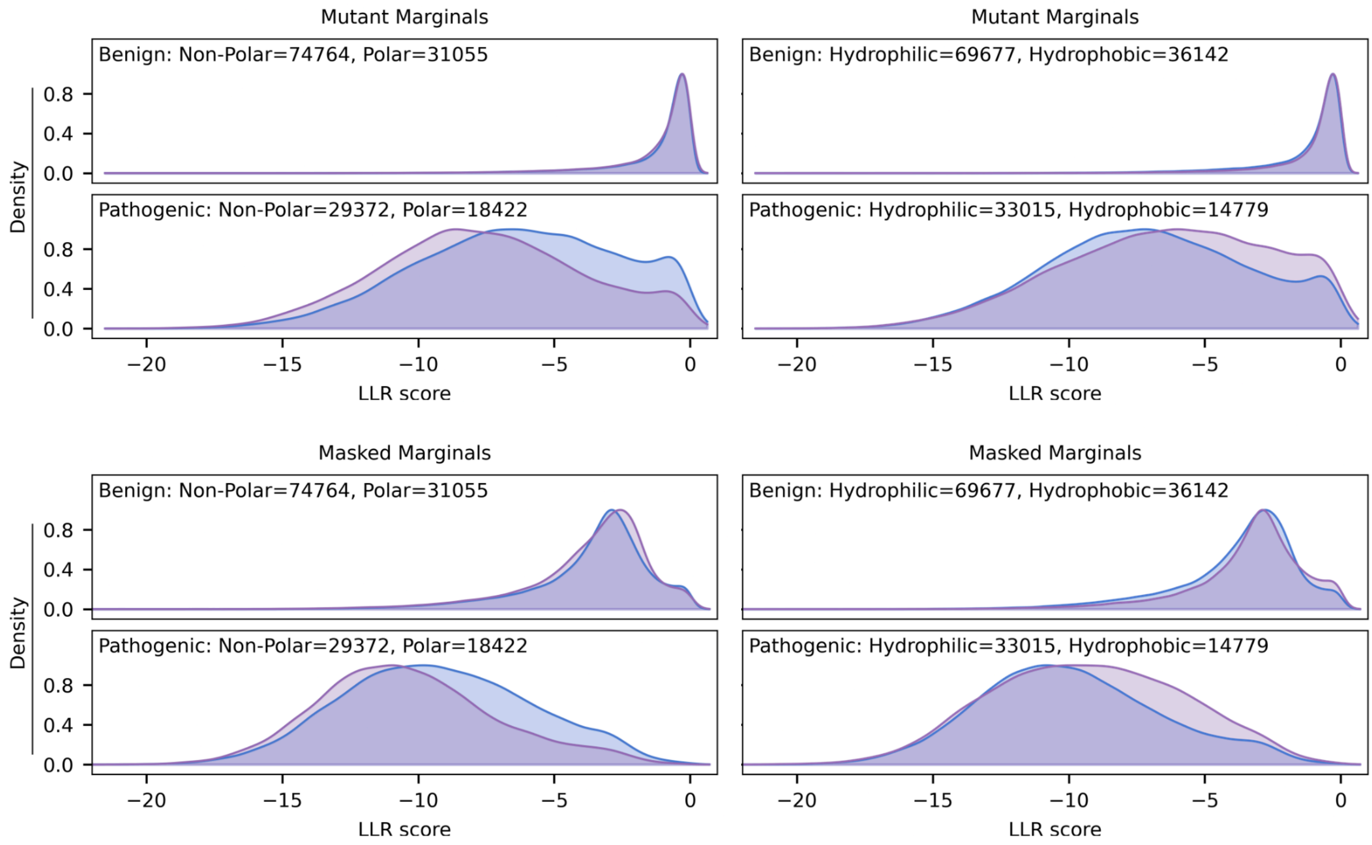

**Figure S5. LLR-score distributions under different input representations.** LLR-score distributions for selected residue-level attributes under wild-type (top), mutant (center), and masked (bottom) sequence representations, grouped by pathogenic (left) and benign (right) variants.

**Figure S6.** Effect of M1 exclusion on ESM1b calibration

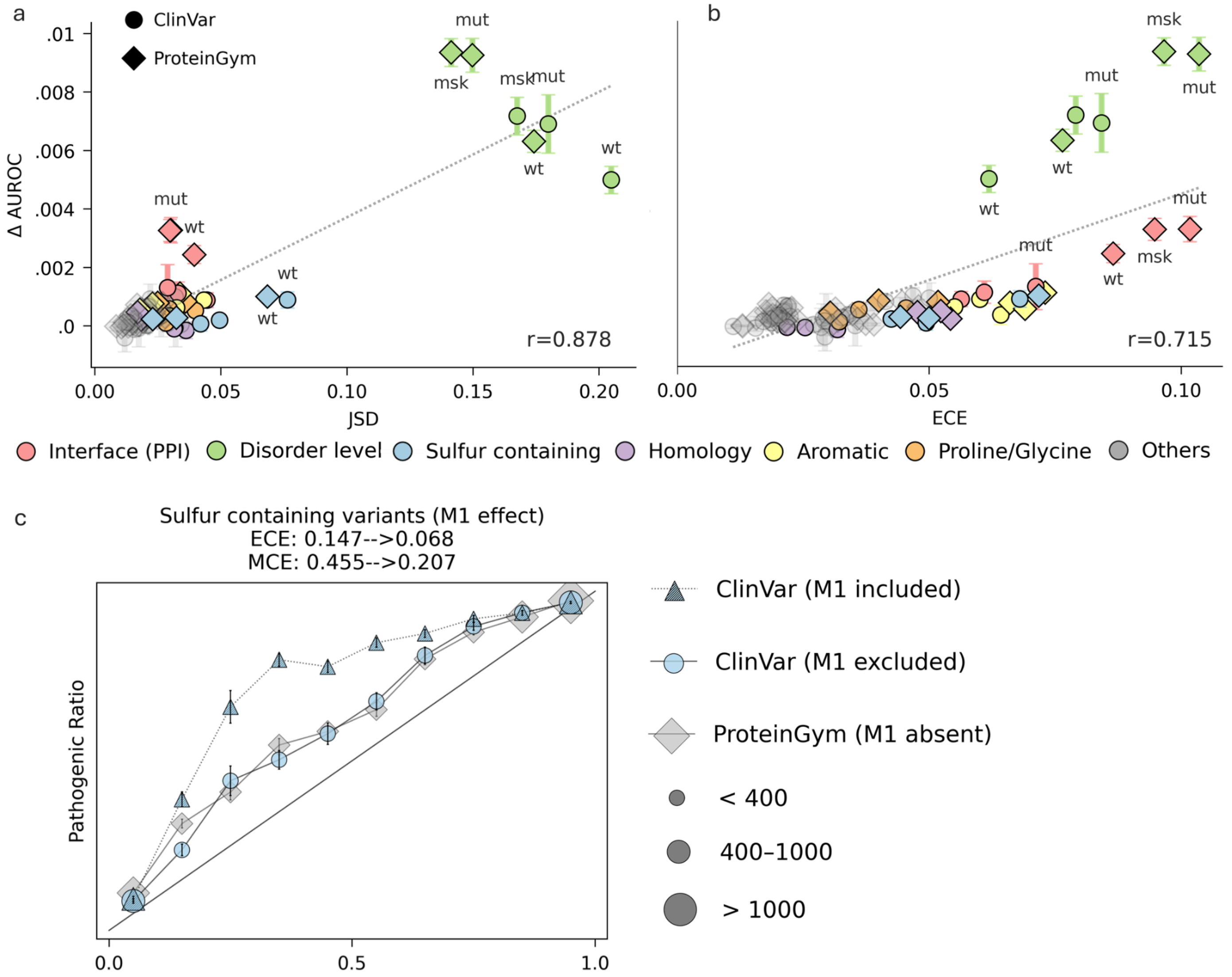

**Figure S6. Effect of M1 exclusion on ESM1b calibration.** (**a-b**) Change in global AUROC (y-axis; relative to the uncalibrated model) as a function of changes in score distribution (JSD) (**a**) and ECE (**b**), with first-methionine variants excluded from ClinVar_Balanced (n=634). Differential calibration was performed by fitting separate subgroup-specific logistic rescaling functions using one-third of the data for training and two-thirds for testing (Methods). Error bars indicate ±1 SD across 100 non-parametric bootstrap test iterations per fold. Circles and diamonds correspond to results on the ClinVar_Balanced and ProteinGym clinical substitution benchmarks, respectively. Colored symbols highlight selected calibrated attributes; transparent gray symbols denote other tested attributes (Table S3). wt, mut, and msk denote wild-type, mutant, and masked sequence representations, respectively. Dotted lines indicate linear regression fits. (**c**) Reliability histogram for sulfur-containing variants shows predicted confidence (x-axis, 10 equal-width bins) versus observed pathogenic frequency (y-axis). Perfect calibration follows y=x; lines show the mean trend across datasets: ClinVar_Balanced with M1 included (triangles), ClinVar_Balanced with M1 excluded (circles) and ProteinGym (diamonds); error bars indicate ±1 SD across 100 global calibration iterations each using 6,000 randomly sampled variants for training and the remaining variants for testing. Marker size denotes bin sample count. ECE = Expected Calibration Error, MCE = Maximal calibration error (Methods).

**Figure S7.** Effects of score-distribution and pathogenic-fraction differences on differential calibration

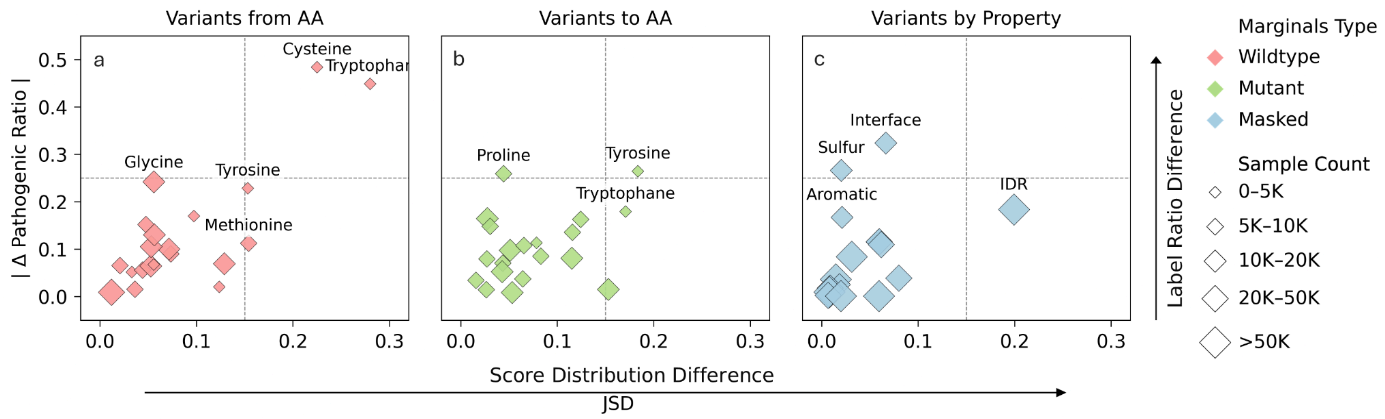

**Figure S7. Guiding calibration with subgroup-specific score distributions.** Changes in pathogenic fraction across residue-subgroups often occur without significant changes in score-distribution but the reverse is rare. y-axis; absolute deviation from the global pathogenic fraction. x-axis; Jensen-Shannon divergence of scores distribution between subgroup partitions. Variants are grouped by substitutions of the wild-type amino acid (a), substitutions to specific amino acids (b), and residue property-based variants (c). Diamond size denotes the smaller subgroup in each partition, and color indicates the scoring strategy (pink - WT, green - MUT, blue - MSK).

**Figure S8.** matrices to estimate AUROC gains from differential calibration

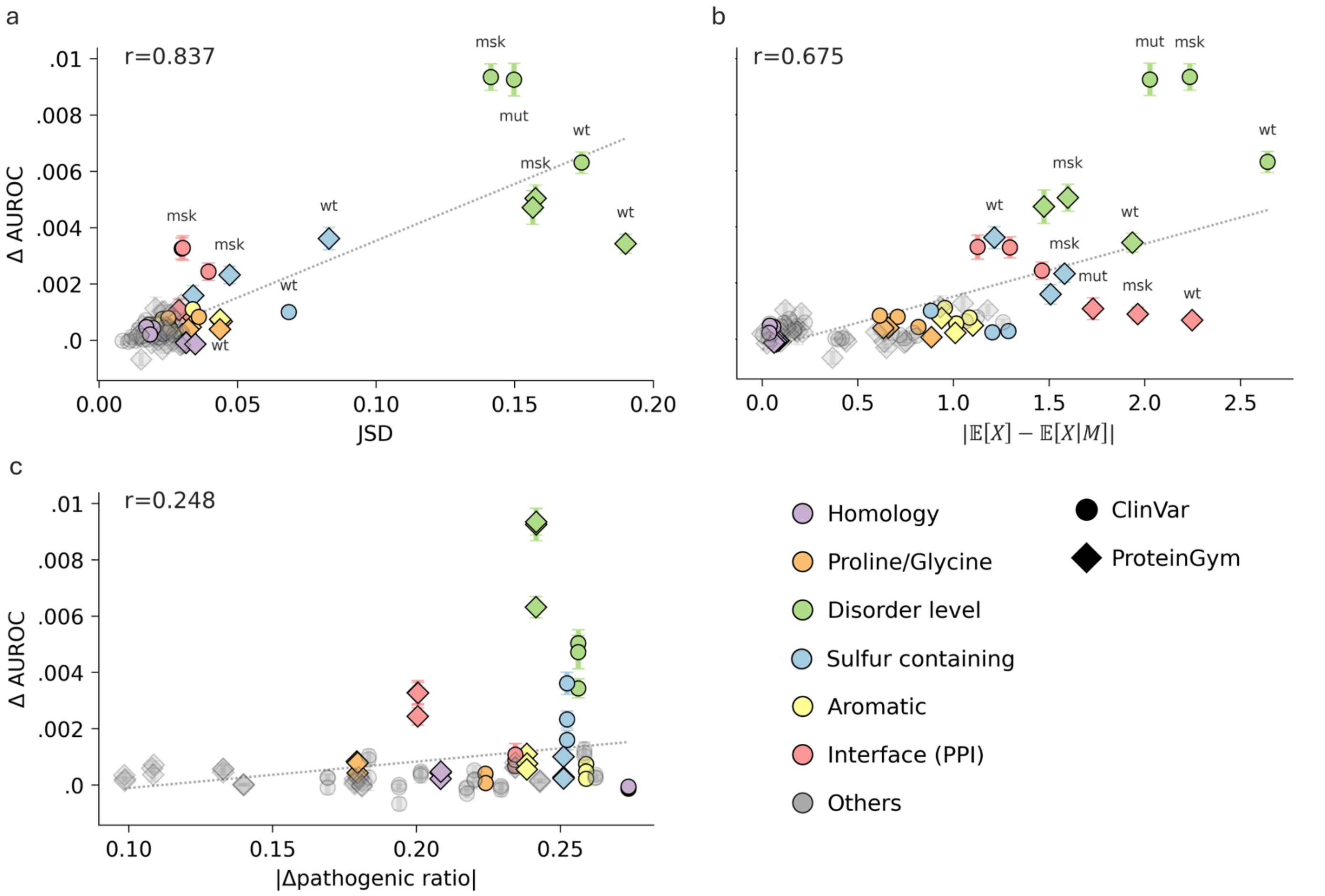

**Figure S8. Alternative metrics for predicting AUROC gains from differential calibration (ESM1b).** Global AUROC change (ΔAUROC; y-axis, relative to the uncalibrated model) as a function of (a) class-conditional score-distribution divergence (JSD), (b) the absolute difference in mean LLR scores between the full dataset and subgroup, and (c) the absolute difference in pathogenic fraction between the full dataset and subgroup. JSD shows the strongest association with AUROC gains followed by mean-score differences whereas pathogenic-fraction differences show a weak association. Differential calibration was performed by fitting separate subgroup-specific logistic rescaling functions using one-third of the data for training and two-thirds for testing (Methods). Error bars indicate ±1 SD across 100 non-parametric bootstrap test iterations per fold. Circles and diamonds denote ClinVar_Balanced and ProteinGym clinical substitution benchmark, respectively; colored symbols indicate the residue-level attribute, and transparent gray symbols denote other tested attributes (Table S3). **wt**, **mut**, and **msk** denote wild-type, mutant, and masked sequence representations, respectively. Dotted lines indicate linear regression fits.

**Figure S9.** Normalized JSD correlation with AUROC gains

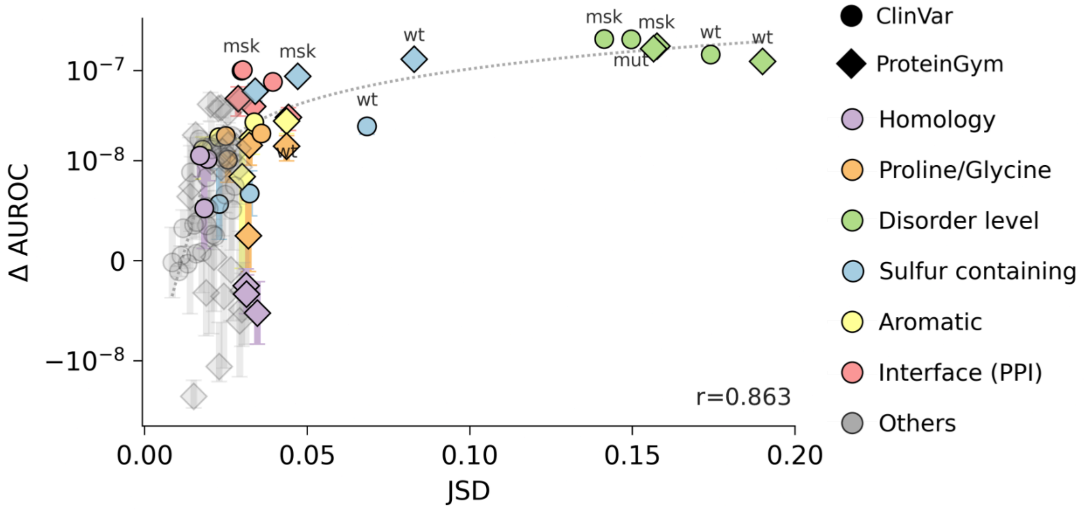

**Figure S9. Relative contribution of score-distribution differences to AUROC gain (ESM1b).** Normalized change in overall AUROC (y-axis; relative to the uncalibrated model, log scale) as a function of class-conditional score-distribution shift (x-axis; Jensen–Shannon divergence). Normalization was calculated by dividing ΔAUROC by the number of samples per subgroup (minimum count between positive and negative selections). The normalized AUROC contribution increases with score-distribution divergence; Pearson correlation coefficient (*r*) is shown. Differential calibration was performed using one-third of the data for training and two-thirds for testing (Methods). Error bars indicate ±1 SD across three training folds and 100 non-parametric bootstrap test iterations per fold. Circles and diamonds denote ClinVar_Balanced and ProteinGym, respectively; symbol color denotes the calibrated residue attribute (legend), and transparent gray symbols denote other tested attributes (Table S3). Dashed lines indicate linear regression fits.

**Figure S10.** Protein-level aggregation of pathogenic-variant distributions across residue subgroups

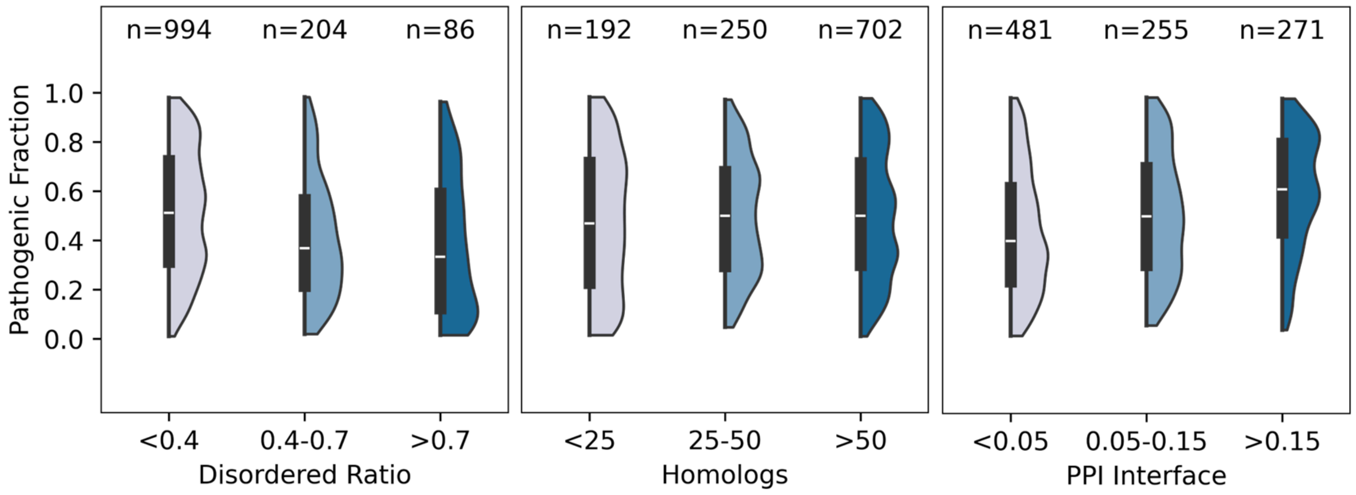

**Figure S10. Distribution of pathogenic variants at the protein level.** Trends captured at the residue level are also observed at the protein level. Distribution of pathogenic variants (y-axis) as a function of residue-level attributes (x-axis) in 1,285 well-annotated ClinVar proteins.

**Figure S11.** RaCoon's Calibration Tree

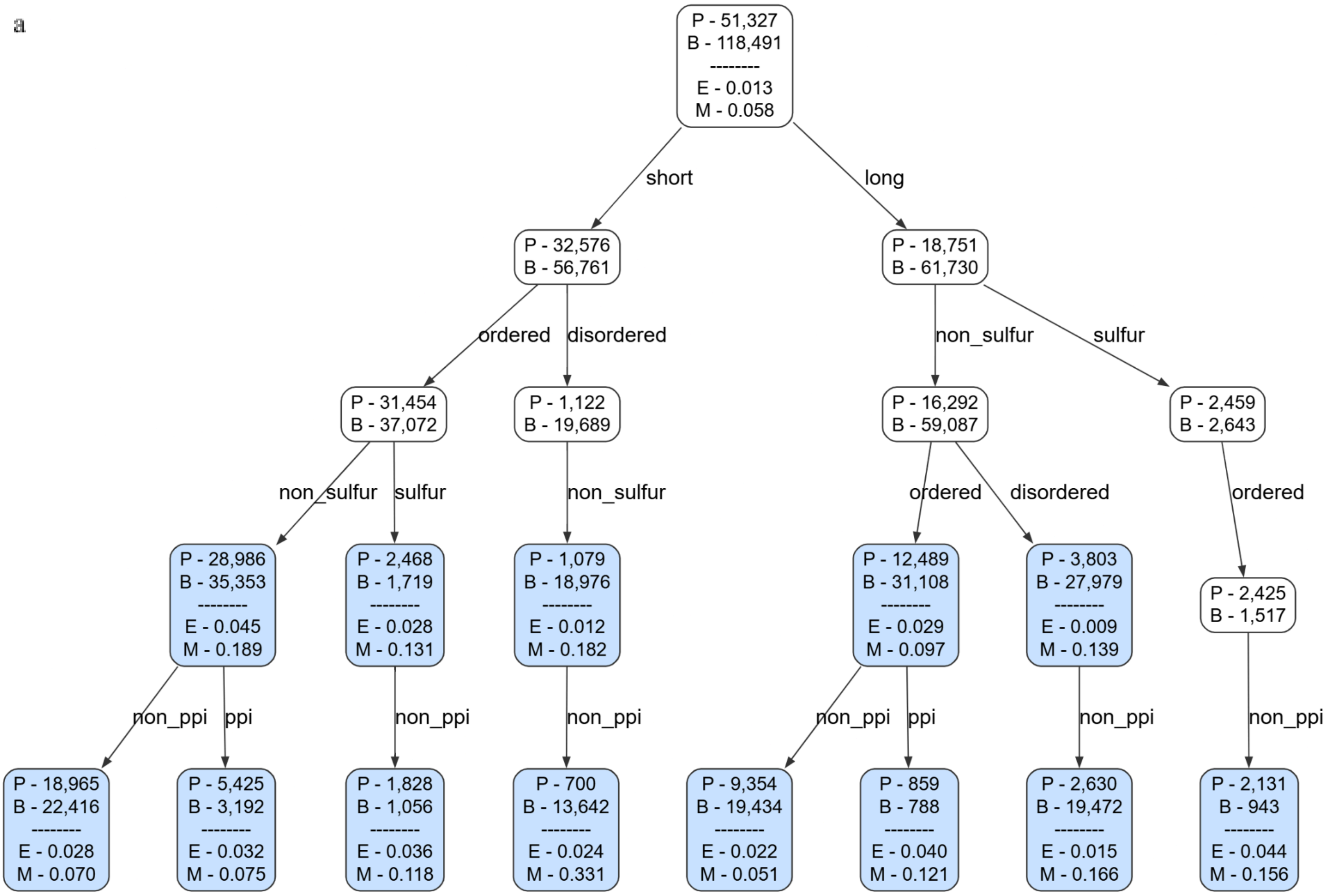

b

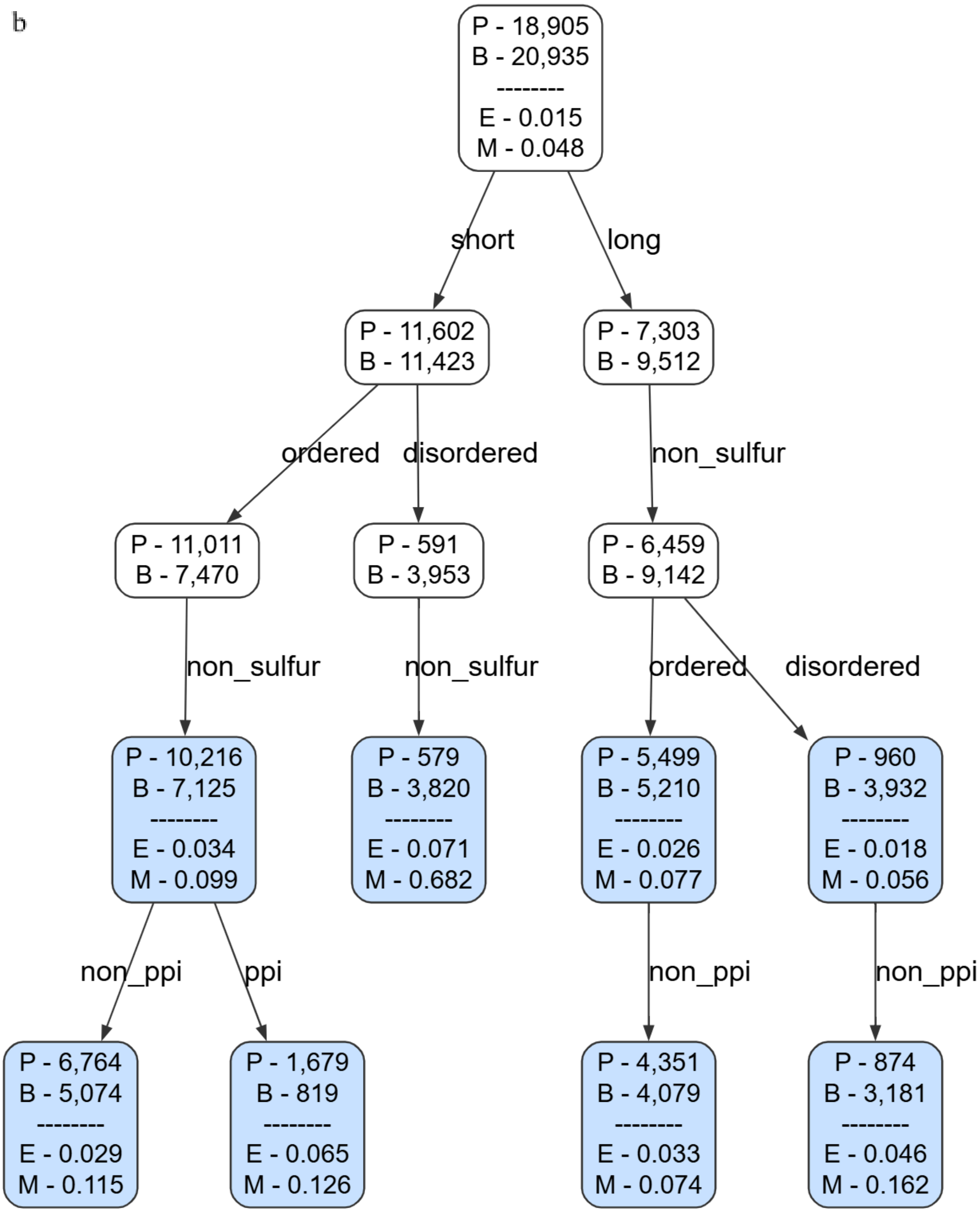
P - 18,905
B - 20,935
--------
E - 0.015
M - 0.048
short
long
P - 11,602
B - 11,423
P - 7,303
B - 9,512
ordered
disordered
non_sulfur
P - 11,011
B - 7,470
P - 591
B - 3,953
P - 6,459
B - 9,142
non_sulfur
non_sulfur
ordered
disordered
P - 10,216
B - 7,125
--------
E - 0.034
M - 0.099
P - 579
B - 3,820
--------
E - 0.071
M - 0.682
P - 5,499
B - 5,210
--------
E - 0.026
M - 0.077
P - 960
B - 3,932
--------
E - 0.018
M - 0.056
non_ppi
ppi
non_ppi
non_ppi
P - 6,764
B - 5,074
--------
E - 0.029
M - 0.115
P - 1,679
B - 819
--------
E - 0.065
M - 0.126
P - 4,351
B - 4,079
--------
E - 0.033
M - 0.074
P - 874
B - 3,181
--------
E - 0.046
M - 0.162

c

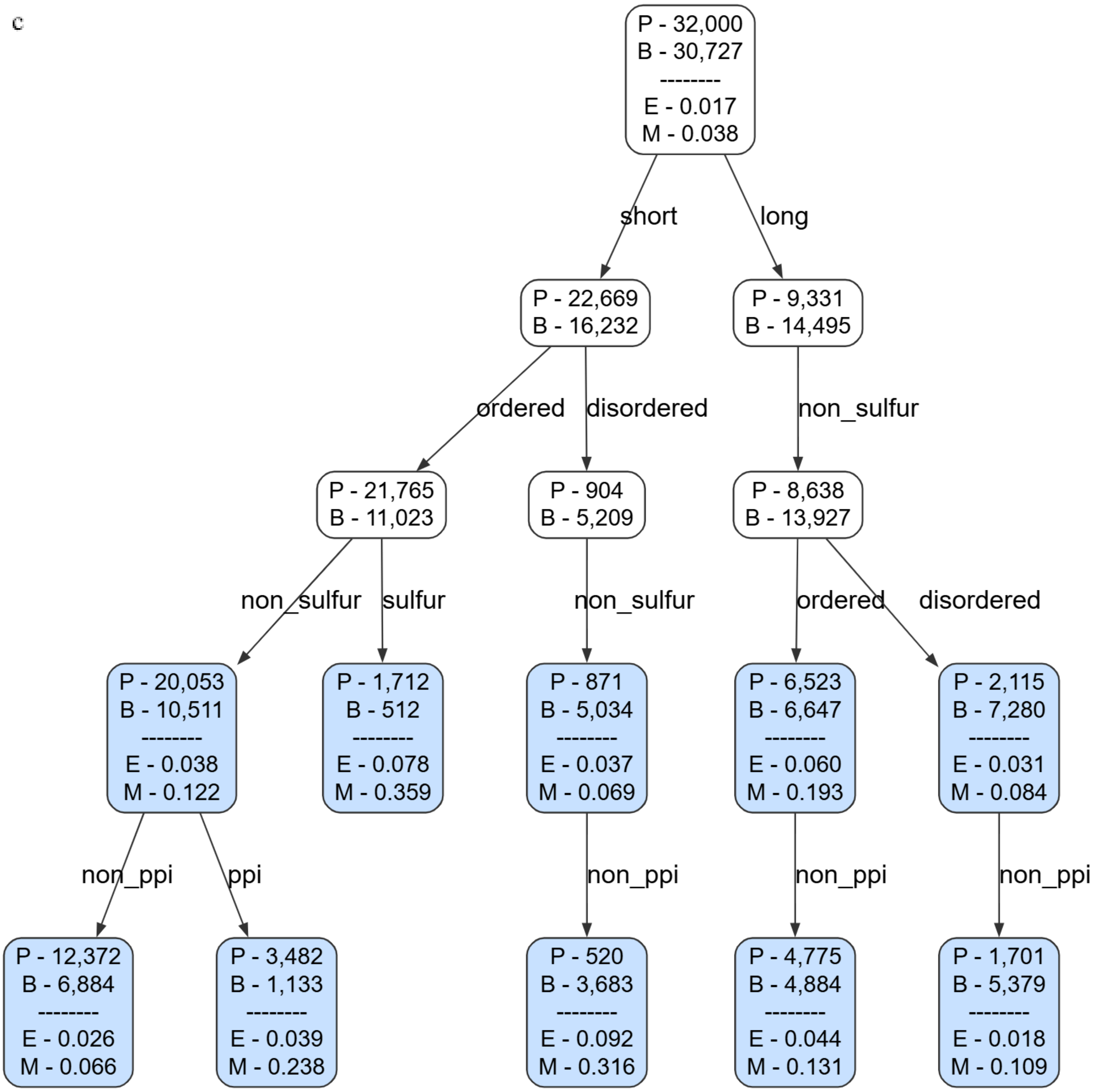

**Figure S11. Different calibration trees obtained by RaCoon across datasets (M1 variants excluded).** Calibration trees obtained after partitioning and pruning the calibration dataset (Methods, Algorithms 1–2). Leaf retention thresholds: ≥400 pathogenic, ≥400 benign, and ≥1,600 total variants. Blue nodes denote calibrated subgroups included in the ECE/MCE assessment (leaves and eligible internal nodes). P and B denote the numbers of pathogenic and benign variants per node, respectively (training and test combined). E and M denote ECE and MCE, respectively, computed on held-out variants mapped to each node. These metrics are omitted for sparse internal nodes with fewer than 400 mapped test variants. MCE is calculated only for bins containing at least 50 variants, which can result in MCE values lower than ECE. **(a)** Calibration: ClinVar_HQ; test: ClinVar_HQ. **(b)** Calibration: ClinVar_Balanced; test: ProteinGym. **(c)** Calibration: ProteinGym; test: ProteinGym.

**Figure S12.** Racoon hyperparameters abalations

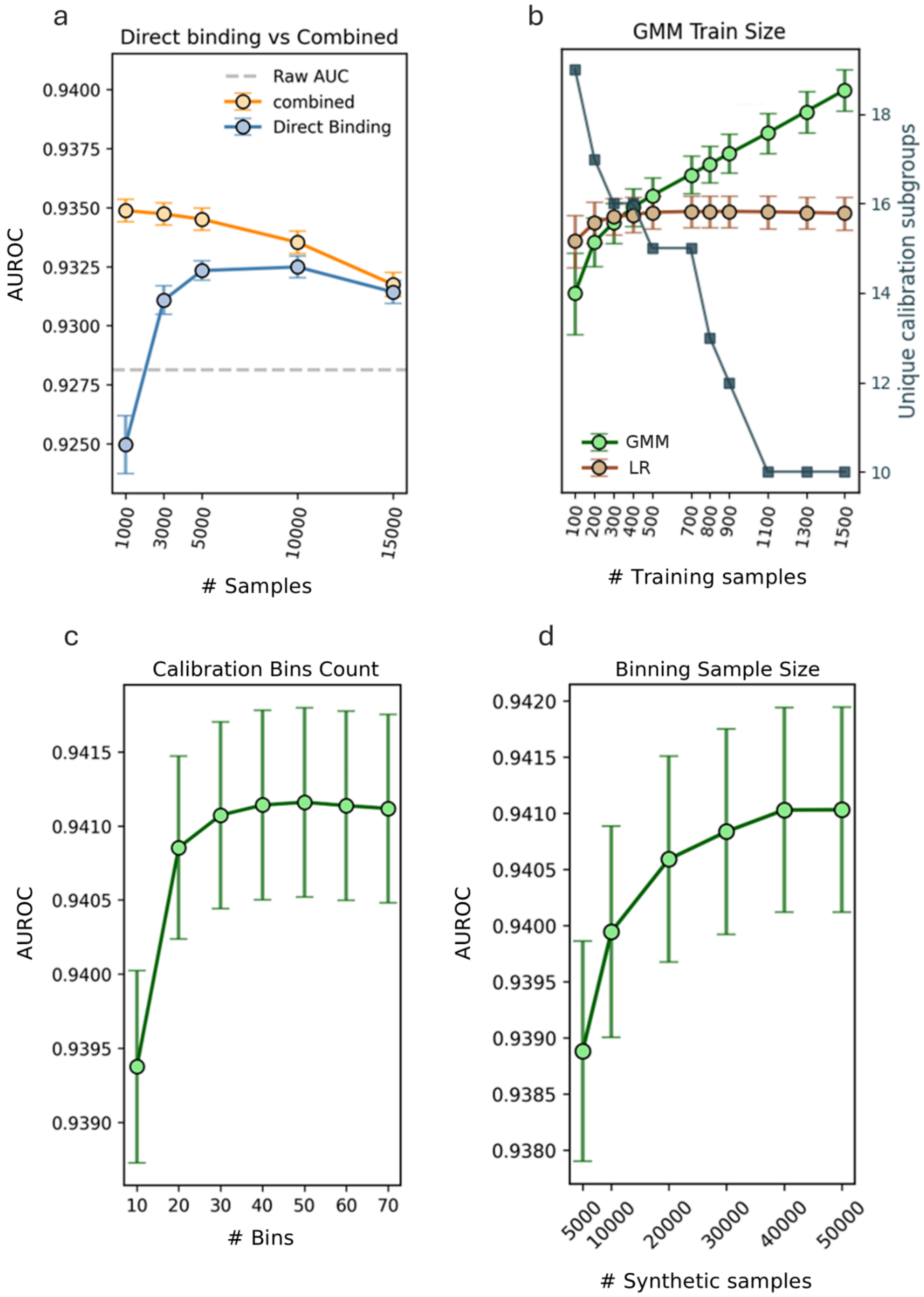

**Figure S12. Ablation studies of the RaCoon pipeline.** All results show changes in global AUROC on ClinVar_HQ. Test samples were not used during calibration. Error bars indicate ± 1 SD across 100 training (calibration) iterations, each averaged over 100 non-parametric bootstrap test iterations. **a.** Effect of binning method on AUROC: direct binning of raw LLR scores (blue) versus a hybrid method combining $n$ = 40,000 synthetic samples from benign and pathogenic GMMs fitted to raw LLRs (orange).**b.** Effect of training size on AUROC for GMM-based (green) and logistic-regression (LR; brown) calibration and the number of retained subgroups (blue, right axis). The x-axis denotes $n$ variants per GMM; logistic regression uses $2n$ randomly sampled variants per node. Higher training requirements raise pruning thresholds, leaving fewer subgroups. **c.** Effect of the calibration histogram bins on AUROC using 100,000 synthetic samples from the benign and pathogenic GMMs. **d.** Effect of the number of synthetic GMMs samples on AUROC using 50 calibration bins.

**Figure S13** RaCoon performance evaluation after M1 exclusion

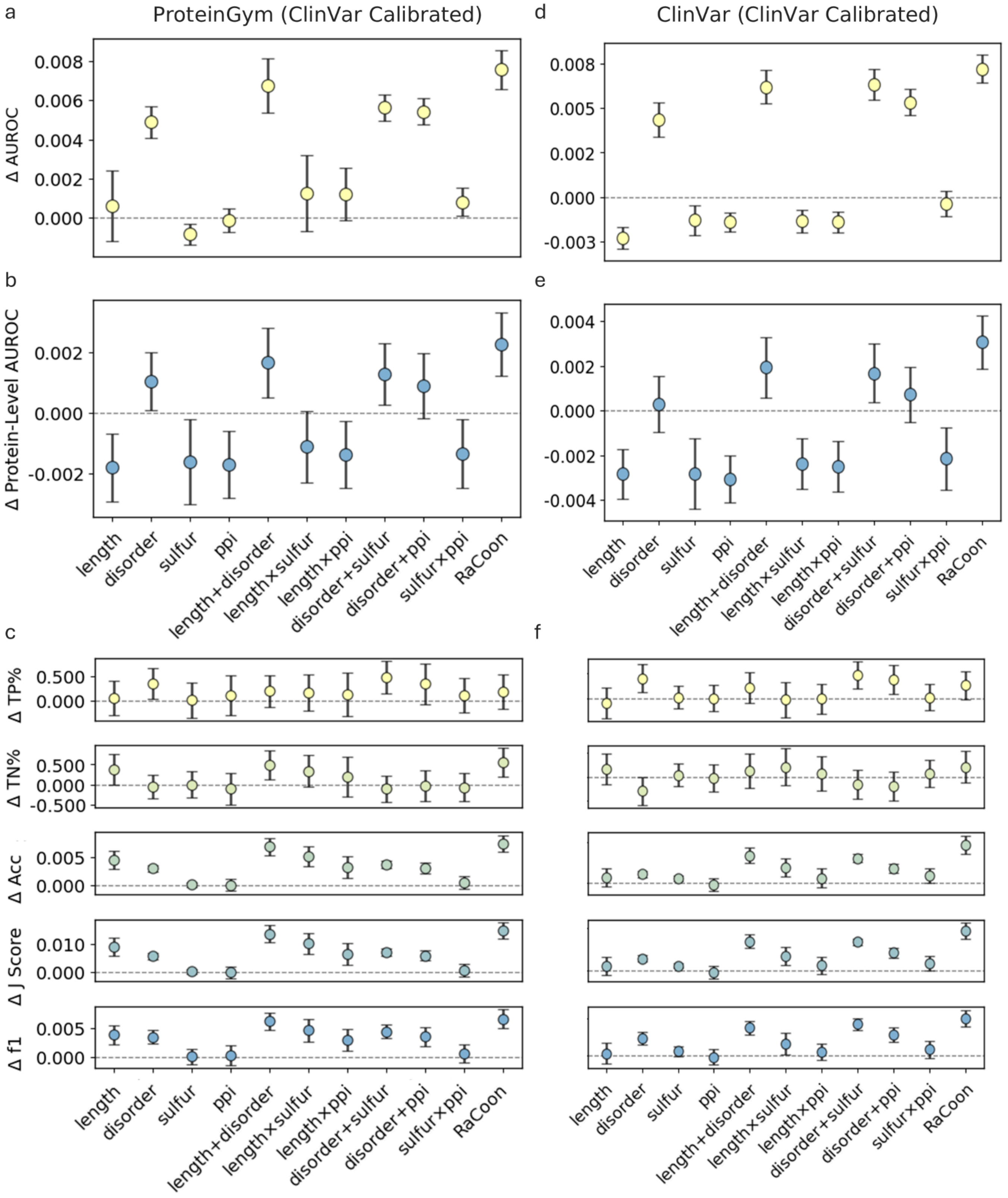

**Figure S13. RaCoon performance evaluation after M1 exclusion.** Changes in global AUROC (**a,d**), mean per-protein AUROC (**b,e**), and classification metrics (c,f), relative to uncalibrated ESM1b. Calibration was performed on ClinVar_Balanced, with evaluation on ProteinGym (a–c) or held-out ClinVar_Balanced variants (**d-f**). M1 variants (n = 634) were excluded from ClinVar_Balanced before partitioning into calibration and test sets; ProteinGym contains no M1 variants. Single and paired attributes on the x-axis indicate the attributes used for calibration; "disorder" denotes ordered versus disordered residues, and "RaCoon" denotes the full pipeline. Error bars indicate ±1 SD across 100 calibration iterations, each evaluated using 100 non-parametric bootstrap test resamples. Classification thresholds were optimized per model using the Youden J-statistic. TP, true positive; TN, true negative; Acc, accuracy.

**Figure S14.** RaCoon performance on 46 balanced ClinVar proteins

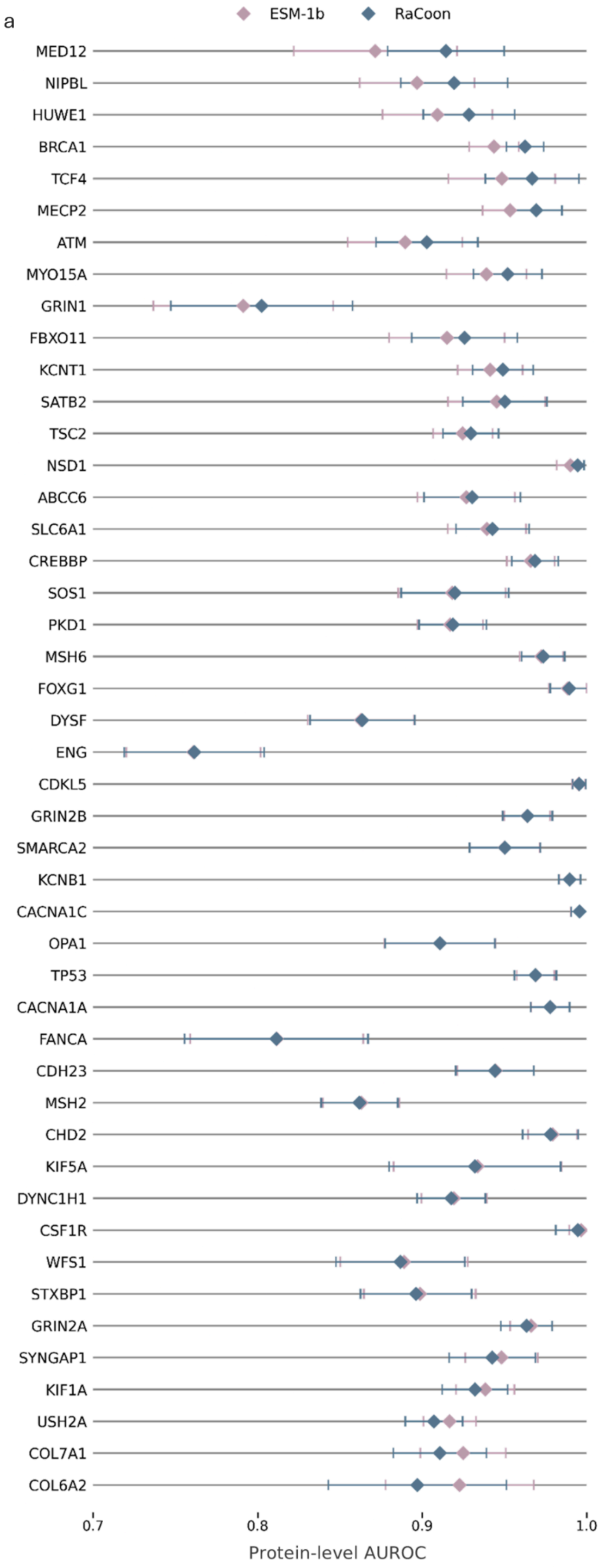
a
ESM-1b
RaCoon
MED12
NIPBL
HUWE1
BRCA1
TCF4
MECP2
ATM
MYO15A
GRIN1
FBXO11
KCNT1
SATB2
TSC2
NSD1
ABCC6
SLC6A1
CREBBP
SOS1
PKD1
MSH6
FOXG1
DYSF
ENG
CDKL5
GRIN2B
SMARCA2
KCNB1
CACNA1C
OPA1
TP53
CACNA1A
FANCA
CDH23
MSH2
CHD2
KIF5A
DYNC1H1
CSF1R
WFS1
STXBP1
GRIN2A
SYNGAP1
KIF1A
USH2A
COL7A1
COL6A2
0.7
0.8
0.9
1.0
Protein-level AUROC

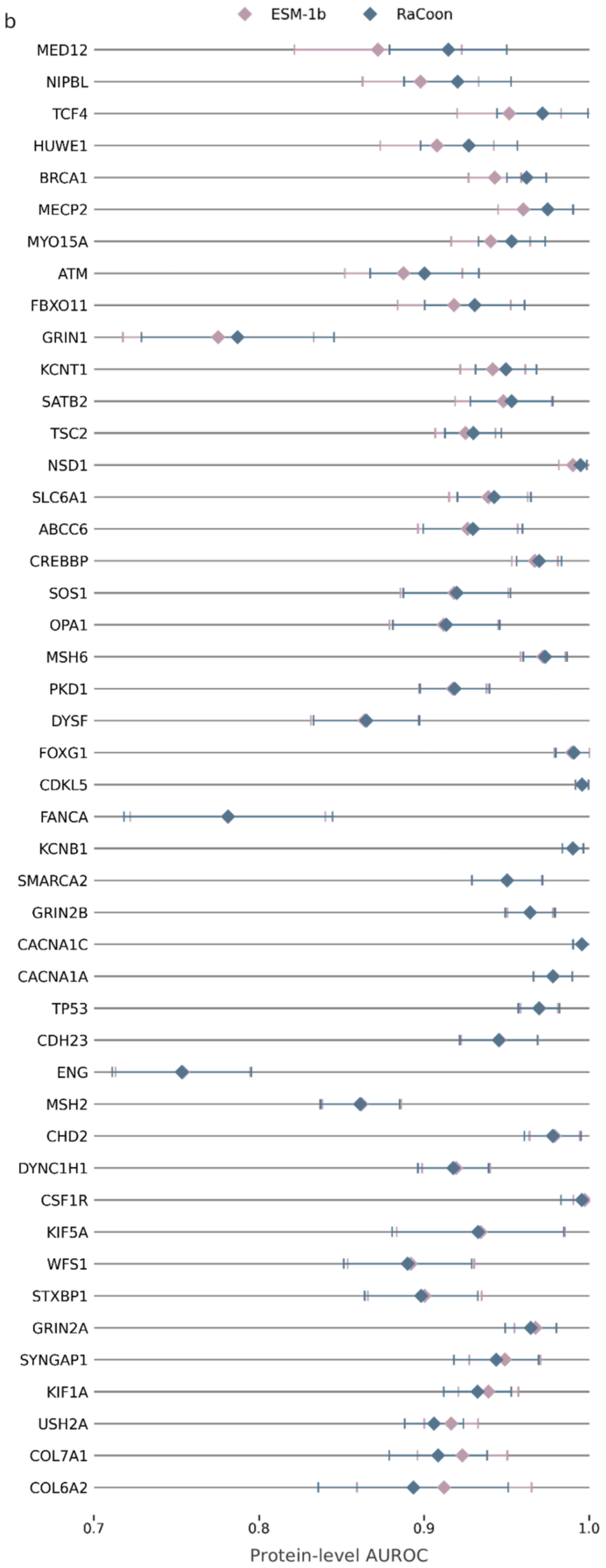

b
ESM-1b
RaCoon
MED12
NIPBL
TCF4
HUWE1
BRCA1
MECP2
MYO15A
ATM
FBXO11
GRIN1
KCNT1
SATB2
TSC2
NSD1
SLC6A1
ABCC6
CREBBP
SOS1
OPA1
MSH6
PKD1
DYSF
FOXG1
CDKL5
FANCA
KCNB1
SMARCA2
GRIN2B
CACNA1C
CACNA1A
TP53
CDH23
ENG
MSH2
CHD2
DYNC1H1
CSF1R
KIF5A
WFS1
STXBP1
GRIN2A
SYNGAP1
KIF1A
USH2A
COL7A1
COL6A2
0.7
0.8
0.9
1.0
Protein-level AUROC

**Figure S14. RaCoon performance on 46 balanced ClinVar proteins.** Proteins selected to have at least 100 annotations and a pathogenic-to-benign ratio between 30:70 and 70:30. Diamonds represent the mean AUROC per protein for RaCoon (blue) and ESM1b (pink). Error bars indicate variability across 100 calibration iterations on ClinVar_Balanced, each evaluated using 100 non-parametric bootstrap test samples. (**a**) M1 variants included (**b**) M1 variants excluded

**Figure S15.** RaCoon performance evaluation within-dataset

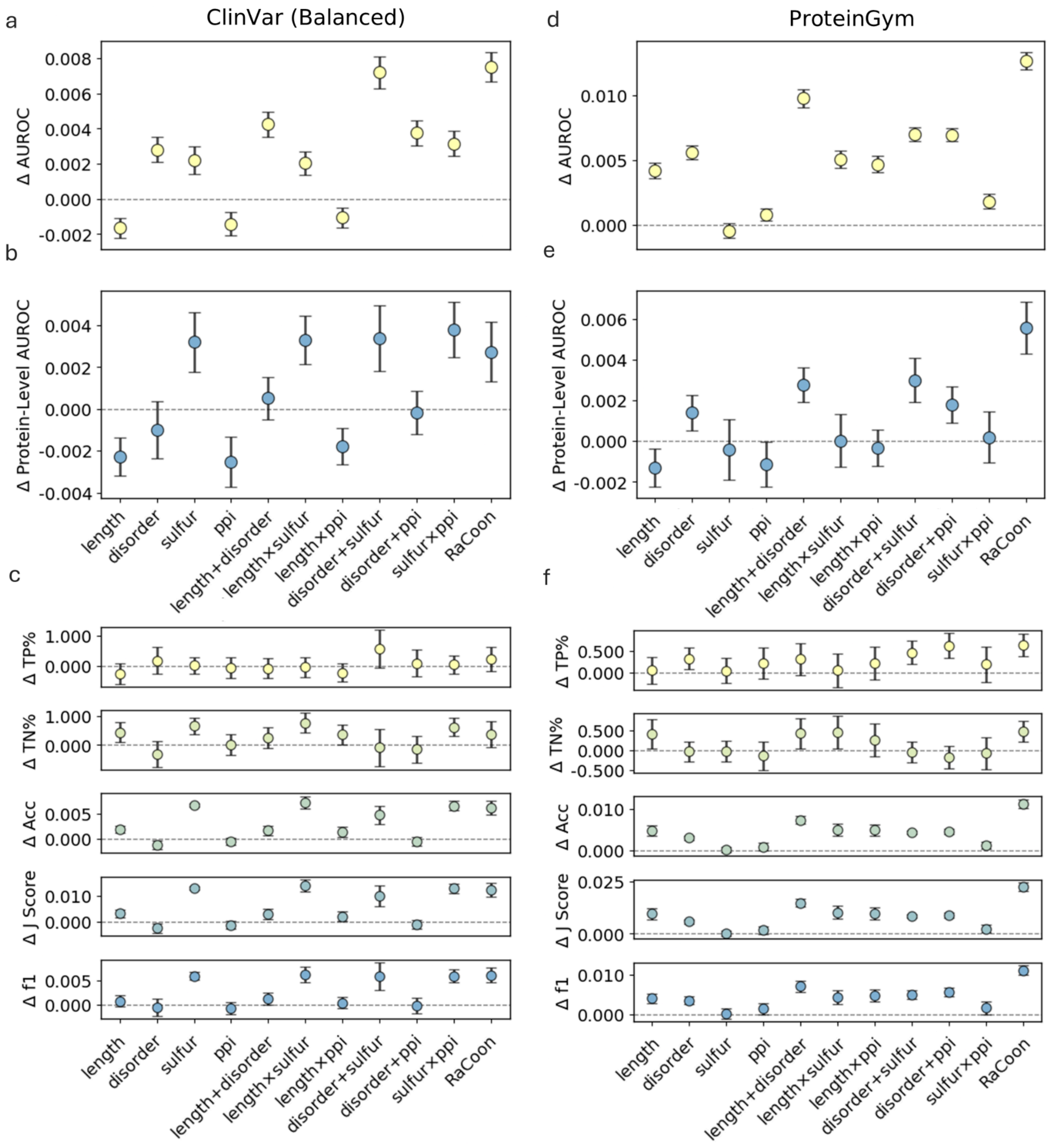

**Figure S15. RaCoon performance evaluation within-dataset.** Changes in overall performance relative to the uncalibrated ESM1b. Error bars indicate ± 1 SD across 100 randomized training iterations, each evaluated with 100 non-parametric bootstrap test resamples. Top: global AUROC; middle: per-protein AUROC; bottom: discrimination metrics. Per-protein AUROC computed for sequences containing ≥ 10 variants with at least one pathogenic and one benign label. Single and paired attributes on the x-axis indicate the attributes used for calibration; "disorder" denotes ordered versus disordered residues, and "RaCoon" denotes the full pipeline. TP = true positive; TN = true negative; thresholds were optimized per model using the Youden J-statistic. **a-c** Evaluation and calibration on ClinVar_Balanced **d-f** Evaluation and ProteinGym Clinical substitution Benchmark.

**Figure S16.** RaCoon performance across protein-level label ratio

a

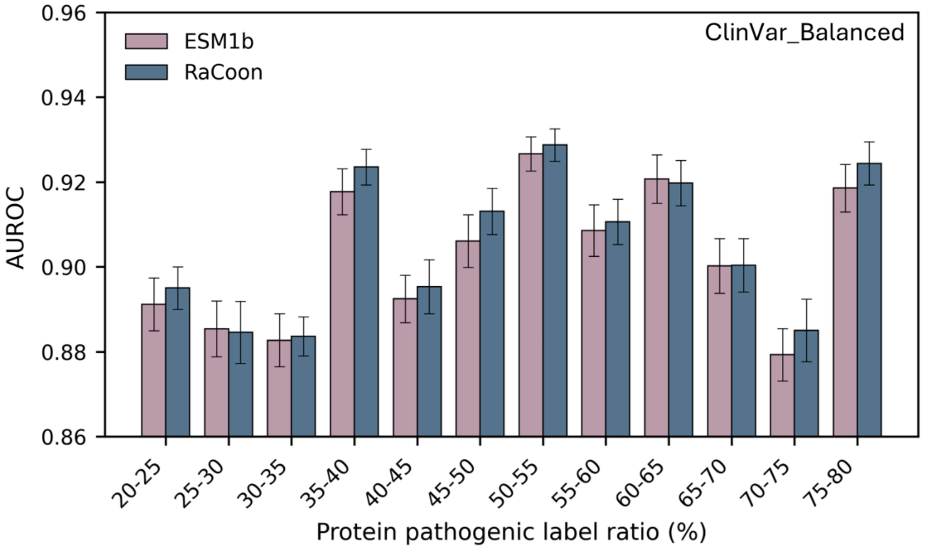

| bin | 20-25 | 25-30 | 30-35 | 35-40 | 40-45 | 45-50 | 50-55 | 55-60 | 60-65 | 65-70 | 70-75 | 75-80 |
|---|---|---|---|---|---|---|---|---|---|---|---|---|
| #prot | 146 | 164 | 162 | 94 | 109 | 52 | 136 | 89 | 94 | 99 | 85 | 107 |
| #mut | 4508 | 3350 | 3864 | 3998 | 3264 | 2529 | 3908 | 2768 | 2752 | 2814 | 2575 | 4144 |

b

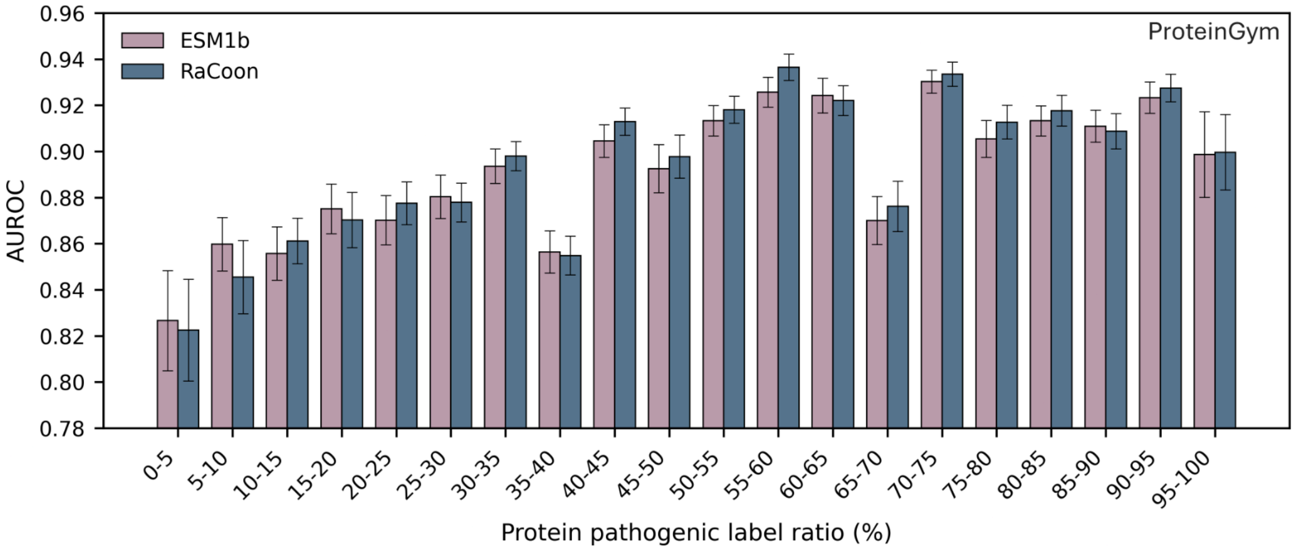

| bin | 0-5 | 5-10 | 10-15 | 15-20 | 20-25 | 25-30 | 30-35 | 35-40 | 40-45 | 45-50 | 50-55 | 55-60 | 60-65 | 65-70 | 70-75 | 75-80 | 80-85 | 85-90 | 90-95 | 95-100 |
|---|---|---|---|---|---|---|---|---|---|---|---|---|---|---|---|---|---|---|---|---|
| #prot | 81 | 179 | 242 | 159 | 172 | 198 | 224 | 85 | 129 | 42 | 219 | 75 | 81 | 124 | 77 | 90 | 115 | 95 | 93 | 45 |
| #mut | 4232 | 4203 | 4261 | 3186 | 2664 | 2359 | 3545 | 2501 | 2392 | 1169 | 2758 | 1883 | 1798 | 1988 | 2934 | 2330 | 3552 | 3998 | 6954 | 4020 |

**Figure S16. RaCoon performance across protein-level label ratio.** AUROC of ESM1b and RaCoon (y-axis) across protein-level pathogenic-fraction bins (x-axis) in (a) ClinVar_Balanced and (b) ProteinGym. Tables below each panel report the numbers of proteins and variants per bin. Table cells are shaded according to the better-performing model in each bin. Error bars indicate ±1 SD across 100 randomized within-dataset calibration-training iterations, each evaluated using 100 non-parametric bootstrap test resamples on variants not seen during calibration.

**Figure S17.** RaCoon ROC Curves

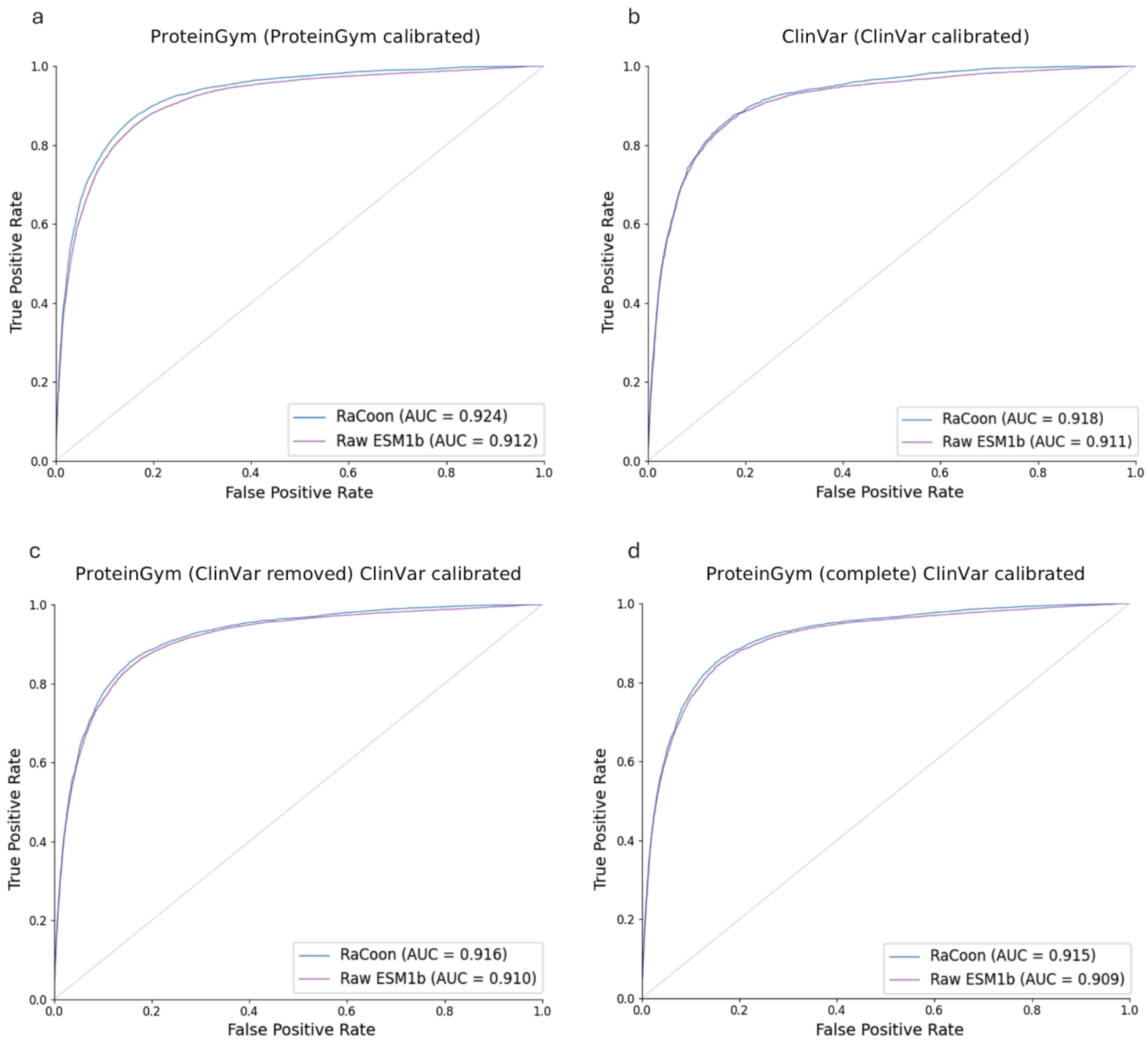

**Figure S17. ROC Curve Racoon vs ESM1b.** Performance is evaluated on test sets of variants not used during RaCoon calibration. **a-b** Within-dataset evaluation on ProteinGym (**a**) and ClinVar_Balanced (**b**). **c-d** Cross-dataset evaluation on ProteinGym using ClinVar_Balanced for calibration: (**c**) variants overlapping with ClinVar removed (even if not used for calibration), (**d**) evaluation on the full ProteinGym benchmark.
